# Optimal experimental designs for estimating genetic and non-genetic effects underlying infectious disease transmission

**DOI:** 10.1101/2022.01.10.475628

**Authors:** Christopher Pooley, Glenn Marion, Stephen Bishop, Andrea Doeschl-Wilson

## Abstract

**Background:** Infectious disease spread in populations is controlled by individuals’ susceptibility (propensity to acquire infection), infectivity (propensity to pass on infection to others) and recoverability (propensity to recover/die). Estimating the effects of genetic risk factors on these host epidemiological traits can help reduce disease spread through genetic control strategies. However, the effects of previously identified ‘disease resistance SNPs’ on these epidemiological traits are usually unknown. Recent advances in computational statistics make it now possible to estimate the effects of single nucleotide polymorphisms (SNPs) on these traits from longitudinal epidemic data (*e*.*g*. infection and/or recovery times of individuals or diagnostic test results). However, little is known how to optimally design disease transmission experiments or field studies to maximise the precision at which pleiotropic SNP effects estimates for susceptibility, infectivity and recoverability can be estimated.

**Results:** We develop and validate analytical expressions for the precision of SNP effects estimates on the three host traits assuming a disease transmission experiment with one or more non-interacting contact groups. Maximising these leads to three distinct ‘experimental’ designs, each specifying a different set of ideal SNP genotype compositions across groups: a) appropriate for a single contact-group, b) a multi-group design termed “pure”, and c) a multi-group design termed “mixed”, where ‘pure’ and ‘mixed’ refer to contact groups consisting of individuals with the same or different SNP genotypes, respectively. Precision estimates for susceptibility and recoverability were found to be less sensitive to the experimental design than infectivity. Data from multiple groups were found more informative about infectivity effects than from a single group containing the same number of individuals. Whilst the analytical expressions suggest that the multi-group pure and mixed designs estimate SNP effects with similar precision, the mixed design is preferable because it uses information from naturally occurring infections rather than those artificially induced. The same optimal design principles apply to estimating other categorical fixed effects, such as vaccinations status, helping to more effectively quantify their epidemiological impact.

An online [software tool](https://theiteam.github.io/SIRE-PC.html) *SIRE-PC* has been developed which calculates the precision of estimated substitution and dominance effects of a single SNP (or vaccine status) associated with all three traits depending on experimental design parameters.

**Conclusions:** The developed methodology and software tool can be used to aid the design of disease transmission experiments for estimating the effect of individual SNPs and other categorical variables underlying host susceptibility, infectivity and recoverability.

## Background

Infectious diseases constitute one of the biggest threats to sustainable livestock and aquaculture production, global food security and human health. Over the last decades, genome wide association studies (GWAS) together with high density sequencing and other - omics have facilitated enormous breakthroughs in disease genetics, with the number of identified genetic loci associated with disease resistance increasing at a rapid rate [1-6]. Accordingly, expectations for reducing infectious disease prevalence through genomic selection for disease resistance are rising, and some real-world applications have already demonstrated that these expectations can be met in practice [7].

The most effective way to reduce infectious disease prevalence in a population is to reduce individual’s susceptibility to infection or their ability to transmit infections, once infected. Yet, remarkably little is known about the role of previously identified ‘resistance’ loci in infectious disease transmission. This is because in most studies, disease resistance refers to the resistance of an infected animal to develop disease or other side-effects from infection (*e*.*g*. performance reduction or death), rather than to resistance to becoming infected or transmitting the infection [8-10]. Hence it is not known whether selection for disease resistance actually reduces disease prevalence, as animals that carry the beneficial resistance alleles may still become infected and transmit the infection. Furthermore, discovery of SNPs associated with disease resistance often originate from large scale disease challenge experiments, in which individuals are artificially infected or exposed to a specific pathogen strain and dose and their response to infection is measured [11-13]. Estimating the effect of genetic loci identified from these studies on disease transmission would, however, require field or experimental epidemic data where the infection is transmitted naturally between individuals.

Epidemiological models are widely used to identify risk factors for disease transmission in populations and to assess the impact of control measures on these. Particularly relevant for genetically heterogeneous populations are compartmental models in which individuals are classified as, for example, susceptible to infection (S), infected and infectious (I), or recovered/removed (dead) (R) [14]. These epidemiological SIR models point naturally to three distinct host genetic traits that characterise the key processes of disease transmission dynamics within a population, *i*.*e*. individual *susceptibility, infectivity* and *recoverability* [15-17]. In the epidemiological context, *susceptibility* is defined as the relative risk of an uninfected individual becoming infected when exposed to a typical infectious individual or infectious material excreted from such an individual, *infectivity* is the propensity of an individual, once infected, to transmit infection to a typical (average) susceptible individual, and *recoverability* is the propensity of an individual, once infected, to recover or die [15, 18, 19]. For SIR models recoverability is the inverse of the duration of the infectious period.

Conceptually, genetic improvement in any or all of these underlying epidemiological host traits reduces disease spread within and across populations. Indeed, recent advances in treating disease as an indirect genetic effect (IGE) have pointed to far great response to selection than had been previously expected [20, 21]. This has been demonstrated for infectious pancreatic necrosis (IPN), a viral disease inflicting large mortalities in Atlantic salmon populations. Previous GWAS had identified a single quantitative trait locus (QTL) that explains over 80% of the genetic variation in mortality caused by IPN [22, 23]. The corresponding candidate gene identified in subsequent fine-mapping studies was found to control primarily IPN virus internalization, *i*.*e*. host susceptibility [24]. A small-scale IPN transmission experiment, in which fish, after accounting for their relatedness structure, were assigned into different epidemic groups according to their IPN resistance genotypes, provided further evidence that the beneficial allele reduced the infectivity of IPN infected fish in addition to their susceptibility, and may also have favourable effects on the duration of the infectious period (*i*.*e*. their recoverability) [25]. This beneficial pleiotropic effect on all three epidemiological host traits may explain the rapid success of IPN-resistance breeding schemes that have led to the observed drastic reduction in IPN prevalence and associated mortalities within just a few generations of selection [26]. Incorporating these effects into epidemiological models can also inform management strategies on how to effectively prevent disease outbreaks in genetically heterogeneous populations [25].

Compared to conventional disease resistance traits used in most GWAS (*e*.*g*. individuals’ infection, disease or survival status, measure of pathogen load or immune or performance measures after infection challenge), the epidemiological host traits have the clear advantage that their role in disease spread is fully specified by the epidemiological models. However, until recently, estimation of genetic effects for these traits has proven challenging as they need to be inferred from observable disease phenotypes.

Novel advances in computational statistics, however, make it now possible to estimate genetic effects for these epidemiological host traits from longitudinal disease records of individuals [15, 27-30]. However, little attention has been paid to how disease transmission experiments should actually be designed to obtain accurate estimates. For example, previous studies have indicated that accurate estimation of genetic infectivity effects requires genetically related individuals distributed across different contact groups [15, 27, 28] and that relatedness among group mates can substantially affect precision and bias of the trait estimates [28, 31]. Hence, the appropriate number and size of contact groups, the genetic group composition, and other parameters that influence precision of estimates, need to be established.

This study has the following aims: Firstly, to derive analytical expressions for the precision of estimates of the effects of a particular SNP on the three underlying epidemiological host traits. For tractability these expressions assume a best case scenario (in which infection and recovery events are exactly known and other, potentially confounding factors, are ignored), and so they represent upper bounds on model parameter precision from real data. Secondly, to use these insights to develop optimal designs of disease transmission experiments aimed at estimating the effects of a single SNP of interest on host susceptibility, infectivity and recoverability. Thirdly, to validate the analytical expressions and designs for a range of realistic data scenarios (*e*.*g*. the inclusion of group effects, other fixed effects, residual noise and cases when only the deaths of individuals are recorded and infection times are unknown). Lastly, to present an easy-to-use online software tool to assist any potential experimenter in constructing a suitable design for their disease transmission experiment.

Although this study focuses on the estimation of SNP effects underlying disease transmission, it should be noted that the methodology and optimal design principles obtained also apply to investigating any other categorical variable (such as vaccination) on host susceptibility, infectivity and recoverability. Finally, the discussion section provides information regarding the application and extension of the methods and results presented here for evaluating the role of a single SNP in disease transmission to identifying loci associated with disease transmission in a GWAS and application to field data.

## Methods

### Key concepts, assumptions, terminology and data

To introduce the terminology and assumptions made in this study, Figure 1 illustrates the key features of a typical disease transmission experiment in farmed animals for the large number of infections that are primarily transmitted by direct contact. The experiment typically consists of one or more “contact groups”, where contacts are assumed to occur randomly within a group and no contact (and hence no transmission) occurs between groups.

**Figure 1:**
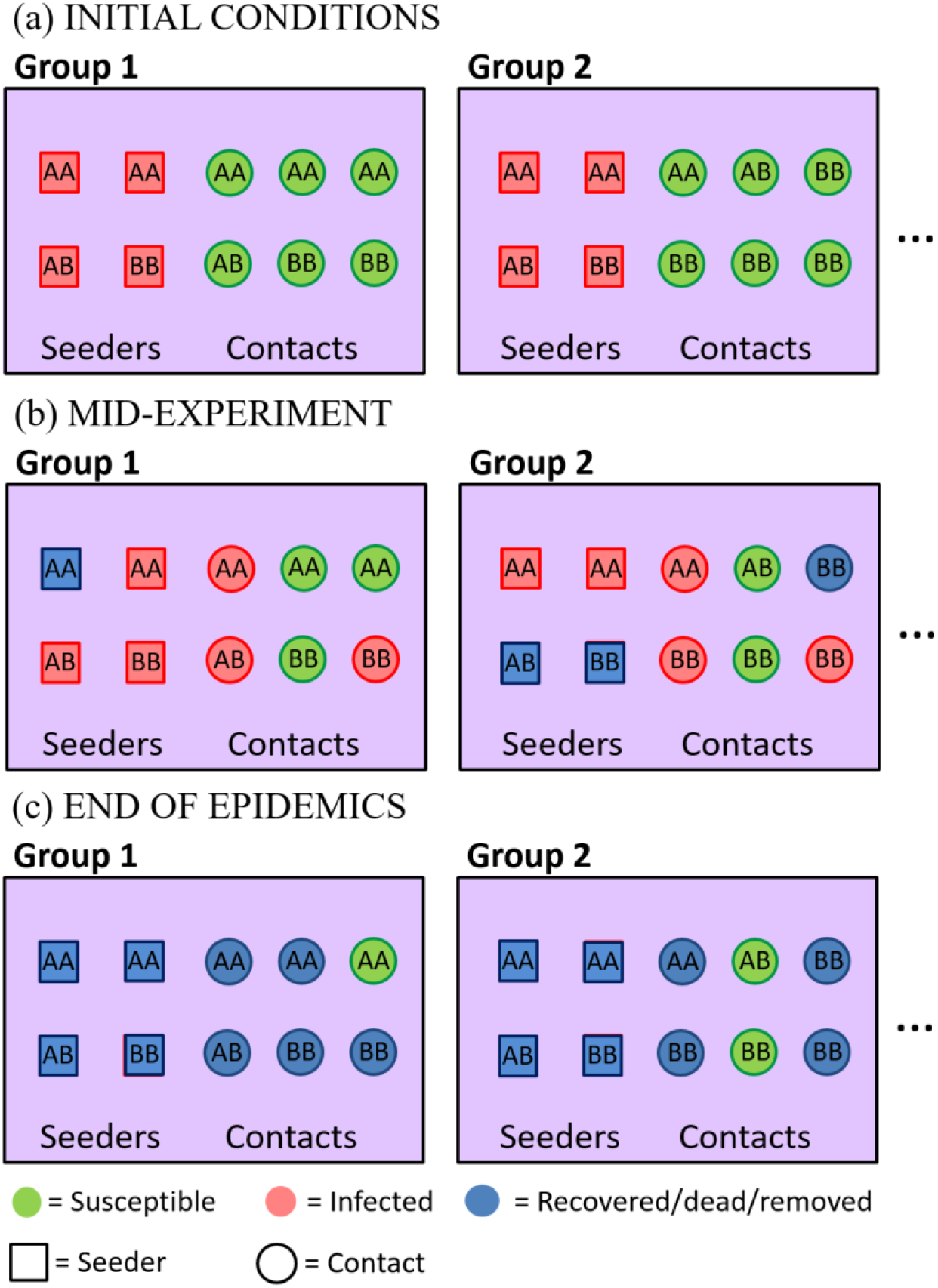
Schematic diagram of a disease transmission experiment. (a) The experiment consists of several contact groups in which some individuals are initially infected “seeders” and some are initially susceptible “contacts”. Each symbols represents an individual, and the annotations *AA, AB* and *BB* refer to the genotype of that individual at a given bi-allelic SNP under investigation. (b) As the experiment progresses some susceptible individuals become infected and some infected individuals recover. (c) If the experiment continues until the epidemics die out, only susceptible and recovered individuals are observed in the final state (note, for practical reasons experiments are usually terminated before this point). Note, the spatial separation of seeders (left) and contacts (right) in this diagram is for illustrative purposes only (random mixing between individuals is assumed).

In this study, which focuses on the estimation of the effects of a particular SNP, it is thus assumed that individuals are randomly distributed across contact groups with regards to the genetic effects on the epidemiological traits not captured by the SNP under consideration (see discussion). This implies, for example, that related individuals (*e*.*g*. full-sibs or half-sibs) are assumed to be equally distributed across contact groups. The overall population is assumed to be composed of diploid individuals with a bi-allelic genetic structure such that “*A*” and “*B*” represent different alleles at a particular SNP or genetic locus under investigation, resulting in three potential genotypes *AA,AB,BB*} (Fig. 1).

The transmission experiment starts with two types of individuals (Fig. 1a): “seeders” that are infected (either artificially or from prior exposure to other infected individuals^1^) at the beginning of the experiment and “contacts” that are susceptible to the disease. After some time (Fig. 1b), the infection has passed from the seeders to some of the contacts, and possibly also between contacts, whilst some infected individuals may have recovered or died (removed) from disease. Eventually (Fig. 1c), all infected individuals have recovered and typically some susceptible individuals remain that did not succumb to infection. In reality, however, the experiment may be terminated before all epidemics have finished and censoring of data will need to be accounted for in the analysis. In this study it is assumed that transmission dynamics are the same for seeder to contact individuals as for contact to contact individuals.

Genotypic variation across contact groups is characterised by the following key quantities defined for each group: *H*_seed_ and *H*_cont_ give the proportion of homozygotes (*i*.*e. AA* or *BB*) in the seeders and contacts, respectively, and χ_seed_ and χ_cont_ give the so-called “homozygote balance”, defined as the proportion of *AA* minus the proportion of *BB* individuals^2^. These quantities are used in later analysis and, along with other key parameters (described below), are summarised in Table 1.

**Table 1.**
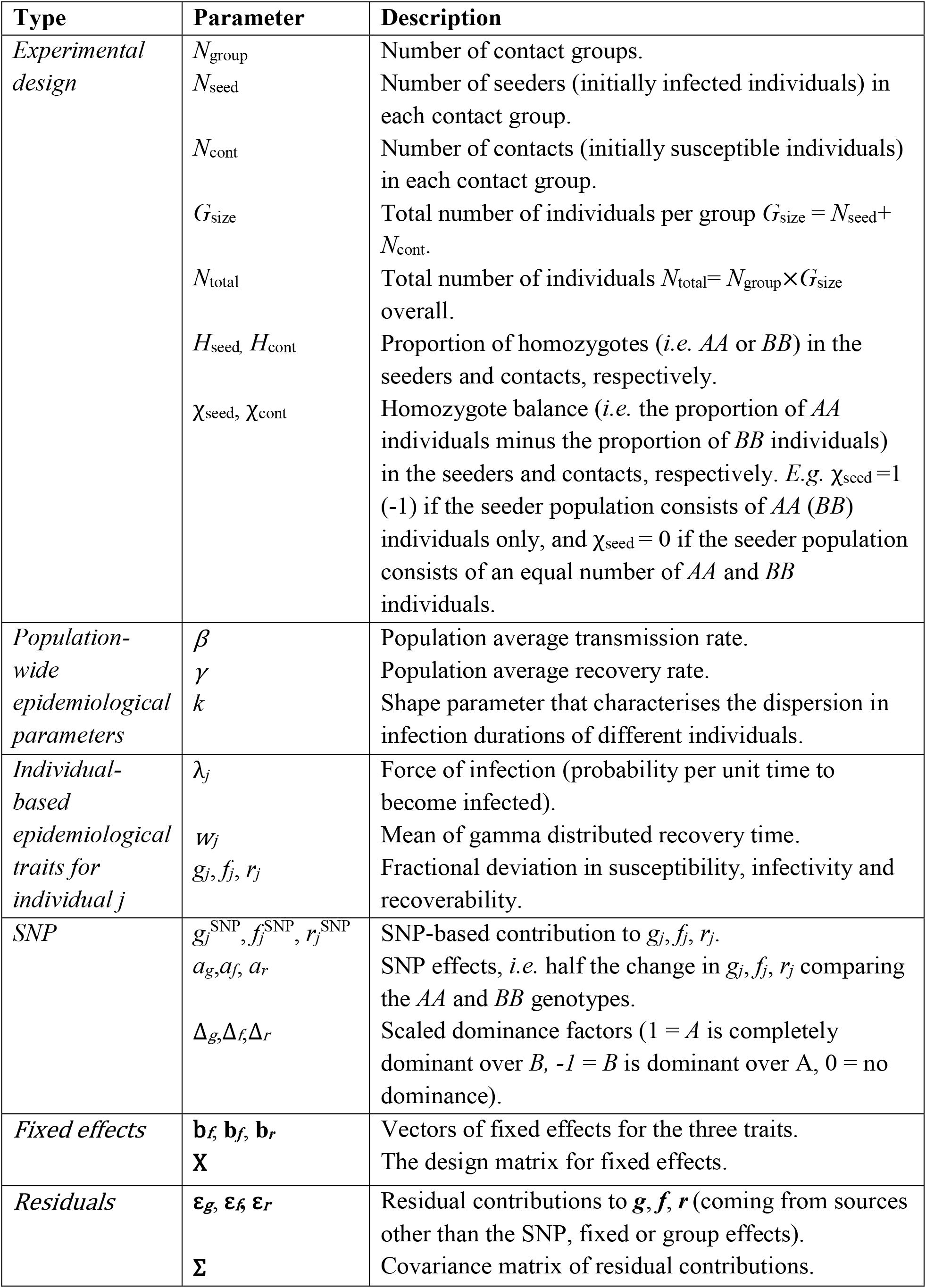
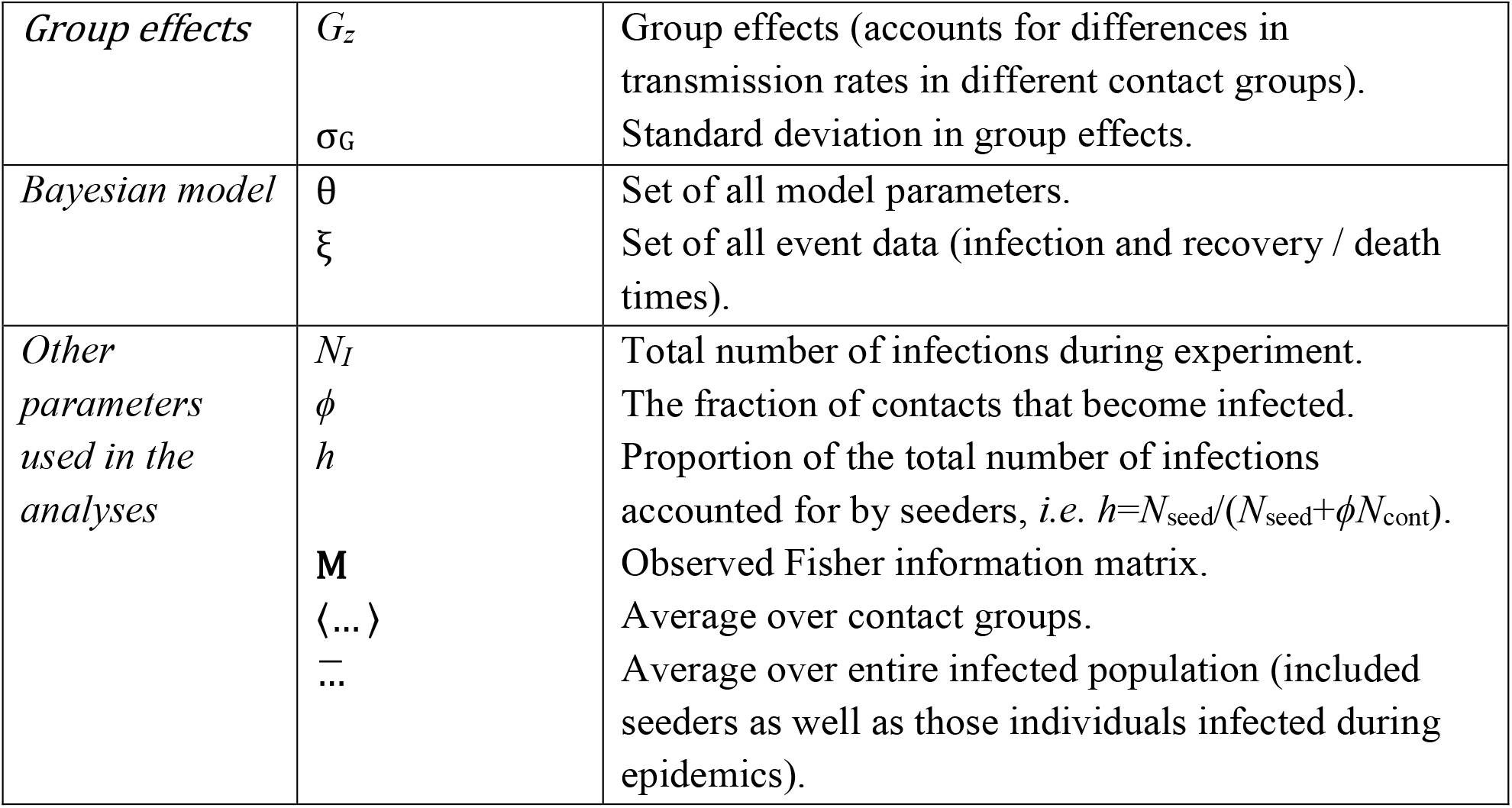
List of key parameters and quantities.

Previous studies have shown that SNP and other genetic effects for the epidemiological parameters can be inferred from a wide range of available data that can be routinely collected from disease transmission experiments [15, 25, 29, 30]. These may consist of the times at which individuals become infected and/or recover/die, or alternatively results from disease diagnostics tests which furnish information on the disease status of individuals at particular points in time. Note, estimates can be inferred even for censored data and it is not required that transmission routes (who infects who) are known [15].

Based on these concepts, for a given disease and epidemiological data from a fixed number of genotyped animals and contact groups, the optimal experimental design is determined by the relative number of seeders *N*_seed_, contacts *N*_cont_, proportion of homozygotes *H*_seed_ and *H*_cont_, and homozygote balance χ_seed_ and χ_cont_ across groups that maximise the precision with which SNP effects on susceptibility, infectivity and recoverability can be estimated from the data. In particular, we identify optimal designs for basic blocks of groups (either 1, 4 or 9) which can be replicated one or several times to make up the total contact group number *N*_group_ and associated number of seeder and contact individuals of different genotypes in the experiment.

### The genetic-epidemiological model

The infection dynamics within each contact group described above (and illustrated in Fig. 1) can be represented by an epidemiological SIR model, with individuals classified as being either susceptible to infection (S), infected and infectious (I), or recovered/removed/dead (R) [14]. The incorporation of individual-based trait variation into this model is taken from [15], which we briefly reiterate here for completeness. The force of infection *λ*_*j*_ (*i*.*e*. the probability per unit time individual *j* becomes infected) and mean infection duration *w*_*j*_ are given by

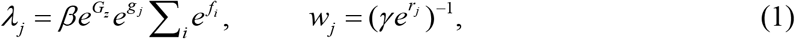

where *β* is a population average transmission rate, *γ* is a population average recovery rate, *g*_*j*_ and *r*_*j*_ represent fractional deviations^3^ in the susceptibility and recoverability of individual *j*, and *f*_*i*_ represents the fractional deviation in infectivity of individual *i* (the sum goes over all currently infected individuals sharing the same contact group as *j*). Finally, *G*_*z*_ is a random effect (for group *z* with mean zero and standard deviation σ_G_) that accounts for group-specific factors that influence the overall speed of an epidemic in one contact group relative to another (*e*.*g*. animals kept in different management conditions or environmental differences).

The time for individual *j* to recover after being infected is taken to be gamma distributed with mean *w*_*j*_ and shape parameter *k* [15].

The individual-based fractional deviations in susceptibility, infectivity and recoverability are parameterised in the following way:

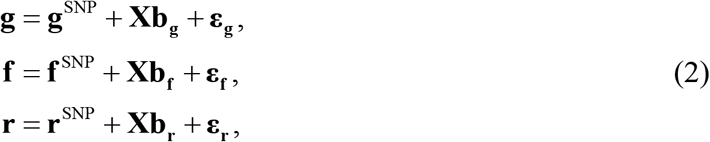

where **g, f** and **r** are vectors (with elements for each of the individuals) that are decomposed into **g**^SNP^, **f**^SNP^ and **r**^SNP^, which gives the contribution from the SNP under investigation, fixed effects **b**_***g***_, **b**_***f***_ and **b**_***r***_, where **X** is a design matrix (*e*.*g*. to account for sex differences in the traits or vaccination status), and residuals **ε**=(**ε**_**g**_,**ε**_**f**_,**ε**_**r**_) that account for contributions from all SNPs other than the one being investigated, and other sources of polygenic and non-genetic variation. Residuals are multivariate-normal distributed with zero mean and covariance matrix **I×Σ**, where **I** is the identity matrix reflecting no correlation between individuals and **Σ** is a 3×3 covariance matrix that characterises potential correlations between traits^4^.

The residual structure in model Eq.(2) does not explicitly distinguish between random genetic and environmental effects, and relies on the assumption that individuals are distributed with regards to the genetic effects on the epidemiological traits not captured by the SNP under consideration (see discussion for relaxing this assumption).

The SNP effects for individual *j* are dependent on *j*’s genotypes in the following way:

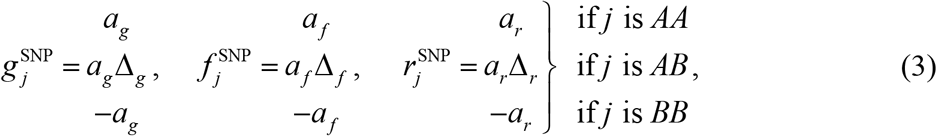

where *a*_*g*_, *a*_*f*_, and *a*_*r*_ give half the difference in traits between the *AA* and *BB* homozygote genotypes and Δ_*g*_, Δ_*f*_ and Δ_*r*_ represent the degree of dominance (a value of 1 corresponds to complete dominance of the *A* allele over the *B* allele and -1 when the reverse is true, whereas absence of dominance is represented by a value of 0) [32].

The model above contains numerous parameters, but from the point of view of establishing SNP-based associations the key quantities are *a*_*g*_, *a*_*f*_, and *a*_*r*_, subsequently referred to as the “SNP effects”, which characterise the changes in susceptibility, infectivity and recoverability conferred by different SNP genotypes (note, Δ_*g*_, Δ_*f*_ and Δ_*r*_ are also important if dominance is of particular interest, as discussed later).

## Results

We first derived analytical expressions for the precisions of estimates of SNP and dominance effects for the three epidemiological traits (see below) which provided first insights for how these are affected by the experimental design (here precision is determined by the expected standard deviations (SD) of the SNP / dominance effect estimates). We then investigate maximising these precisions through optimal design.

For validation, analytical SDs were compared against inferred values (using the software tool SIRE [15]) directly estimated from simulated epidemic and genetic data resulting from different experimental setups (see Additional file 1 for details).

### Analytical expressions

#### Susceptibility and infectivity SNP effects

Based on the model presented in the previous section, it is possible to analytically approximate the marginalised likelihood for the two variables *a*_*g*_, *a*_*f*_ as a two dimensional multivariate-normal distribution with inverse covariance matrix given by the observed 2×2 Fisher information matrix (see Additional file 2 for a derivation of this expression):

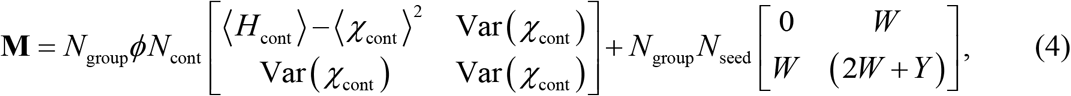

where

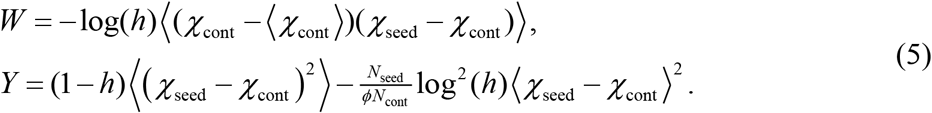

The parameters are defined as follows: *N*_group_ is the number of groups, *ϕ* represents the average fraction of contacts that become infected^5^, *h*=*N*_seed_/(*N*_seed_+*ϕN*_cont_) is the proportion of infected individual that are seeders, *H*_cont_ gives the proportion of homozygotes contacts (*i*.*e*. proportion of *AA* plus *BB*), and, finally, χ_seed_ and χ_cont_ give the homozygote balance (*i*.*e*. proportion of *AA* minus *BB*) in the seeders and contacts, respectively. Note that *H*_cont_, χ_seed_ and χ_cont_ are all group-dependent design parameters. The angle brackets in Eqs.(4) and (5) denote averaging these quantities across groups, and Var(χ_cont_) gives the variance across groups in the homozygote balance for the contacts.

An important point to take from Eq.(4) is that **M** is actually the sum of two matrices: The first corresponds to information provided by infections that occur during the course of the observed epidemics (note this contribution contains a factor *N*_group_*ϕN*_cont_, which is the total expected number of within-group infections) and the second comes from information gained as a result of differences in genetic makeup between seeders and contacts in the initial conditions^6^ (this contains a factor giving the total number of initially infected individuals *N*_seed_).

Inversion of the observed Fisher information matrix defined in Eq.(4) leads to estimates for the standard deviations in *a*_*g*_ (see Additional file 3 for further details)

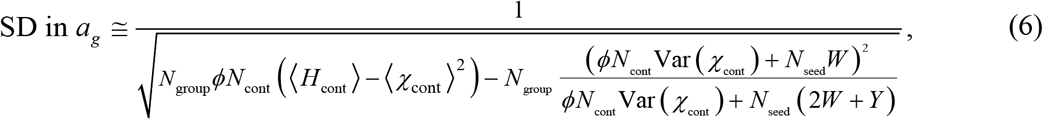

and *a*_*f*_

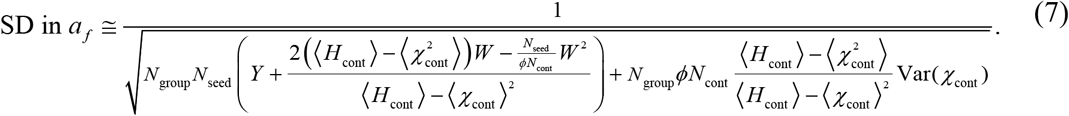

These rather unwieldy expressions reflect a complex confounding between estimating susceptibility and infectivity SNP effects. They show that parameters precisions are dependent not only on the number of seeders and contacts, but also on the genetic compositions in these populations across groups.

Despite their apparent complexity, a number of important design lessons can be drawn from Eqs.(6) and (7): Firstly, they both scale as 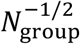 (which means that increasing the number of groups by a factor of four halves the SDs). Secondly, a higher proportion of infections *ϕ* implies greater precision (the more contacts that become infected, the greater the available information on which inferences can be based). Thirdly, for large *N*_cont_ and fixed *N*_seed_ we observe that both SDs scale as 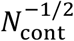 meaning that greater precision results from a larger contact population (a notable exception to this is the case of a single contact group, for which the variance Var(χ_cont_) in Eq.(7) becomes exactly zero, and so this term vanishes). Lastly, precision is maximised when the homozygosity in the contacts is one, *i*.*e*. ⟨*H*_cont_⟩=1, referring to experiments that only contains *AA* and *BB* contact individuals (as *AB* heterozygotes provide less information because they dilute the relative effects of the *A* and *B* alleles in the case of zero dominance investigated here).

#### Recoverability SNP effects

Equivalent analytical expressions for recoverability can be derived (see Additional file 4 for further details). Assuming no dominance, this leads to

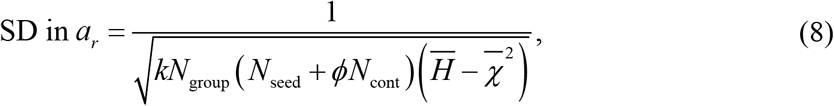

where

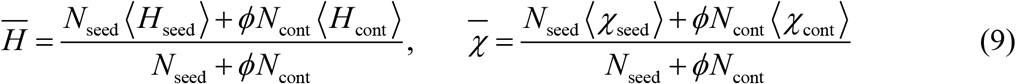

represent the average homozygosity and homozygote balance for the entire infected population (*i*.*e*. including the seeders as well as those contacts infected during the experiment), respectively.

### Dominance

Earlier we made the assumption of no dominance between *A* and *B* alleles. However it’s worth noting that the analytical results above can still be used in the case of complete dominance by means of a simple change in parameter definitions. In the case in which allele *A* has complete dominance over *B*, the genotypes *AA* and *AB* become indistinguishable, and so the homozygote balance parameters χ_seed_ and χ_count_ can be redefined as the proportion of *AA and AB* individuals minus the proportion of *BB* individuals in the seeders and contacts, and the homozygosity *H*_cont_ gets set to exactly one.

In the general case, expressions for the SDs in Δ_*g*_, Δ_*f*_ and Δ_*r*_ are given as follows (see Additional file 5 for details):

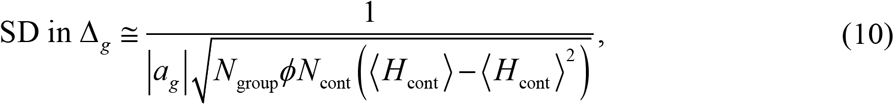

where ⟨*H*_cont_⟩ is the average homozygosity of all contact individuals. Interestingly, this expression is optimised when ⟨*H*_cont_⟩ =½, irrespective of exactly how the homozygote individuals are distributed across groups. Note, the expression in Eq.(10) diverges in the limit of no homozygosity (⟨*H*_cont_⟩=0) or complete homozygosity (⟨*H*_cont_⟩=1), as would be expected.

In the case of infectivity

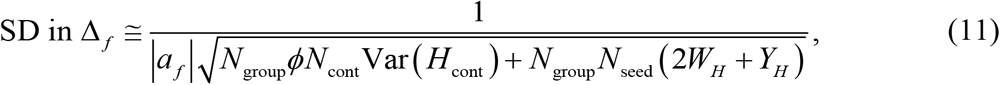

where

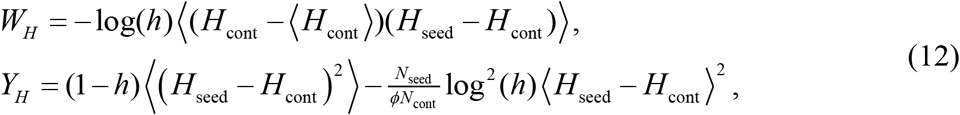

and recoverability

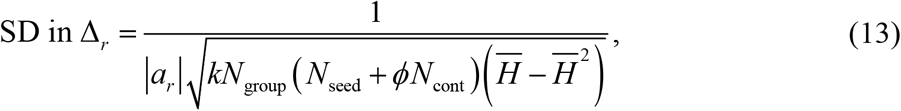

where 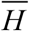 is the average homozygosity over the entire infected population (regardless of their distribution across groups), as defined in Eq.(9).

### Experimental designs

From the outset it should be emphasised that there is no single experimental design optimum. This is for two reasons: Firstly, whilst one particular design might estimate a given parameter as precisely as possible, other parameters in the model might be much less certain (so to some extent optimal design will depend on a trade-off in precision between different parameter estimates). Secondly, practical considerations often restrict what can be implemented in reality (*e*.*g*. physical or budget constraints may restrict the number of groups or group sizes^7^).

As will be demonstrated later, the infectivity parameter *a*_*f*_ is the most difficult of the SNP effects to estimate. It is natural, therefore, to focus on experimental designs that reduce this as much as possible. In the case of a single contact group, mathematical minimisation of Eq.(7) can explicitly be performed leading to a unique optimal solution (see below). In the case of multiple groups, however, such minimisation is challenging due to the complexity of the expression. Nevertheless, it was found that individually maximising each of the two terms in the denominator in Eq.(7) led to two contrasting approaches^8^.

Consequently, three basic designs for disease transmission experiments emerge: One for a single contact group, one referred to as the “pure” design and one referred to as the “mixed” design. These designs are illustrated in Fig. 2 and are discussed in detail below (along with design-specific analytical expressions for the SD in *a*_*f*_).

**Figure 2:**
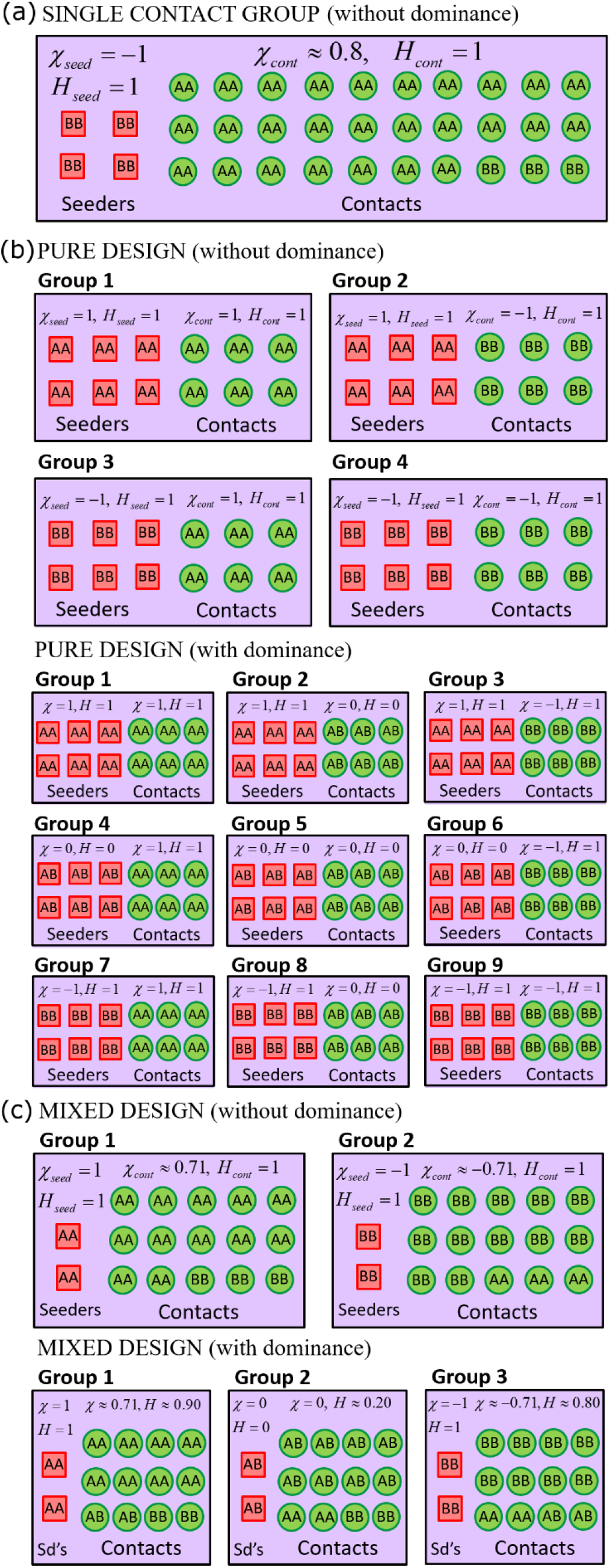
Optimal experimental designs. This figure shows the optimal composition of the seeder and contact populations for different experimental designs: (a) Single contact group design: ∼15% of individuals are seeders, where seeders have genotype *BB* (or *AA*) and contacts predominately have genotype *AA* (or *BB*), with ∼10% *BB*, to allow for estimation of the susceptibility SNP effect *a*_*g*_. Estimation of dominance was found to be challenging using only a single contact group (not shown). (b) Multiple groups “pure” design: ∼47% of individuals are seeders. Seeders and contacts consist of different combinations of *AA* and *BB* across groups (and *AB* when dominance is investigated). (c) Multiple groups “mixed” design: a small number of individuals are seeders (typically two or three, sufficient to initiate epidemics). When dominance is not investigated there is a 83%/17% split in *AA*/*BB* individuals in the contact population in group 1 and *vice-versa* in group 2. When dominance is investigated there is a 80%/10%/10% split in *AA*/*AB*/*BB* individuals in the contact population in group 1, and these proportions are permuted to define the two other groups. Optimisation of these designs was (for the most part) based on maximising the precision with which the infectivity SNP effect *a*_*f*_ can be estimated (since this was generally the most difficult trait to estimate). However in cases where maximal precision for *a*_*f*_ corresponds to minimal precision for *a*_*g*_, values are chosen to give equal precision to the two (*e*.*g*. ∼10% *BB* in (a), as discuss in the paper). The percentages above are, to a large extent, independent of *R*_*0*_ (see Additional file 8) or other factors in the model/data (see Additional file 11). For reference the homozygote balance χ_seed_, χ_cont_ (*i*.*e*. proportion of *AA minus BB* individuals) and homozygosity *H*_seed_ and *H*_cont_ (*i*.*e*. proportion of *AA plus BB* individuals) are shown for each design. The same basic designs can be replicated multiple times within an experiment. Note that the results equally apply to the estimation of non-genetic *e*.*g*. vaccination effects (*AA* replaced with “Vac.” and *BB* replaced with “Unvac.” and dominance not applicable). Also, the spatial separation between seeders and contacts in this diagram is for illustrative purposes only.

Note, typically to increase experimental power it is routine to perform multiple replicates of a given experimental design. This possibility is incorporated into the analytical expressions below by virtue of the fact that *N*_group_ refers to the total number of contact groups across all replicates.

For simplicity the subsequent analytical expressions assume the basic reproduction number *R*_*0*_ is reasonably high such that most contacts become infected (*i*.*e. ϕ* ≈ 1).

### Single contact group

Here we consider the case in which the disease transmission experiment consists of just a single contact group, as illustrated in Fig. 2a. We investigate how the fraction of individuals that are seeders (*i*.*e. N*_seed_*/G*_size_) along with the genetic makeup in the seeders and contacts should be chosen to infer the values for the SNP effects as precisely as possible. This is undertaken by varying each of these quantities in turn whilst keeping the other two fixed. The results are shown in Fig. 3.

**Figure 3:**
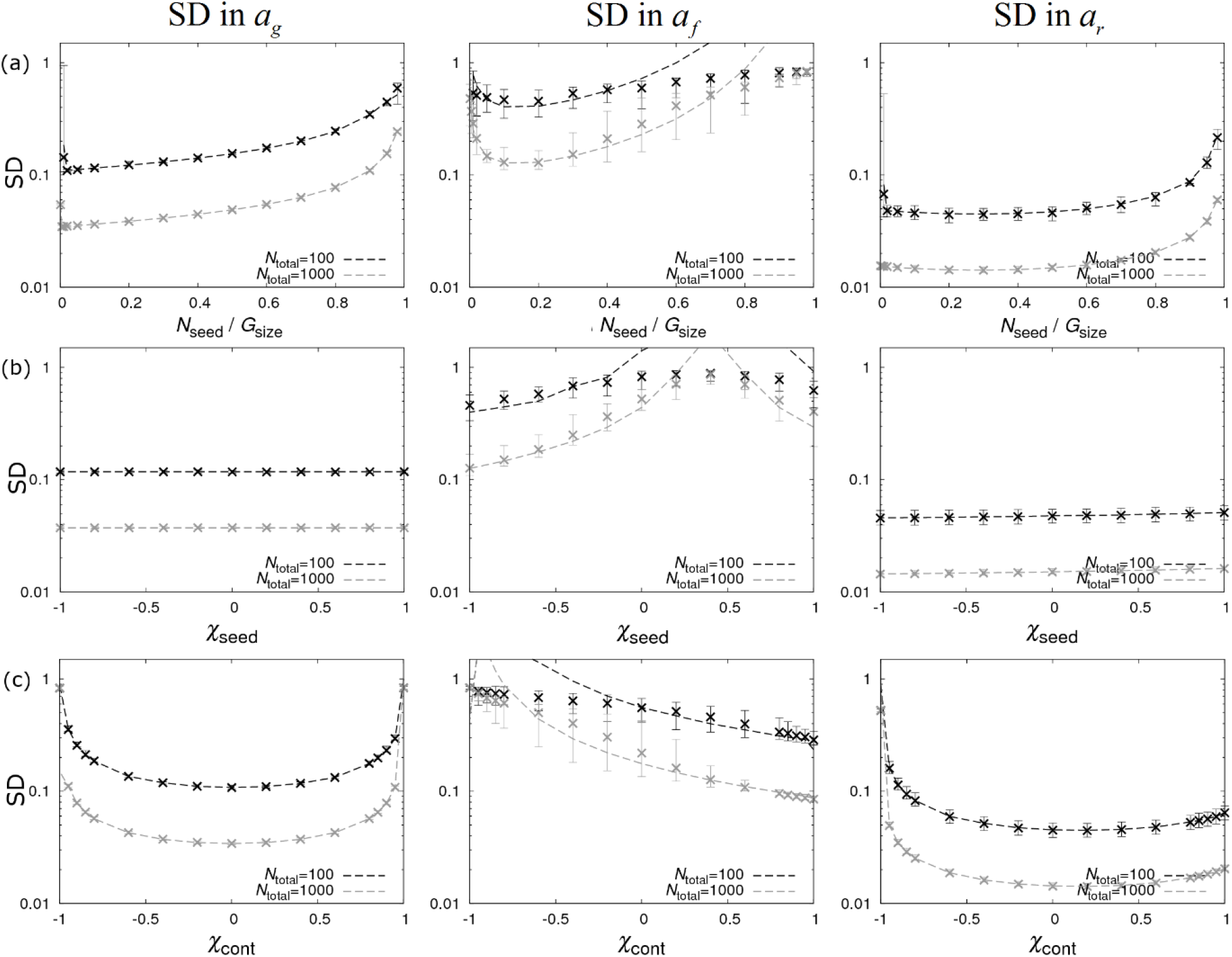
Single contact group design. Precision estimates for the single contact group design (without dominance) in Fig. 2a. The left, middle and right columns show graphs for standard deviations (SDs) in the SNP effects for susceptibility *a*_*g*_, infectivity *a*_*f*_ and recoverability *a*_*r*_ under different scenarios: (a) The fraction of seeder individuals is varied (arbitrarily fixing χ_seed_=-1, χ_cont_=0.4). (b) The composition of SNP genotypes of the seeder population is changed by varying χ_seed_ (fixing *N*_seed_*/G*_size_=0.15 and χ_cont_=0.4). (c) The composition of SNP genotypes of the contact population is changed by varying χ_cont_ (fixing *N*_seed_*/G*_size_=0.15, χ_seed_=-1). Note, low SD implies high precision. Dashed lines represent analytical results and crosses refer to posterior estimates from simulated data (see Additional file 1). *N*_total_ refers to the total number of individuals.

Before describing these graphs in detail some general points can be made (irrespective of experimental design). Firstly, agreement between the analytical curves (dashed lines) and the simulation-based results (crosses, see Additional file 1 for details) is generally very good.

The notable exception to this is when the analytic expressions predict very large SDs (which manifests itself mostly in *a*_*f*_ because of the large SDs associated with this parameter). This discrepancy arises because the assumption of small SNP effects used in the analysis becomes invalid. In this regime analytic expressions tend to be conservative in that they suggest designs are poorer than they actually are. This shortcoming, however, is not very restrictive because it occurs in experimental designs in which very little information is anyway available (which is not where an experimenter would aim to design their experiment, moreover our analytical results would warn against it).

Secondly, the standard deviation of the recoverability estimates for *a*_*r*_ (right-hand column of graphs in Fig. 3) are general lower than the susceptibility estimates for *a*_*g*_ (left-hand column), which are themselves lower than the infectivity estimates (middle column). This was already previously noted in [15], and implies SNP-based differences in recoverability are the easiest to identify, followed by those in susceptibility, with SNP-based differences in infectivity the hardest to estimate.

In the case of recoverability, the reason estimates for *a*_*r*_ are significantly more precise than for the other two traits is because recovery times are usually less dispersed around their mean value than infection events (which have a wide exponential distribution) and also *a*_*r*_ does not suffer from the confounding which can make *a*_*g*_ and *a*_*f*_ much less certain in many circumstances.

Lastly, since precision is expected to scale as the square root of the total number of individuals, the SDs for an experiment containing 1000 individuals are expected to be a factor √10=3.2 times smaller than for an experiment containing 100 individuals. This can be seen on the graphs by an approximately constant distance between the black and grey dashed curves (note the log scale on the *y*-axis).

We now consider optimising the composition of seeder and contact individuals in the single contact group design for maximum precision. Figure 3a shows the case of varying the fraction of seeder individuals whilst fixing an arbitrary^9^ genetic makeup in the seeder and contact populations. Looking at the results for the SD in *a*_*g*_ (left-hand graph in Fig. 3a) we see, generally speaking, that the SD reduces for fewer seeders, which is unsurprising given that information regarding susceptibility comes from infection times of contacts. For a very low number of seeders, there is an additional effect due to the problem of epidemic extinction which leads to an increase in the SD (see Additional file 6). Consequently, careful consideration must be given as to how many seeders are necessary to successfully instigate epidemics (this will depend on *R*_*0*_).

In contrast, the SD for the SNP effect for infectivity *a*_*f*_ (as shown by the middle graph in Fig. 3a) has a different optimum. For a single contact group the analytical expression in Eq.(7) simplifies to

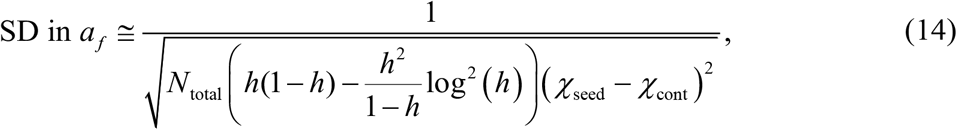

where, due to the approximation *ϕ*≈1, *h*=*N*_seed_/*G*_*size*_ is the fraction of seeder individuals (corresponding expressions for the other two traits are given in Additional file 7). The functional dependence on *h* gives the profile in the graph, which reaches its minimum when *h*=0.15 (hence the choice of 15% of seeders mentioned in Fig. 2a). In fact numerical analysis shows that, to a good approximation, this result remains true irrespective of the value of *R*_*0*_ (on which *ϕ* depends), as demonstrated in Additional file 8.

Figure 3b shows how SDs change as the result of varying the genetic makeup of the seeder population, as characterised by χ_seed_ (arbitrarily fixing χ_cont_=0.4 and using the optimum proportion of seeders *h*=0.15). Here the left-hand edge of the graph corresponds to the case when all seeders are *BB* and the right edge is when they are all *AA* (points in between represent a mixture of the two). We find that this variation makes very little difference to parameter precisions for *a*_*g*_ and *a*_*r*_, but the SD in *a*_*f*_ changes dramatically. This is because information regarding infectivity actually *relies* on differences in genetic makeup between the seeders and contacts. This is driven by the term (χ_seed_-χ_cont_)^2^ in Eq.(14), and we see that the analytical curves diverge in the limit χ_seed_→ χ_cont_.

This can be explained intuitively as follows. Suppose χ_seed_=-1, such that the seeders are solely *BB* individuals and χ_cont_=1, such that the contacts are solely *AA* individuals. Due to the fact that susceptible individuals become infected as the epidemic progresses, the infected population becomes more and more a mixture of *AA* and *BB* individuals. Thus, a comparison of how quickly^10^ the epidemic develops early on as compared to later gives direct evidence for the relative infectivity of *AA* compared to *BB* individuals, and hence *A* compared to *B* alleles^*11*^.

Figure 3c shows a contrasting design space, whereby the genetic makeup in the *contact* population varies between studies, thus we vary χ_cont_ (fixing χ_seed_=-1 and *h*=0.15). Here we find that the SD in *a*_*f*_ is minimised when χ_cont_=1, because this is as genetically different to the seeder population as possible. Unfortunately, however, here the SD in the susceptibility SNP effect *a*_*g*_ diverges (because there are no *BB* contacts and so no information regarding their relative susceptibility). Therefore to attain a reasonable precision for *a*_*g*_, χ_cont_ must be less than one. A reasonable choice is χ_cont_ ≈0.8, which, as can be seen from the right-hand side of Fig. 3c, leads to only a modest increase in the SD in *a*_*f*_ with the SD in *a*_*g*_ smaller than that for *a*_*f*_. This corresponds to around ≈10% *BB* individuals in the contact population, as quoted in Fig. 2a.

As it stands the single contact group design only contains *AA* and *BB* individuals, and so can provide no information regarding the dominance relationship between *A* and *B* alleles. Introduction of *AB* individuals into the seeders and contacts can inform Δ_*g*_ and Δ_*r*_, but it turns out that almost nothing can be inferred regarding the infectivity dominance factor Δ_*f*_ (results not shown). Consequently, this possibility is not further investigated here.

### Multiple contact groups - “Pure” design

Perhaps the most intuitive experimental design, which we term the “pure” design, is illustrated in Fig. 2b. When dominance is not being investigated, this consists of running replicates each of which consists of four separate groups in which the seeders and contacts are genetically homogeneous (*i*.*e*. “pure”) but consist of different combinations of genotype *AA* and *BB*. Such an approach is appealing because it allows conclusions to easily be drawn directly from the data. For example if epidemics in contact groups with similar genetic composition progress significantly faster in groups that contain *AA* seeders (*i*.*e*. groups 1 and 2 in Fig. 2b) compared to those which contain *BB* seeders (*i*.*e*. groups 3 and 4), this provides direct evidence that *A* alleles confer greater infectivity than *B* alleles (largely regardless of their relative susceptibility). Similarly, the relative susceptibility of *A* compared to *B* alleles can be found by comparing the relative epidemic speeds of groups where the infection was initiated by the same seeder genotypes, but the genotypes of the contact individuals differ (*i*.*e*. comparing groups 1 to 2, and 3 to 4 in Fig. 2b). This design has been implemented in [25] to estimate genotypic effects for a specific resistance marker on susceptibility, infectivity and recoverability.

Figure 4a shows how the SDs change as the fraction of seeder individuals is varied. For the pure design the analytical expression in Eq.(7) simplifies to

**Figure 4:**
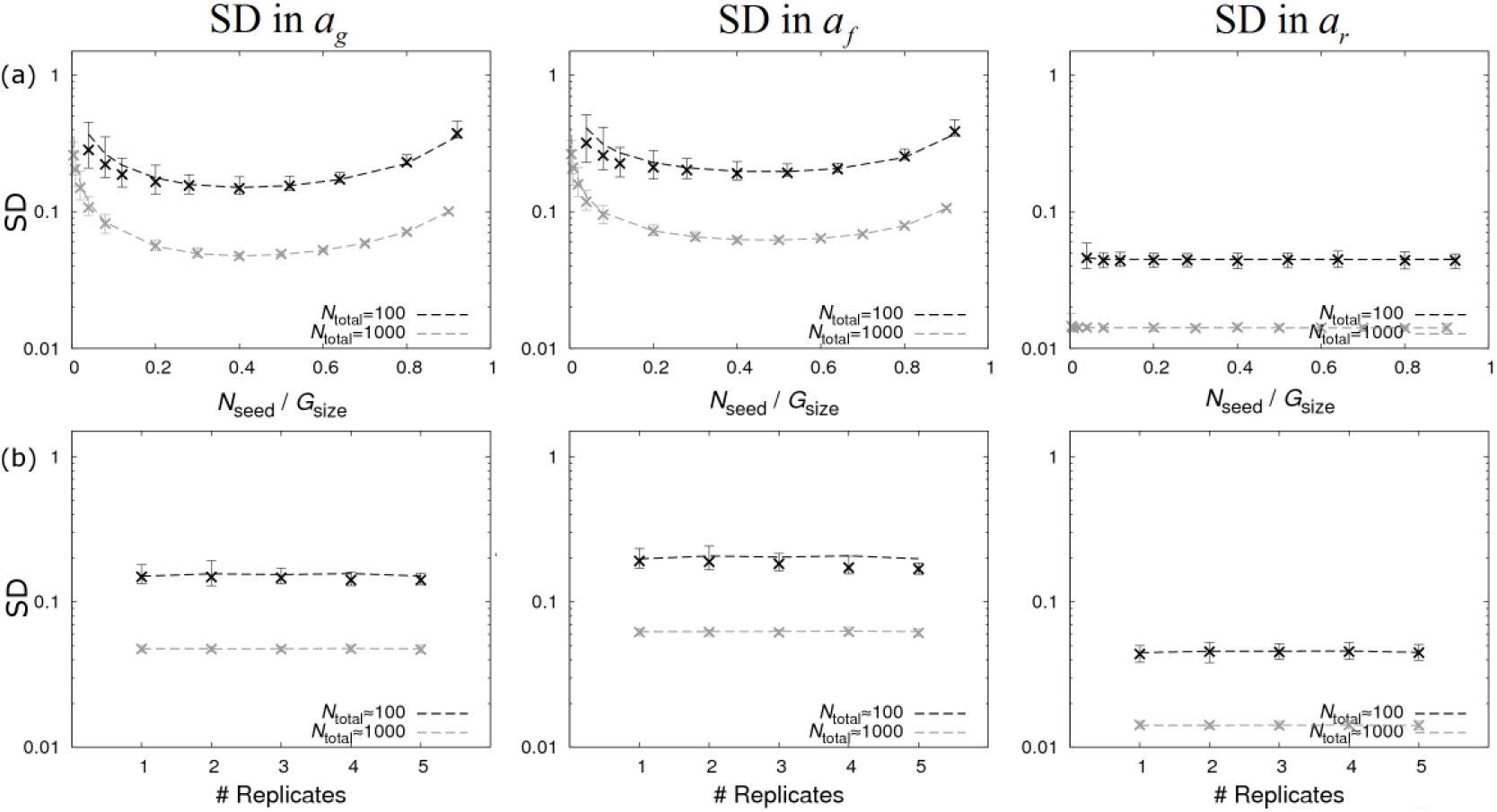
The “pure” design. Precision estimates for the pure design (without dominance) consisting of four contact groups per replicate with homogeneous seeder/contact SNP genotypes illustrated in Fig. 2b. The left, middle and right columns show graphs for standard deviations (SDs) in the SNP effects for susceptibility *a*_*g*_, infectivity *a*_*f*_ and recoverability *a*_*r*_ under different scenarios: (a) The fraction of seeder individuals in each contact group is varied. (b) The number of experimental replicates, each consisting of four contact groups, is varied (keeping to total number of individuals approximately fixed). Dashed lines represent analytical results and crosses refer to posterior estimates from simulated data (see Additional file 1). *N*_total_ refers to the total number of individuals.

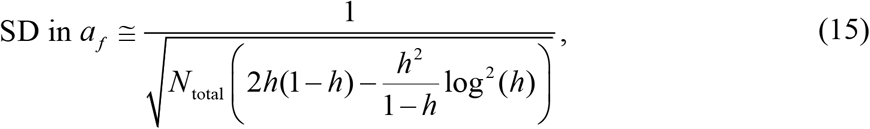

which is optimised when *h*=0.47, *i*.*e*. 47% of individuals should be seeders (expressions for the other two traits are given in Additional file 9). Again, this conclusion holds largely independent of *R*_*0*_, as demonstrated in Additional file 8, even when the proportion of contacts that become infected substantially reduces as *R*_*0*_→1. Comparing optimum solutions, we find that the SD in the SNP effect for infectivity *a*_*f*_ is around 1.6 times smaller for the pure design in Eq.(15) when compared to the single contact group design in Eq.(14). This means that disease transmission experiments using the pure design require a factor 2.5 times fewer individuals to generate an equivalent precision. This highlights the point that multiple groups substantially improve parameter estimates in the case of infectivity.

The design without dominance in Fig. 2b consists of four groups. Suppose instead we design an experiment with 8 groups by copying the same basic design over two replicates (each containing half the number of individuals). Such an approach is investigated in Fig. 4b, where the number of replicates is changed whilst fixing (approximately) the total number of individuals. We find almost no variation in inference precision. This suggests that the experimenter is free to choose the number of individuals per group (as usually dictated by practical considerations) and the number of replicates is informed by the total number of individuals available for the experiment as a whole. It should be noted, however, that design replication does play an important role in moderating the potential reduction in precision arising from group effects (as well as other systematic effects) that should be accounted for. This is discussed later in the “realistic model and data scenarios” section later.

The pure design above proved effective at precisely estimating *a*_*g*_, *a*_*f*_ and *a*_*r*_. However, because it doesn’t contain *AB* individuals, it cannot provide information regarding the dominance relationship between the *A* and *B* alleles. To address this we here introduce the pure design with dominance, as illustrated by the second design in Fig. 2b. This consists of running replicates of nine separate groups in which the seeders and contacts are genetically homogeneous *or* “pure” within each group (*i*.*e*. all individuals have the same genotype) but take different combinations of genotype *AA, AB* and *BB* across groups (see Additional file 9 for further details). The corresponding analytical equation for the SD in *a*_*f*_ is given by Eq.(15) multiplied by a constant factor √(3/2) ≈1.2. Hence this design leads to only a modest reduction in precision in SNP effects, whilst having the benefit of providing dominance parameter estimates.

### Multiple contact groups – “Mixed” design

The so-called “mixed design” uses replicates of the setup illustrated in Fig. 2c. Here contacts in group 1 contain a mixture of genotypes and the contacts in group 2 contain the complementary mixture (with *AA* and *BB* interchanged). Unlike the pure design, the mixed design doesn’t rely on a large number of seeders (in fact the smaller the better).

Results for the mixed design are shown in Fig. 5. The middle graph in Fig. 5a shows how the SD in *a*_*f*_ varies as the composition of SNP genotypes in the two contact populations is changed. Equation (7) simplifies to (assuming the first term in the denominator is negligible, which is valid in the limit of few seeders)

**Figure 5:**
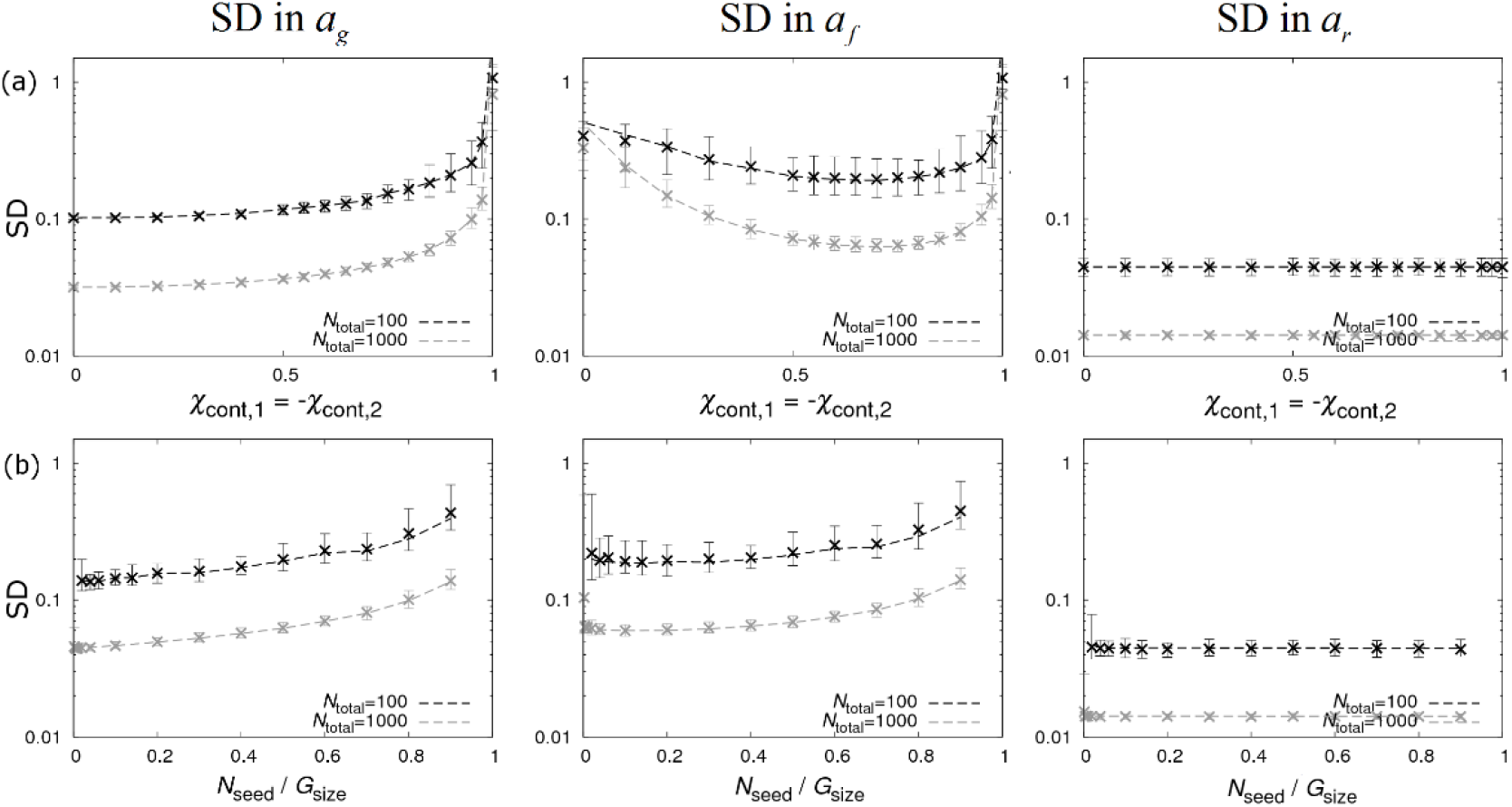
The “mixed” design. Precision estimates for the mixed design (without dominance) consisting of two contact groups per replicate with homogeneous seeder SNP genotypes and heterogeneous contact SNP genotypes illustrated in Fig. 2c. The left, middle and right columns show graphs for standard deviations (SDs) in the SNP effects for susceptibility *a*_*g*_, infectivity *a*_*f*_ and recoverability *a*_*r*_ under different scenarios: (a) The composition of SNP genotypes of the contact population in group 1 is changed by varying χ_cont,1_ whilst using the opposite value χ_cont,2_= -χ_cont,1_ in group 2 and *N*_seed_=3. (b) The fraction of seeders is varied (fixing χ_cont,2_= -χ_cont,1_=1/√2). Dashed lines represent analytical results and crosses come from posterior estimates from simulated data (see Additional file 1). *N*_total_ refers to the total number of individuals.

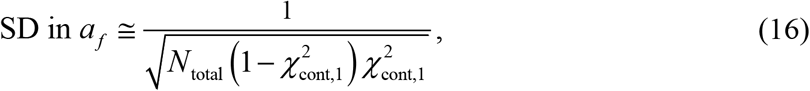

where χ_cont,2_=1-χ_cont,1_. This is minimised when χ_count,1_=1/√2 (or χ_count,1_=-1/√2), corresponding to 15% *BB* and 85% *AA* in the contact population of group 1 and ≈15% *AA* and ≈85% *BB* in the contact population of group 2 ^12^. Expressions for the SDs in the susceptibility and recoverability SNP effects are given in Additional file 10.

An intuitive explanation of how this experimental design works is as follows. As with most disease transmission experiments, when an infection occurs it is not known from which individual that infection originates. However, a key feature of the mixed design is that in group 1 there is a *greater probability* infections are initiated by *BB* individuals simply because there are more of them. Likewise, in group 2 *AA* individuals cause most of the infections. Consequently, if the epidemic in group 1 proceeds more quickly than in group 2, it is tempting to conclude that the *B* allele confers greater infectivity than *A*. Caution, however, is required due to potential confounding between infectivity and susceptibility (as a similar argument could be made to suggest that *B* alleles confers greater *susceptibility*). However considering the groups separately, the relative rate at which *AA* and *BB* individuals becomes infected gives direct evidence for differences in their susceptibility (irrespective of infectivity). This explains why it is necessary for groups to contain a mixture of genotypes, because this allows for any potential confounding to be overcome so allowing precise estimation of both *a*_*g*_ and *a*_*f*_. The right-hand graph in Fig. 5a shows that estimation of the recoverability SNP effect *a*_*r*_ remains the most precise of the three.

Figure 5b shows that precision is greatest when the fraction of seeders is small (subject to the extinction problem mentioned earlier). This highlights the fact that the mixed design predominantly gains information from infections within groups, whereas the pure design relies heavily of information gained from the initially infected seeders (Fig. 4a).

Again, because no *AB* individuals are present in the mixed design, as it stands it cannot be used to provide information regarding dominance. For completeness, therefore, we also include the mixed design with dominance, as illustrated in Fig. 2c (see Additional file 10 for further details).

For comparison, the expected precisions for the SNP effects and dominance parameters using the optimal designs is shown in Table 2.

**Table 2.**
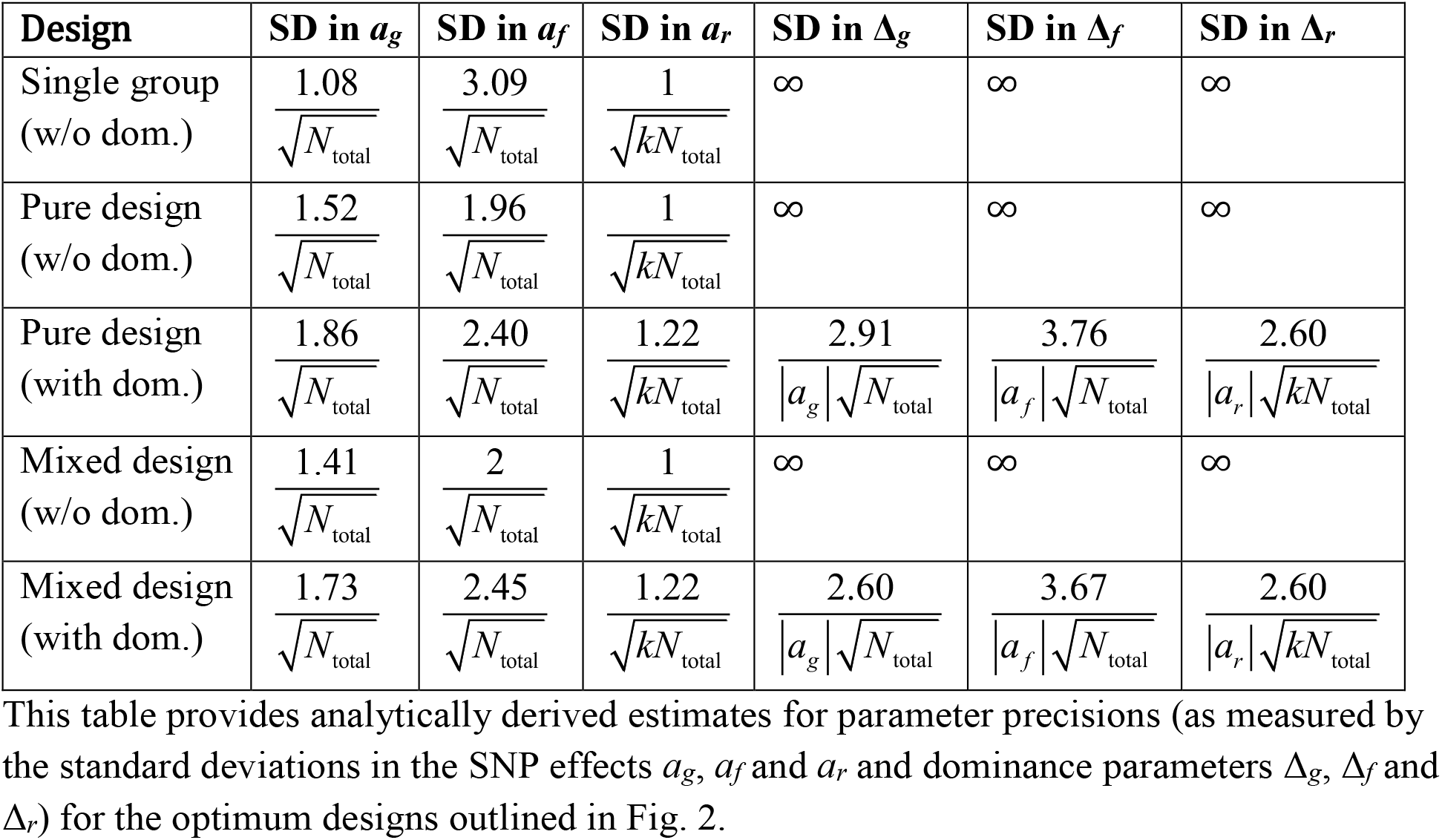
Parameter precision estimates.

### Other fixed effects

Note this paper has focused on estimating SNP effects, but it is important to point out that the analytical results and experimental designs outlined above are equally applicable to quantifying susceptibility, infectivity and recoverability differences coming from other systematic effects. For example if the influence of vaccination status is being studied the *AA* and *BB* genotypes can simply be replaced by “vaccinated” and “unvaccinated” classifications (note in this case there is no clear analogue of the *AB* genotype, so the dominance designs in Figs. 2b and 2c become redundant).

### Realistic model and data scenarios

Derivation of the analytical results above made use of some key simplifying assumptions. Termed the “best case scenario”, these included that infection and recovery times of individuals are precisely known and that the traits are only dependant on the SNP itself (*i*.*e*. the residuals and fixed and group effects in Eq.(2) were ignored). Here we assess the impact of relaxing these assumptions and investigate what implications this has on experimental design.

In particular five different sources of additional variation in the model/data were investigated separately: 1) introducing residual variation in traits **ε** in Eq.(2), 2) adding group effects *G*_*z*_ in Eq.(1), 3) adding a fixed effect (*e*.*g*. **X***b*_*g0*_) in Eq.(2)^13^, 4) analysing data with unknown infection times, and 5) assuming periodic disease status checks on individuals. Results of this investigation (details in Additional file 11) showed that whilst statistical power was found to be reduced (by varying amounts), the optimal design features expressed in Fig. 2 remained (approximately) unchanged^14^.

We now look at sequentially adding residual, group and fixed effect contributions to the basic SNP model to see how this impacts on the precision of SNP effects. Focusing on the optimal pure and mixed designs (without dominance), results of this are shown in Fig. 6. This assumes a fixed total number of individuals *N*_total_=1000 and looks at cases in which 4 and 12 (through replication) contact groups are used (for results with only 100 individuals see Additional file 12). The following conclusions can be drawn: Firstly, SDs in *a*_*r*_ are smallest (and least affected by additional sources of variation) followed by SDs in *a*_*g*_, then SDs in *a*_*f*_. Secondly, the contributions coming from the basic SNP-only model and the residual contributions are largely independent of choice of design (pure or mixed) or number of experimental replicates. Thirdly, the contribution from the group effect substantially reduces as the number of contact groups goes up from 4 to 12 and falls towards zero (results not shown) with increasing number of contact groups / replicates (see also Additional file 13). Fourthly, for the mixed design the group effect has almost no effect on precision of *a*_*g*_^15^, but a slightly larger effect on infectivity compared to the pure design. Lastly, fixed effects contribute very little provided they are not substantially correlated with the genotype of individuals (see Additional file 14).

**Figure 6:**
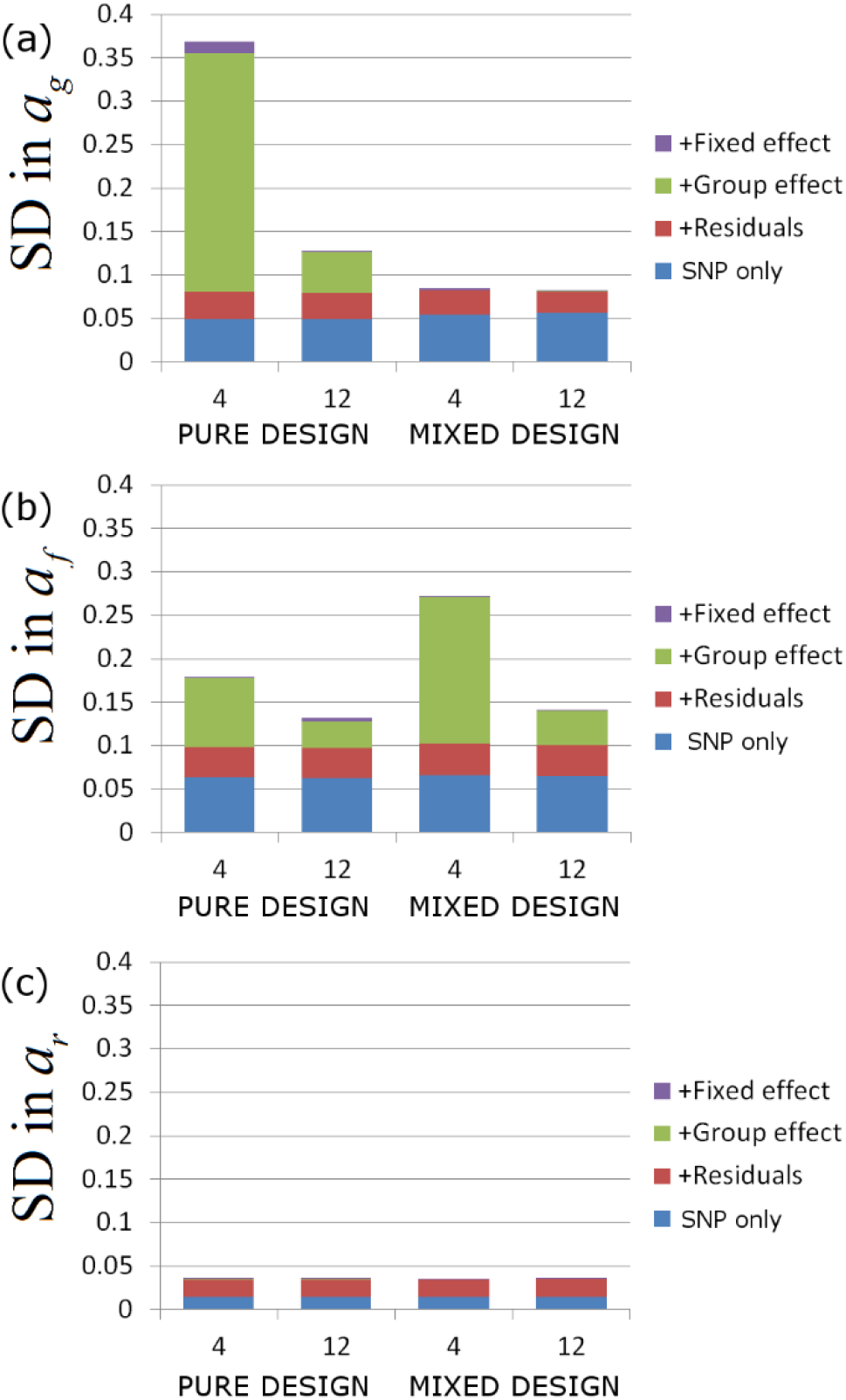
Partitioning contributions to standard deviations in SNP effects. Residuals, group effects and a fixed effect are sequentially added to the basic SNP only model (infection and recovery times assumed known). The corresponding increase in the SDs of the SNP effects is investigated for (a) the susceptibility *a*_*g*_, (b) the infectivity *a*_*f*_, and (c) the recoverability *a*_*r*_. For comparison, four different scenarios are investigated: a pure design (without dominance) with respectively *N*_group_=4 (*i*.*e*. a single replicate of the basic design) and *N*_group_=12 (*i*.*e*. 3 replicates of the basic design) and a mixed design (without dominance) with *N*_group_=4 (*i*.*e*. 2 replicates) and *N*_group_=12 (*i*.*e*. 6 replicates). In each case approximately 1000 individuals were partitioned equally amongst the contact groups. The residuals were chosen to have covariance matrix Σ_*gg*_=Σ_*ff*_=Σ_*rr*_=1, Σ_*gf*_=0.3, Σ_*gr*_=-0.4, and Σ_*fr*_=-0.2, the group effects had a SD of σ_G_=0.2, and the fixed effect (assumed to represent sex with gender randomly allocated) had size *b*_*g0*_=*b*_*f0*_=*b*_*r0*_=0.2. Results were found to be largely insensitive to these essentially arbitrary choices.

### Design tool software

The analytical expressions derived in this study together with the diverse experimental designs have been implemented in a user-friendly online software tool SIRE-PC (Susceptibility Infectivity Recoverability Estimation Precision Calculator), as can be seen in the screenshot in Fig. 7. This calculator takes as user inputs details of the experimental design (specifically the number and genetic composition of seeders and contacts in each group, the number of replicates being carried out, an estimate for the fraction of contacts expected to become infected *ϕ* ^16^ and *k*, the shape parameter charactering dispersion in recovery times) and outputs the total number of individuals used in the experiment and analytical estimates for the standard deviations in the SNP effect parameters *a*_*g*_, *a*_*f*_, and *a*_*r*_ in Eqs.(6)-(8) and the dominance parameters Δ_*g*_, Δ_*f*_ and Δ_*r*_, in Eqs.(10), (11), and (13). Note these expressions give the complete and no dominance cases, but the tool actually allows intermediate dominances to be investigated as well.

**Figure 7:**
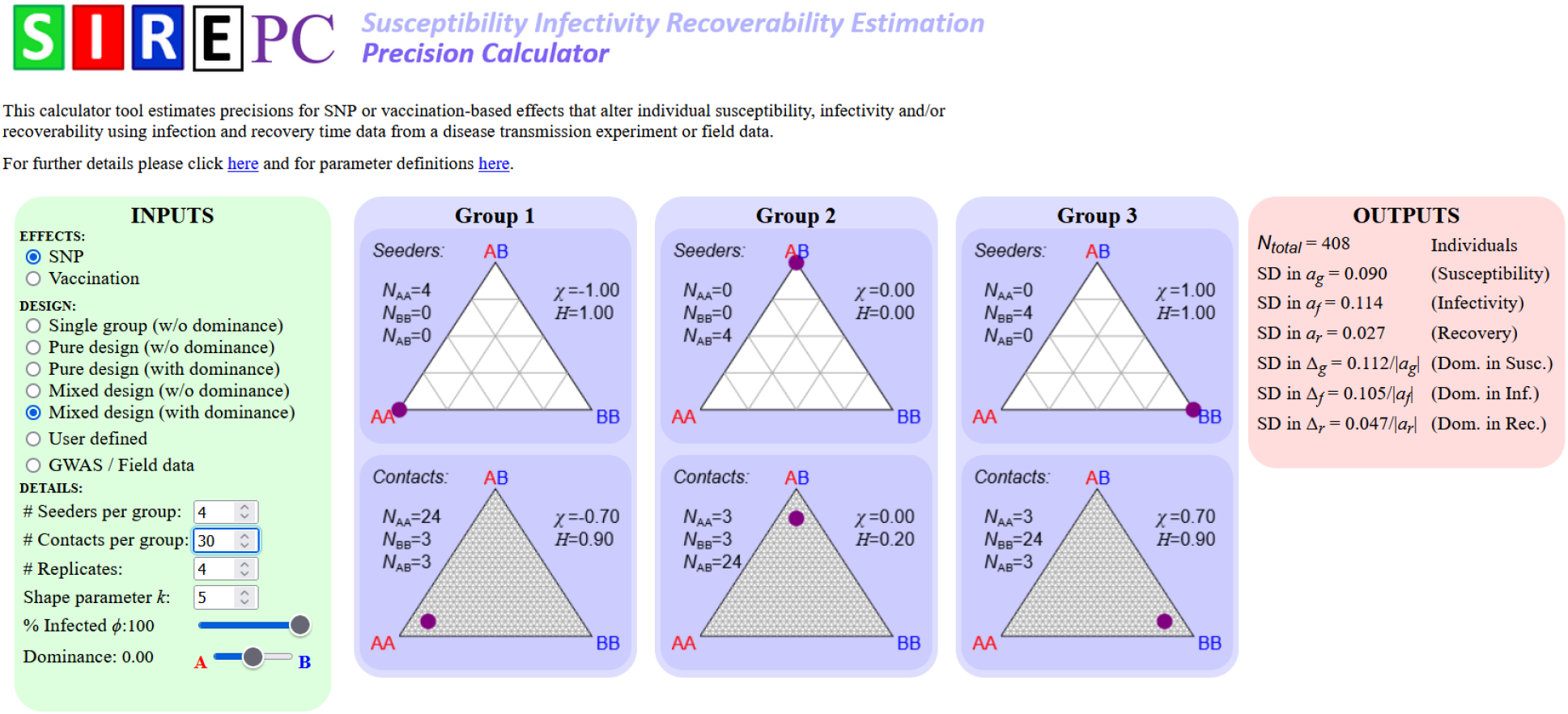
Precision Calculator tool. SIRE-PC (Susceptibility Infectivity Recoverability Estimation Precision Calculator) is an easy-to-use online software that calculates the analytical expressions provided in the results section to help aid experimental design.

Establishing appropriate experimental design is achieved by adjusting the input values, subject to any practical/logistic limitations (*e*.*g*. the number individuals per contact group may be fixed), with the aim of minimising the output standard deviations. To facilitate this process the tool includes the optimal experimental designs in Fig. 2, as well as providing the option for arbitrary user-defined designs to be investigated.

Additionally, the tool allows for precision estimates in the case of studying vaccination effects on the three host epidemiological traits, as well as application to GWAS and field data (see below).

## Discussion

There has been increasing acknowledgment within the livestock genetics community that infectious disease spread in populations may not only depend on individuals’ genetic susceptibility to infection, but also by their genetic infectivity and recoverability [25, 28, 30, 31, 33]. Whilst methods for estimating SNP and other genetic effects for these novel host traits from epidemiological data are emerging [15, 27-29], so far limited consideration has been given to the optimal design of transmission experiments. This study demonstrates that considerable improvements in the precision of SNP effect estimates associated with all three host epidemiological traits can be achieved through choosing the appropriate experimental design. This is here explicitly illustrated by means of considering a single SNP with potential effects on all three host epidemiological traits, but the same basic design features apply to any other categorical fixed effect (*e*.*g*. vaccination status of individuals).

This study provides analytical expressions for the precision of estimated SNP substitution and dominance effects associated with host susceptibility, infectivity and recoverability, which have been embedded into an online software tool to assist with the design of transmission experiments and for statistical power analyses for sample size requirements in experimental studies. To make the derivations tractable the calculations are shown for a best case scenario in which non-SNP contributions were ignored and infection and recovery times were known. Nonetheless, the derived expressions were found to be in strong agreement with numerical results obtained by performing inference on data from simulated epidemics that account for a range of complications and confounding likely present in real observations. The parameter that was found to be most difficult to precisely estimate was *a*_*f*_, which characterises differences in infectivity. Consequently, optimal experimental design largely focused on improving precision of *a*_*f*_ estimates. Three types of design emerged.

The first of these considered just a single contact group. It was found that, in principle at least, it is possible to infer *a*_*f*_ given a large enough population. However, this would not be a recommended option for a well-designed disease transmission experiment because multiple groups provide a way of significantly increasing statistical power.

When implementing designs with multiple groups, two fundamentally different strategies were considered: the “pure” and the “mixed” designs. The pure design, as show in Fig. 2b, uses different combinations of genotype in the seeders and contacts (in cases in which dominance is not being investigated this consists of replicates of 4 groups, and when it is, replicates of 9 groups). Choosing an approximately equal number of seeders and contacts led to similar precisions for the inferred SNP susceptibility and infectivity. On the other hand, the mixed design, as shown in Figs. 2c, relies on different frequencies of genotypes in the contact population across groups, with just a few seeders^17^ (here replicates of 2 or 3 groups are needed depending on whether dominance is being investigated or not).

For a fixed total number of individuals, both the pure and mixed designs were found to be similar in terms of their precision for estimating SNP effects (see Table 2). However two features of the mixed design makes it advantageous over the pure design: Firstly, when group random effects are included it was found to be significantly better at predicting susceptibility SNP effects^18^ (because differences in infection times of individuals of different genotypes provides direct evidence for susceptibility variation, irrespective of group effect), and secondly, it requires far fewer seeders. The latter is particularly important for disease transmission experiments in which seeders are artificially infected through injection, and so may behave differently to contacts who naturally acquire infection during the experiment^19^. Consequently, this paper advocates the mixed design as the best approach to take. Interestingly, this design is much less intuitive than the pure design, illustrating the importance of the derived analytical expressions above.

When implementing optimal multiple group designs, the analytical expressions in Table 2 suggest that the precision of parameters is largely independent of the number of individuals within each group, given a fixed total (for example running 4 groups using the pure design in Fig. 2b gave a very similar level of precision to running two replicates of 4 groups, each containing half the number of individuals). However in the more realistic scenario of significant random differences in disease transmission between groups (necessitating incorporation of the group effect term *G*_*z*_ in Eq.(1)), more replicates consisting of smaller groups was found to be beneficial (especially true for the infectivity SNP effect).

### Implications for genome-wide association studies

This study investigated experimental designs for which the composition of the seeder and contact populations are tailored specifically to estimate the effects of a specific SNP of interest on all three host epidemiological traits. Suppose, however, we are interested in performing a genome-wide association study (GWAS). In this case allocation of seeders and contacts according to their SNP genotype is not possible (because the SNP composition will be different for each SNP), so how should experiments be optimally designed? An analysis based on considering an arbitrary SNP in Hardy-Weinberg equilibrium [32] with *A* allele frequency *p* for a population consisting of unrelated individuals is given in Additional file 15.

This analysis shows that, as with the mixed design, precisions of both SNP effects and dominance parameters is maximised when epidemics are instigated with few seeders.

The following results are derived in Additional file 15:

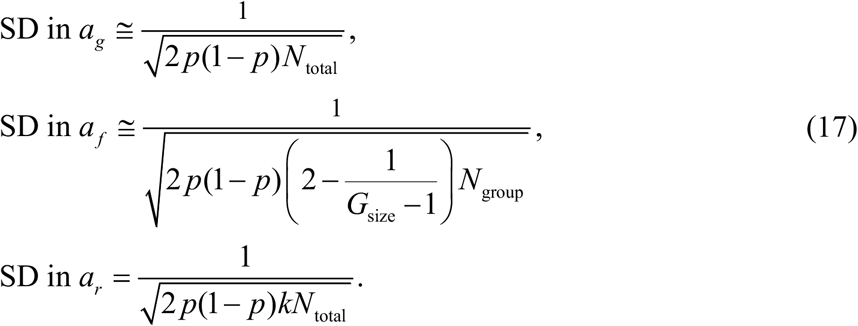

Note here that these SDs crucially depend on *p*, which makes sense in the limits p→0 and p→1 as the population becomes uniformly homozygote with no information regarding SNP effects. Crucially, compared to the results in Table 2, the SD in *a*_*f*_ now contains *N*_group_ in the denominator instead of *N*_total_. This means that increasing the number of individuals in each contact group no longer substantially increases the precision with which *a*_*f*_ can be estimated (a feature noted in [15]). Consequently when performing GWAS, many contact groups with fewer individuals lead to greater precision in estimating the SNP effect for infectivity (which, interestingly, is not the case for the susceptibility or recoverability SNP effects). Although the derivations were based on genetically unrelated individuals, these observations are expected to remain valid also in genetically structured populations.

### Field data

We now consider the possibilities and additional complications which arise when considering field data (that is data obtained from real-world disease outbreaks). As with the GWAS discussion above, here we do not have the luxury of being able to choose compositions of SNPs within different groups. Nevertheless, the analytical expression in Eq.(17) provide power calculations that can estimate what could, in principle, be inferred from field data (and again point to the fact that smaller groups sizes are more likely to yield good estimates for infectivity SNP effects). Seeders in our study are “index” cases which instigates epidemic, and the fact that there is usually just one index case fortunately coincides with the optimum for the precisions of SNP effects as discussed above.

In the case of investing vaccination effects, if there are some groups of individuals with high vaccination rate and others with low (or no) vaccination, this would naturally lend itself to something akin to the optimal mixed design proposed in this paper. Hence we would expect this sort of data to being highly informative about not only susceptibility and recoverability effects, but also infectivity.

It should be mentioned, however, that analysis of real-world data comes with addition complications: 1) proper accounting for related individuals, 2) the fact that groups are not entirely closed (*e*.*g*. cows in different field may share milking facilities) and 3) not all individual start in the susceptible state, especially for endemic diseases. Tackling these problems may require further development of the approaches outlined in this paper.

### Further considerations

In this paper epidemics were modelled using SIR dynamics, but it is important to point out that the results are equally applicable to diseases for which individuals do not recover (*i*.*e*. the SI model). Whilst more complicated compartmental model were not investigated, *e*.*g*. the inclusion of an exposed (infected but not infectious) state, the basic idea of accentuating variation across contact groups in the factor under study (*e*.*g*. by ensuring large differences in the genotypic composition in the mixed and pure designs) to increase variation in epidemic speed (which in turn provides evidence for variation in infectivity) is expected to remain valid.

Table 2 provides a useful guide as to the size of SNP effects that can be observed from a given experiment. It suggests that for datasets comprising 1000 individuals or fewer, only SNPs of large effects (typically explaining more than 15% of the total phenotypic variation) on the epidemiological host traits can be accurately estimated. SNPs of small to moderate effects would require significantly more data, in particular for infectivity. Although potentially challenging for livestock species due to the cost, such largescale experiments may be feasible for aquaculture and smaller laboratory species (*e*.*g*. insects).

Although SNPs with large effects on disease resistance have been identified [11, 22, 34], evidence suggests that disease resistance is mostly polygenic [35]. So far little is known about the genetic architecture underlying host infectivity and recoverability, but it seems reasonable to expect that these may be mostly under polygenic regulation too. This study has ignored any polygenic contributions to Eq.(2) [27] by assuming members from different families are distributed randomly across groups. Incorporation of such effects may lead to new insights into optimal disease transmission design, and will be the subject of future research.

### Conclusions

The aim of this paper was to identify optimal disease transmission experimental designs to estimate the effects of a particular SNP of interest (or other factors) on the susceptibility, infectivity and recoverability of individuals. It was found that whilst susceptibility and recoverability were relatively insensitive to design (both being clearly related to the infection and recovery times of individuals themselves), infectivity was not (because its effects are evident in the behaviour of others). In particular, to precisely estimate genetic effects on infectivity a so-called “mixed” design was identified, which specifies the optimal proportions of different genotypes in the contact populations of different contact groups. Replication of this basic design was found to be effective at reducing confounding arising from group effects.

An easy-to-use [software tool](https://theiteam.github.io/SIRE-PC.html) accompanying this paper can be used to aid experimental design by providing estimates for parameter precisions. The results shown here illustrate that such estimates are reliable and robust to noise and a range of potential confounding factors likely to be present in real-world disease systems.

## Declarations

## Ethics approval and consent to participate

Not applicable.

## Consent for publication

Not applicable.

## Availability of data and materials

All data generated or analysed during this study are included in this published article and its supplementary information files.

## Authors’ contributions

CMP derived the analytical expressions, generated the main results presented in the paper, and developed the accompanying software tool. GM and ADW contributed to the underlying methodology, testing and to the writing of the manuscript. SCB made instrumental contributions to the early stages of this project. All authors contributed to the writing and CMP, GM and ADW to the approval of the final manuscript.

## Acknowledgements

Not applicable.

## Additional files

**Additional file 1**

Title: Posterior inference from simulated data

Description: Provides details on how inference was performed on simulated datasets to generate results against which the analytical expressions could be compared.

**Additional file 2**

Title: Deriving the observed Fisher information matrix

Description: This section derives analytical expressions for the matrix in Eq.(4).

**Additional file 3**

Title: Inversion of the observed Fisher information matrix

Description: This shows how the observed Fisher information matrix in Eq.(4) is inverted to give the results in Eqs.(6) and (7).

**Additional file 4**

Title: Derivation of standard deviations for recoverability SNP effect

Description: We derive the standard deviation for the SNP parameter related to recoverability *a*_*r*_.

**Additional file 5**

Title: Dominance

Description: We derive standard deviations in the dominance parameters Δ_*g*_, Δ_*f*_ and Δ_*r*_.

**Additional file 6**

Title: Epidemic extinction

Description: Shows how the probability of epidemic extinction varies as a function of the number of seeder individuals *N*_seed_ for different basic reproductive ratios *R*_*0*_.

**Additional file 7**

Title: Further details for the single contact group design

Description: Provides analytical expressions for the standard deviations in the SNP effect parameters *a*_*g*_, *a*_*f*_ and *a*_*r*_ for the single contact group design.

**Additional file 8**

Title: Impact of changing R_0_

Description: Numerically investigates how changing R_0_ affects the optimal design choices outlined in Fig. 2.

**Additional file 9**

Title: Further details for the “pure” design

Description: Provides analytical expressions for the standard deviations in the SNP effect parameters *a*_*g*_, *a*_*f*_ and *a*_*r*_ and dominance parameters Δ_*g*_, Δ_*f*_ and Δ_*r*_ for the “pure” design.

**Additional file 10**

Title: Further details for the “mixed” design

Description: Provides analytical expressions for the standard deviations in the SNP effect parameters *a*_*g*_, *a*_*f*_ and *a*_*r*_ and dominance parameters Δ_*g*_, Δ_*f*_ and Δ_*r*_ for the “mixed” design.

**Additional file 11**

Title: Impact of realistic model/data scenarios on design

Description: Considering the optimal designs in Fig. 2, this additional file investigates the impact of separately introducing five additional sources of variation into the model/data (which, for the purposes of analysis, were ignored).

**Additional file 12**

Title: Partitioning contributions to the SDs of SNP effects

Description: Figure 6 in the paper shows the result of sequentially adding residuals, group effects and a fixed effect to the basic SNP model. This presents the corresponding results assuming only 100 individuals.

**Additional file 13**

Title: Design replication

Description: This additional file investigates how the reduction in precision when incorporating group effects can be moderated by means of design replication (that is repeating the same basic designs in Fig. 2 several times)

**Additional file 14**

Title: Fixed effect correlated with SNP

Description: This additional file investigates adding a single large fixed effect with elements in the design matrix **X** set in such a way as to give a certain degree of correlation with the SNP.

**Additional file 15**

Title: Genotype determined by Hardy-Weinberg equilibrium

Description: Rather than defining the proportions of genotypes in the seeder and contact populations, here we consider the case in which individual genotypes are randomly allocated with *A* allele frequency *p*, assuming Hardy-Weinberg equilibrium.

## Footnotes

1 In the case of field data these would be index cases.

2 *E*.*g*. χ_seed_=1 if seeders are solely *AA* individuals, χ_seed_=-1 if they are solely *BB* individuals, and somewhere in between in the general case.

3 “Fractional deviation” is defined by the exponential dependency in Eq.(1), *e*.*g. g*_*j*_=0.1 corresponds to individual *j* being a fraction ≈10% more susceptible than a population-wide reference.

4 Over and above those coming from the SNP and fixed effects themselves.

5 In the limit of large basic reproductive ratio *R*_*0*_, *ϕ* is 1, but for *R*_*0*_ close to 1, *ϕ* may be substantially smaller.

6 This manifests itself by the pattern of infection occurring early on.

7 *E*.*g*. in the case of livestock, pens are usually designed to house a certain number of animals.

8 In fact although not proven, numerical investigations suggest that experimental designs that have significant contributions from both terms in the denominator of Eq.(7) are no better than the ones shown in Fig. 2.

9 Here seeders are set to be *BB* individuals, corresponding to a homozygote balance of χ_seed_=-1, and contacts are set to be 70% *AA* and 30% *BB*, corresponding to χ_cont_=0.7-0.3=0.4.

10 Note here that relative “speed” has to explicitly take into account the number of infected individuals, as defined by the model.

11 Importantly, the genetic composition in the contact population does *not* change (in the example it remains solely *AA*), and so there is no confounding between susceptibility and infectivity.

12 Consideration of the small first term in the denominator of Eq.(7) shows that the seeder population is actually optimised when there are solely *AA* seeders in group 1 and solely *BB* seeders in group 2.

13 Specifically **X** is vector with elements randomly selected to be +0.5 and -0.5 representing male/female and *b*_*g0*_, *b*_*f0*_, *b*_*r0*_ are fixed effects that characterise how sex affects the three trait.

14 The only exception to this was that under the pure design the optimal fraction of seeders reduced to around 20% when group effects were introduced, but this increased back towards the optimal 47% with repeated experimental replicates.

15 This is because groups contain mixtures of genotype, and so their relative susceptibility can be directly estimated (irrespective of group effect).

16 Typically this is close to 1.

17 Typically two or three, sufficiently large to avoid the problem of epidemic extinction.

18 Although there is a corresponding small reduction in the precision for infectivity SNP effects.

19 This can be mitigated by the so-called extended experimental design (in which inoculated individuals are used to infect seeders prior to the start of the experiment), however, this not only increases the cost of the experiment, but also introduces additional uncertainty in the infection times of seeders.

## References

1. Visscher PM, Wray NR, Zhang Q, Sklar P, McCarthy MI, Brown MA, et al. 10 years of GWAS discovery: biology, function, and translation. Am J Hum Genet. 2017;101:5–22.

2. Stear M, Fairlie-Clarke K, Jonsson N, Mallard B, Groth D. Genetic variation in immunity and disease resistance in dairy cows and other livestock. Cambridge: Burleigh Dodds Science Publishing Limited; 2017.

3. Sharma A, Lee JS, Dang CG, Sudrajad P, Kim HC, Yeon SH, et al. Stories and challenges of genome wide association studies in livestock—a review. Asian Australas J Anim Sci. 2015;28:1371.

4. Shrestha V, Awale M, Karn A. Genome wide association study (GWAS) on disease resistance in maize. In: Disease Resistance in Crop Plants. Springer; 2019: p. 113–30

5. Freebern E, Santos DJ, Fang L, Jiang J, Gaddis KLP, Liu GE, et al. GWAS and fine-mapping of livability and six disease traits in Holstein cattle. BMC Genomics. 2020;21:1–11.

6. Biemans F, de Jong M, Bijma P. A genome-wide association study for susceptibility and infectivity of Holstein Friesian dairy cattle to digital dermatitis. J Dairy Sci. 2019;102:6248–62.

7. Houston RD, Haley CS, Hamilton A, Guy DR, Mota-Velasco JC, Gheyas AA, et al. The susceptibility of Atlantic salmon fry to freshwater infectious pancreatic necrosis is largely explained by a major QTL. Heredity. 2010;105:318–27.

8. Doeschl-Wilson A, Knap P, Opriessnig T, More S. Livestock disease resilience: From individual to herd level. Animal. 2021:100286.

9. Francis DH. Enterotoxigenic Escherichia coli infection in pigs and its diagnosis. J Swine Health Prod. 2002;10:171–75.

10. Authority EFS, Boelaert F, Hugas M, Ortiz Pelaez A, Rizzi V, Stella P, et al. The European Union summary report on data of the surveillance of ruminants for the presence of transmissible spongiform encephalopathies (TSEs) in 2015. EFSA Journal. 2016;14:e04643.

11. Boddicker N, Waide EH, Rowland R, Lunney JK, Garrick DJ, Reecy JM, et al. Evidence for a major QTL associated with host response to porcine reproductive and respiratory syndrome virus challenge. J Anim Sci. 2012;90:1733–46.

12. Psifidi A. The Genetics of Disease Resistance in Poultry. In: Poultry Health: A Guide for Professionals. CABI; 2021: p. 20

13. Oget C, Tosser-Klopp G, Rupp R. Genetic and genomic studies in ovine mastitis. Small Rumin Res. 2019;176:55–64.

14. Keeling MJ, Rohani P. Modeling infectious diseases in humans and animals. Princeton: Princeton university press; 2011.

15. Pooley CM, Marion G, Bishop SC, Bailey RI, Doeschl-Wilson AB. Estimating individuals’ genetic and non-genetic effects underlying infectious disease transmission from temporal epidemic data. PLoS Comput Biol. 2020;16:e1008447.

16. Hethcote HW, Van Ark JW. Epidemiological models for heterogeneous populations: proportionate mixing, parameter estimation, and immunization programs. Math Biosci. 1987;84:85–118.

17. Bitsouni V, Lycett S, Opriessnig T, Doeschl-Wilson A. Predicting vaccine effectiveness in livestock populations: A theoretical framework applied to PRRS virus infections in pigs. PLoS One. 2019;14:e0220738.

18. Doeschl-Wilson AB, Davidson R, Conington J, Roughsedge T, Hutchings MR, Villanueva B. Implications of host genetic variation on the risk and prevalence of infectious diseases transmitted through the environment. Genetics. 2011;188:683–93.

19. Raphaka K, Sánchez-Molano E, Tsairidou S, Anacleto O, Glass EJ, Woolliams JA, et al. Impact of genetic selection for increased cattle resistance to bovine tuberculosis on disease transmission dynamics. Front Vet Sci. 2018;5:237.

20. Hulst AD, de Jong MC, Bijma P. Why genetic selection to reduce the prevalence of infectious diseases is way more promising than currently believed. Genetics. 2021;217:iyab024.

21. Bijma P, Hulst AD, de Jong M. The quantitative genetics of the prevalence of infectious diseases: hidden genetic variation due to Indirect Genetic Effects dominates heritable variation and response to selection. Genetics. 2021:iyab141.

22. Houston RD, Haley CS, Hamilton A, Guy DR, Tinch AE, Taggart JB, et al. Major quantitative trait loci affect resistance to infectious pancreatic necrosis in Atlantic salmon (Salmo salar). Genetics. 2008;178:1109–15.

23. Moen T, Baranski M, Sonesson AK, Kjøglum S. Confirmation and fine-mapping of a major QTL for resistance to infectious pancreatic necrosis in Atlantic salmon (Salmo salar): population-level associations between markers and trait. BMC Genomics. 2009;10:1–14.

24. Moen T, Torgersen J, Santi N, Davidson WS, Baranski M, Ødegård J, et al. Epithelial cadherin determines resistance to infectious pancreatic necrosis virus in Atlantic salmon. Genetics. 2015;200:1313–26.

25. Doeschl-Wilson A, Anacleto O, Nielsen H, Karlsson-Drangsholt T, Lillehammer M, Gjerde B. New opportunities for genetic disease control: beyond disease resistance. In Proceedings of the World Congress on Genetics Applied to Livestock Production (Auckland). 11-16 Feb 2018

26. Gjedrem T, Rye M. Selection response in fish and shellfish: a review. Rev Aquac. 2018;10:168–79.

27. Anacleto O, Garcia-Cortés LA, Lipschutz-Powell D, Woolliams JA, Doeschl-Wilson AB. A novel statistical model to estimate host genetic effects affecting disease transmission. Genetics. 2015;201:871–84.

28. Anche MT, Bijma P, De Jong MC. Genetic analysis of infectious diseases: estimating gene effects for susceptibility and infectivity. Genet Sel Evol. 2015;47:85.

29. Biemans F, de Jong MC, Bijma P. A model to estimate effects of SNPs on host susceptibility and infectivity for an endemic infectious disease. Genet Sel Evol. 2017;49:53.

30. Welderufael B, Løvendahl P, De Koning D-J, Janss LL, Fikse W. Genome-wide association study for susceptibility to and recoverability from mastitis in Danish Holstein cows. Front Genet. 2018;9:141.

31. Lipschutz-Powell D, Woolliams JA, Bijma P, Doeschl-Wilson AB. Indirect genetic effects and the spread of infectious disease: are we capturing the full heritable variation underlying disease prevalence? PloS One. 2012;7:e39551.

32. Falconer D, Mackay T. Introduction to quantitative genetics. UK: Longman Group. 1996.

33. Tsairidou S, Anacleto O, Raphaka K, Sanchez-Molano E, Banos G, Woolliams J, et al. Enhancing genetic disease control by selecting for lower host infectivity. In Proceedings of the World Congress on Genetics Applied to Livestock Production (Auckland). 11-16 Feb 2018

34. Fife M, Howell J, Salmon N, Hocking P, Van Diemen P, Jones M, et al. Genome-wide SNP analysis identifies major QTL for Salmonella colonization in the chicken. Anim Genet. 2011;42:134–40.

35. Manolio TA, Collins FS, Cox NJ, Goldstein DB, Hindorff LA, Hunter DJ, et al. Finding the missing heritability of complex diseases. Nature. 2009;461:747–53.

36. Gillespie DT. Exact stochastic simulation of coupled chemical reactions. J Phys Chem. 1977;81:2340–61.

37. Gibson GJ, Renshaw E. Estimating parameters in stochastic compartmental models using Markov chain methods. Math Med Biol: A Journal of the IMA. 1998;15:19–40.

38. O’Neill PD, Roberts GO. Bayesian inference for partially observed stochastic epidemics. J R Stat Soc Ser A Stat Soc. 1999;162:121–9.

39. Efron B, Hinkley DV. Assessing the accuracy of the maximum likelihood estimator: Observed versus expected Fisher information. Biometrika. 1978;65:457–83.

