## Additional file 1 for "Optimal experimental designs for estimating genetic and non-genetic effects underlying infectious disease transmission"

### Posterior inference from simulated data

This additional file provides details on how inference was performed on simulated datasets to generate results against which the analytical expressions could be compared (specifically the crosses in Figs. 3-5 and the bar charts in Fig. 6).

**Simulation** - Data were generated by simulating epidemics in each contact group using a modified Gillespie algorithm [36], as described below. The following set of epidemiological parameters were used: $\beta$=0.3/($G_{size}$-1), $\gamma$=0.1 and $k$=5. Since $G_{size}$ is the total number of individuals per contact group, this choice corresponds to a fixed basic reproductive ratio of $R_0$= $\beta(G_{size}$-1)/γ=3. For the SNP effects: $a_g$=$a_f$=$a_r$=$\Delta_g$=$\Delta_f$ =$\Delta_r$=0. Although this might seem a somewhat contrived choice it is important to note that these values don't represent a particularly special case. This is because here the aim is to estimate parameter precisions rather than parameter values themselves, and the analytical expressions show that these precisions are, to a first approximation at least, independent of the model parameters θ (this independence is empirically demonstrated in Fig. 4 of [15]).

**Data** - The data taken from the simulations is, for the most part, just the infection and recovery times of those individuals which became infected. However in Additional file 11, where more realistic data scenarios are investigated, other possibilities are considered, such as only knowing infection times or periodic checking of disease status (*e.g.* through diagnostic tests results).

**Inference** - This is performed using a recently developed software tool SIRE (Susceptibility, Infectivity and Recoverability Estimation) [15], which incorporates a Bayesian methodology that is flexible to different data types (*e.g.* infection and/or recovery times of individuals or diagnostic test results) and can account for data uncertainty in a statistically consistent way. It incorporates a Markov chain Monte Carlo (MCMC) algorithm. For the estimations in this study $10^3$ burn-in iterations were performed followed by $10^5$ iterations, sufficient for chains to become well mixed.

To account for stochastic variation, each of the points in Figs. 3-5 summarises results of inference performed on 50 simulated epidemic datasets (where each cross indicates the mean posterior SD over all 50 replicates and the error bars represent 95% variation in this mean).

### Simulating epidemics in genetically heterogeneous contact groups

The Doob-Gillespie algorithm [36] provides a means of taking into account inherent stochasticity in Markovian compartmental models (*i.e.* models for which the transition rates depend solely on the current state of the system). The model used in this paper combines Markovian infection transitions with more realistic non-Markovian recovery dynamics. Below we describe how these recovery events are incorporated into the standard Doob-Gillespie framework.

The purpose of this procedure is to build up a time-ordered sequence of infection and recovery event times indexed by event number *e*. The following notation is used: $t_e$ is the event time, $x_e$ is the event type (infection *in.* or recovery *re.*), $j_e$ is the affected individual, and $t_j^I$ $t_j^R$ are the infection and recovery times for individual $j$, respectively.

**Initialization:** Each epidemic is assumed to be started by one (or potentially more) initially infected individual $j$ at some initial time point $t_0$. The infection duration $\delta t_j$ for this individual is drawn from a gamma distribution parameterised in terms of an individual-based mean and shape parameter:

$$\delta t_j \sim Gamma(w_j, k) \qquad \text{(A1)}$$

(note, the dependency of $w_j$ on θ is given through Eqs.(1)-(3). This allows us to set $t_j^I = t_0$ and $t_j^R = t_0 + \delta t_j$. Individual $j$ is then placed onto a list $\mathcal{R}$, which represents all currently infected individuals. We set event index to $e=1$.

**Step 1**: Calculate the time to the next infection event. This is done by first evaluating the total transition rate that any individual becomes infected

$$\Lambda = \sum_s \lambda_s, \qquad \text{(A2)}$$

where the sum $s$ goes over all currently susceptible individuals and the force of infection $\lambda_s$ (which gives the probability per unit time of $s$ becoming infected) is given by Eq.(1). In accordance with a Poisson process, the time to the next infection event is generated by drawing a sample from the exponential distribution $\Lambda e^{-\Lambda \Delta t}$. In practice, this is achieved by selecting an inter-event time using

$$\Delta t = -\frac{\log(u)}{\Lambda}, \qquad \text{(A3)}$$

where $u$ is a (uniform) randomly generated number between $0$ and $1$. The new event time is then defined by

$$t^{new} = t_{e-1} + \Delta t. \qquad \text{(A4)}$$

**Step 2**: Choosing the event type. For the SIR model two possibilities exist:

**a)** If $t^{\text{new}}$ is greater than the smallest recovery time of all the individuals in $\mathcal{R}$, which we label $j_{min}$, then we remove $j_{\min}$ from $\mathcal{R}$ and set

$$t_e = t_{j_{\min}}^R, \;\; x_e = re., \;\; j_e = j_{\min}. \qquad \text{(A5)}$$

**b)** Otherwise, we set

$$t_e = t^{new}, \;\; x_e = in., \qquad \text{(A6)}$$

and select the actual individual that becomes infected with probability

$$\text{Prob}(j_e = s) = \frac{\lambda_s}{\Lambda}. \qquad \text{(A7)}$$

The infection duration $\delta t_{j_e}$ for $j_e$ is sampled using Eq.(A1), and the infection and recovery times are set to

$$\begin{aligned} t_{j_e}^{I} &= t_e, \\ t_{j_e}^{R} &= t_e + \delta t_{j_e}. \end{aligned} \tag{A8}$$

Individual $j_e$ is then placed onto the list $\mathcal{R}$.

**Step 3:** Increment $e$ and jump to step 1 if there are any remaining infected individuals.

**End:** Insert recovery times for any remaining individuals $j$ in $\mathcal{R}$

$$t_e = t_j^R, \;\; x_e = re., \;\; j_e = j, \tag{A9}$$

incrementing $e$ after each addition.

The above algorithm describes simulation of a single contact group. The procedure is repeated separately for each contact group (which is valid because groups are assumed to be closed) to generate a complete set of infection and recover events ξ for the entire population of individuals.
