## Additional file 2 for "Optimal experimental designs for estimating genetic and non-genetic effects underlying infectious disease transmission"

### Deriving the observed Fisher information matrix

Here we derive analytical expressions for the matrix in Eq.(4). For tractability this analysis makes some simplifying assumptions: if an individual becomes infected both infection and recovery times are precisely known, epidemics are continued until they die out (*i.e.* no censoring), and the traits are dependant only on the SNP itself (*i.e.* the fixed effects $\mathbf{b}_g$, $\mathbf{b}_f$, $\mathbf{b}_r$, group effects $\mathbf{G}$ and residuals $\boldsymbol{\varepsilon}_g$, $\boldsymbol{\varepsilon}_f$, $\boldsymbol{\varepsilon}_r$ in Eq.(2) are all ignored). We subsequently refer to this as the "best case scenario", because it represents the idealised case in which information is maximised in any given experimental setup.

Whilst the best case scenario is not valid in most real world applications, this approach is useful for two reasons: Firstly, it provides upper limits on the precision with which parameters can be estimated. Secondly, it might be expected that optimal experimental design under this idealised scenario is likely to be similar to cases in which some, or possibly all, of the assumptions are violated (note, whilst optimal design is expected to be approximately unchanged, the statistical power to make inference will substantially reduce as data becomes more uncertain). This premise is explicitly tested below and found to be more or less valid (with some minor exceptions discussed later).

From [15], the likelihood of generating observed event data ξ given a set of model parameters θ=($\beta$, $\gamma$, $k$, $a_g$, $a_f$, $a_r$, $\Delta_g$, $\Delta_f$, $\Delta_r$) is [37,38]

$$L(\xi\,|\,\theta)=\prod\nolimits_{z}\Big[\Big(\prod\nolimits_{j\in z}\lambda_j\Big)\Big(\prod\nolimits_{e\in z}e^{-\Lambda_e(t_e)\times(t_e-t_{e-1})}\Big)\times\Big(\prod\nolimits_{m\in z}F_\Gamma(\delta t_m\,|\,w_m,k)\Big)\Big]. \quad \text{(A1)}$$

Note, the functional dependence on the parameters θ comes through the force of infection $\lambda_j$ and mean infection duration $w_m$ in Eq.(1), which themselves depend on $\mathbf{g}$, $\mathbf{f}$ and $\mathbf{r}$ in Eq.(2). The indices in Eq.(A1) are now described: $z$ goes over contact groups and within each group $j$ goes over individuals that become infected (*excluding* seeders), $m$ also goes over infected individuals (*including* seeders) and $e$ goes over both infection and recovery events (with corresponding event times $t_e$). The notation $j\in z$ indicates all those individuals $j$ that are in group $z$, and the symbol "∈" (contained within) is used in a similar context throughout this paper. The force of infection $\lambda_j$ is given by Eq.(1), immediately prior to individual $j$ becoming infected.

The gamma distributed probability density function $F_\Gamma$ in Eq.(A1) gives the probability an individual is infected for duration $\delta t_m$ given a mean duration $w_m$ and shape parameter $k$. The time dependent total rate of infection events is given by

$$\Lambda_e(t_e)=\sum\nolimits_{s\in e}\lambda_s, \quad \text{(A2)}$$

where $s$ extends over all susceptible individuals in group $z$ immediately prior to event time $t_e$.

The likelihood in Eq.(A1) factorizes into two contributions: a part that depends solely on the infection process and a part which depends solely on the recovery process. This implies that expressions for precisions in parameters relating to infection and recovery, respectively, can be derived independently of each other. These derivations are shown below and later, in Additional file 4, analysis is performed for the recoverability parameters.

#### Infection parameters

Substituting the infection rates from Eq.(1) into Eq.(A1) and making use of Eqs.(3) and (A2) (ignoring residuals, fixed and group effects, as assumed under the best case scenario) yields the distribution

$$\pi(\beta, a_g, a_f, \Delta_g, \Delta_f \mid \xi) \propto \beta^{N_I} e^{-\beta J} \prod_z \left[ \prod_{j \in z} \left( e^{g_j^{\text{SNP}}} \sum_{i \in j} e^{f_i^{\text{SNP}}} \right) \right], \tag{A3}$$

where $N_I$ is the total number of observed infection events across all groups[1]. The index $z$ goes over the contact groups and within each group, $j$ goes over individuals that become infected. Here $g_j^{\text{SNP}}$ is the fractional deviation in susceptibility of individual $j$, as defined in Eq.(3), and $f_i^{\text{SNP}}$ is the corresponding deviation in infectivity of individual $i$, where $i \in j$ sums over all contact group members of $j$ that were infected just before $j$ becomes infected. Lastly

$$J = \sum_z \left( \sum_{e \in z} \left[ \left( \sum_{s \in e} e^{g_s^{\text{SNP}}} \right) \left( \sum_{i \in e} e^{f_i^{\text{SNP}}} \right) (t_e - t_{e-1}) \right] \right), \tag{A4}$$

where the indices $s$ and $i$ sum over the susceptible and infected populations immediately prior to observed event time $t_e$.

The expression in Eq.(A3) is gamma distributed with respect to $\beta$, so this model parameter can be integrated out to give

$$\pi(a_g, a_f, \Delta_g, \Delta_f \mid \xi) \propto N_I! J^{-N_I - 1} \prod_z \left[ \prod_{j \in z} \left( e^{g_j^{\text{SNP}}} \sum_{i \in j} e^{f_i^{\text{SNP}}} \right) \right]. \tag{A5}$$

To avoid overly complicated analytical expressions we consider the case in which no dominance is assumed, *i.e.* $\Delta_g=\Delta_f=0$ (the situation of complete dominance is discussed later), and focus on the distribution for the parameters $a_g$ and $a_f$. In particular, this is approximated by a multivariate normal distribution

$$\pi(a_g, a_f \mid \xi, \Delta_g, \Delta_f) \cong N\left( \langle \mathbf{a} \rangle, \mathbf{M}^{-1} \right) \tag{A6}$$

centred on a vector given by the distribution means $\langle \mathbf{a} \rangle = (\langle a_g \rangle, \langle a_f \rangle)$ and with covariance matrix $\mathbf{M}^{-1}$ (this approximation is equivalent to performing a Taylor expansion in the log-likelihood and truncating at second order). $\mathbf{M}$ is known as the "observed Fisher information matrix" [39], and is defined by

$$M_{vu} = -\frac{\partial^2}{\partial a_v \partial a_u} \log\left[ \pi(a_g, a_f \mid \xi, \Delta_g, \Delta_f) \right], \tag{A7}$$

where indices $v$ and $u$ take the values "$g$" or "$f$".

To make progress we further assume that both $a_g$ and $a_f$ are small and take a continuum limit[2] (although it is later found in the simulation study that the derived analytical expressions are

---

[1] All infection events that occur to contacts during the experiment, *i.e.* discounting seeders.

[2] In other words, quantities that are in reality subject to stochastic variation are approximated by their values under the assumption of no noise.

remarkably accurate, even when these conditions are not satisfied). To simplify matters further, we assume that each contact group contains an identical number of seeders $N_{\mathrm{seed}}$ (*i.e.* initially infected individuals) and contacts $N_{\mathrm{cont}}$ (*i.e.* initially susceptible individuals)[3].

Under the zero-dominance assumption, the SNP effects from Eq.(3) can be expressed in the more concise format:

$$g_j^{\mathrm{SNP}} = \kappa_j a_g , \quad f_j^{\mathrm{SNP}} = \kappa_j a_f , \qquad \text{(A8)}$$

where $\kappa_j$={1,0,-1} depending on whether the genotype of individual $j$ is {*AA*,*AB*,*BB*}.

Taking the log of Eq.(A5) and substituting in Eq.(A8) gives (up to a constant and assuming $N_I \gg 1$)

$$\log\left(P\right) = -N_I \log(J) + \sum_z \left[ \sum_{j\in z} \left( \kappa_j a_g + \log\left( \sum_{i\in j} e^{\kappa_i a_f} \right) \right) \right], \qquad \text{(A9)}$$

where we use the abbreviation $P=\pi(a_g,a_f|\xi,\Delta_g,\Delta_f)$. Taking the double derivative of this expression w.r.t. $a_g$ gives

$$\begin{aligned} \frac{\partial^2 \log\left(P\right)}{\partial a_g^2} = &-\frac{N_I}{J}\left( \sum_z \left( \sum_{e\in z} \left[ \left( \sum_{s\in e} \kappa_s^2 e^{\kappa_s a_g} \right) \left( \sum_{i\in e} e^{\kappa_i a_f} \right) \left( t_e - t_{e-1} \right) \right] \right) \right) \\ &+ \frac{N_I}{J^2}\left( \sum_z \left( \sum_{e\in z} \left[ \left( \sum_{s\in e} \kappa_s e^{\kappa_s a_g} \right) \left( \sum_{i\in e} e^{\kappa_i a_f} \right) \left( t_e - t_{e-1} \right) \right] \right) \right)^2 . \end{aligned} \qquad \text{(A10)}$$

To make progress we take the limit when $a_g$ and $a_f$ are small (allowing the exponentials in Eq.(A10) to be approximated by one) and assume that quantities which are in reality subject to stochastic variation are approximated by their continuum counterparts (with no noise). Under this scenario the fraction of susceptible individuals $\nu_{\omega,z}^S(t)$ in the three genotypes $\omega$={*AA*,*AB*,*BB*} remains constant as a function time $t$[4]. Consequently (remembering that $s$ and $i$ sum over the susceptible and infected individuals immediately prior to event $e$ in contact group $z$) the following approximations can be made:

$$\begin{aligned} &\sum_{s\in e} \kappa_s e^{\kappa_s a_g} \cong \left( \nu_{AA,z}^S(t) - \nu_{BB,z}^S(t) \right) S_e \cong \chi_{\mathrm{cont},z} S_e, \\ &\sum_{s\in e} \kappa_s^2 e^{\kappa_s a_g} \cong \left( \nu_{AA,z}^S(t) + \nu_{BB,z}^S(t) \right) S_e \cong H_{\mathrm{cont},z} S_e, \\ &\sum_{s\in e} e^{\kappa_s a_g} \cong S_e, \\ &\sum_{i\in e} e^{\kappa_i a_g} \cong I_e, \end{aligned} \qquad \text{(A11)}$$

where $S_e$ and $I_e$ are the total number of susceptible and infected individuals immediately prior to event $e$, $\chi_{\mathrm{cont},z} = \nu_{AA,z}^S(0) - \nu_{BB,z}^S(0)$ is defined to be the homozygote balance in the contacts in group

[3] Note, this assumption is not strictly necessary but greatly simplifies the derived expressions.

[4] Because when $a_g$ is small, individuals in the three genotypes experience the same force of infection, so in the continuum limit the proportions of the genotypes remain unchanged.

$z$ (a quantity which goes from 1 when all contacts are *AA* to -1 when they are all *BB*) and $H_{\text{cont},z} = v^S_{AA,z}(0) + v^S_{BB,z}(0)$ is the corresponding homozygosity.

Substituting Eq.(A11) into (A10) gives

$$\begin{aligned} \frac{\partial^2 \log(P)}{\partial a_g^2} = &-N_I\left(\frac{1}{J}\sum_z\left(H_{\text{cont},z}\sum_{e\in z}\left[S_eI_e\left(t_e - t_{e-1}\right)\right]\right)\right) \\ &+N_I\left(\frac{1}{J}\sum_z\left(\chi_{\text{cont},z}\sum_{e\in z}\left[S_eI_e\left(t_e - t_{e-1}\right)\right]\right)\right)^2. \end{aligned} \tag{A12}$$

In the continuum limit, and assuming that epidemics are observed until completion, leads to

$$\sum_{e\in z}\left[S_eI_e\left(t_e - t_{e-1}\right)\right] \cong \int_{t=0}^{\infty} S_z(t)I_z(t)dt, \tag{A13}$$

where $S_z(t)$ and $I_z(t)$ now vary continuously in time. Summing $\lambda_j$ in Eq.(1) over all individuals (and remembering that here we are neglecting group effects and taking the limit $a_g \to 0$ and $a_f \to 0$) shows that the continuum dynamics for $S_z$ are determined by

$$\frac{dS_z}{dt} = -\beta S_z I_z. \tag{A14}$$

Note, this represents the standard SIR equation for a genetically homogenous population. Substituting Eq.(A14) into (A13) leads to

$$\begin{aligned} \sum_{e\in z}\left[S_eI_e\left(t_e - t_{e-1}\right)\right] &\cong -\int_{t=0}^{\infty}\frac{1}{\beta}\frac{dS_z}{dt}dt \\ &\cong -\int_{N_{\text{cont}}}^{N_{\text{cont}}-\phi N_{\text{cont}}}\frac{1}{\beta}dS_z \\ &\cong \frac{1}{\beta}\phi N_{\text{cont}}, \end{aligned} \tag{A15}$$

where $\phi$ is the expected fraction of contacts that become infected during the experiment. Placing this into Eq.(A12) and noting that

$$J \cong \sum_z\left(\sum_{e\in z}\left[S_eI_e\left(t_e - t_{e-1}\right)\right]\right) \cong \frac{N_{\text{group}}}{\beta}\phi N_{\text{cont}} \tag{A16}$$

gives the final result

$$\begin{aligned} \frac{\partial^2 \log(P)}{\partial a_g^2} &= -\phi N_{\text{cont}}\left(\sum_z H_{\text{cont},z}\right) + \frac{\phi N_{\text{cont}}}{N_{\text{group}}}\left(\sum_z \chi_{\text{cont},z}\right)^2 \\ &= -N_{\text{group}}\phi N_{\text{cont}}\left(\langle H_{\text{cont}}\rangle - \langle \chi_{\text{cont}}\rangle^2\right), \end{aligned} \tag{A17}$$

where angle brackets denote averaging over contact groups.

Similarly, making use of the result

$$\begin{aligned}\sum_{e\in z}\left[\chi_{I,z}(t)S_eI_e\left(t_e-t_{e-1}\right)\right] &\cong \int_{t=0}^{\infty}\chi_{I,z}(t)\frac{1}{\beta}\frac{dS_z}{dt}dt \\ &\cong -\int_{N_{\text{cont}}}^{N_{\text{cont}}-\phi N_{\text{cont}}}\chi_{I,z}(t)\frac{1}{\beta}dS_z \\ &\cong \tfrac{1}{\beta}\sum_{j\in z}\chi_{I,z}(t),\end{aligned} \tag{A18}$$

where $\chi_{I,z}(t)=v^I_{AA,z}(t)-v^I_{BB,z}(t)$ is the time varying homozygote balance in the infected population and $j$ goes over the infection events in group $z$, other partial derivatives can be derived:

$$\begin{aligned}\frac{\partial^2\log(P)}{\partial a_g\partial a_f} &= -\sum_z\left[\chi_{\text{cont},z}\sum_{j\in z}\chi_{I,z}(t)\right]+\langle\chi_{\text{cont}}\rangle\left[\sum_z\left(\sum_{j\in z}\chi_{I,z}(t)\right)\right], \\ \frac{\partial^2\log(P)}{\partial a_f^2} &= -\sum_z\left[\sum_{j\in z}\chi^2_{I,z}(t)\right]+\frac{1}{N_I}\left[\sum_z\left(\sum_{j\in z}\chi_{I,z}(t)\right)\right]^2.\end{aligned} \tag{A19}$$

The reason that $\chi_{I,z}(t)$ changes in time (as opposed to the homozygote balance in the susceptible population $\chi_{S,z}(t)$, which takes the constant value $\chi_{\text{cont},z}$ within a given contact group) can be explained with the following example. Suppose initially the infected population is composed solely of *AA* individuals and the susceptible population solely of *BB* individuals. As the epidemic progresses susceptible individuals become infected, and so the composition of the infected individuals becomes a mixture of *AA* and *BB*. If we ignore recoveries[5] and assume a large population, this process can be approximated by the following weighted average

$$\begin{aligned}\chi_{I,z}(t) &= \frac{\chi_{\text{seed},z}N_{\text{seed}}+\chi_{\text{cont},z}n}{N_{\text{seed}}+n} \\ &= \chi_{\text{cont},z}+\frac{N_{\text{seed}}}{N_{\text{seed}}+n}\left(\chi_{\text{seed},z}-\chi_{\text{cont},z}\right),\end{aligned} \tag{A20}$$

where $n$ is the number of infection events up to time $t$ (note this gives the correct limits: when $n=0$ so $\chi_{I,z}(0)=\chi_{\text{seed},z}$, and when $n\to\infty$ so $\chi_{I,z}(t)=\chi_{\text{cont},z}$). Substituting Eq.(A20) into (A19) gives

$$\begin{aligned}\frac{\partial^2\log(P)}{\partial a_g\partial a_f} &= -\sum_z\left[\phi N_{\text{cont}}\chi^2_{\text{cont},z}+q_z\chi_{\text{cont},z}\left(\chi_{\text{seed},z}-\chi_{\text{cont},z}\right)\right] \\ &\quad+\langle\chi_{\text{cont}}\rangle\left[\sum_z\left(\phi N_{\text{cont}}\chi_{\text{cont},z}+q_z\left(\chi_{\text{seed},z}-\chi_{\text{cont},z}\right)\right)\right], \\ \frac{\partial^2\log(P)}{\partial a_f^2} &= -\sum_z\left[\phi N_{\text{cont}}\chi^2_{\text{cont},z}+2q_z\chi_{\text{cont},z}\left(\chi_{\text{seed},z}-\chi_{\text{cont},z}\right)+w_z\left(\chi_{\text{seed},z}-\chi_{\text{cont},z}\right)^2\right] \\ &\quad+\frac{1}{N_I}\left[\sum_z\left(\phi N_{\text{cont}}\chi_{\text{cont},z}+q_z\left(\chi_{\text{seed},z}-\chi_{\text{cont},z}\right)\right)\right]^2.\end{aligned} \tag{A21}$$

---

[5] This is valid because most of the variation in $\chi_{I,z}(t)$ occurs at the beginning of the epidemic.

The sums used to define $q_z$ and $w_z$ in Eq.(A21) can be approximated by integrals:

$$q_z = \sum_{n=0}^{\phi N_{\text{cont}}} \frac{N_{\text{seed}}}{N_{\text{seed}} + n} \cong \int_0^{\phi N_{\text{cont}}} \frac{N_{\text{seed}}}{N_{\text{seed}} + n} dn = N_{\text{seed}} \log\left( \frac{N_{\text{seed}} + \phi N_{\text{cont}}}{N_{\text{seed}}} \right) = -N_{\text{seed}} \log(h),$$
$$w_z = \sum_{n=0}^{\phi N_{\text{cont}}} \left( \frac{N_{\text{seed}}}{N_{\text{seed}} + n} \right)^2 \cong \int_0^{\phi N_{\text{cont}}} \left( \frac{N_{\text{seed}}}{N_{\text{seed}} + n} \right)^2 dn = \frac{N_{\text{seed}} \phi N_{\text{cont}}}{N_{\text{seed}} + \phi N_{\text{cont}}} = N_{\text{seed}}(1-h), \qquad \text{(A22)}$$

where $h=N_{\text{seed}}/(N_{\text{seed}}+\phi N_{\text{cont}})$ gives the proportion of total infections that are accounted for by seeders.

The partial derivatives in Eqs.(A17) and (A21) can be re-expressed in the following way:

$$\frac{\partial^2 \log(P)}{\partial a_g^2} = -N_{\text{group}} \phi N_{\text{cont}} \left( \langle H_{\text{cont}} \rangle - \langle \chi_{\text{cont}} \rangle^2 \right),$$
$$\frac{\partial^2 \log(P)}{\partial a_g \partial a_f} = -N_{\text{group}} \phi N_{\text{cont}} \text{Var}(\chi_{\text{cont}}) - N_{\text{group}} N_{\text{seed}} W, \qquad \text{(A23)}$$
$$\frac{\partial^2 \log(P)}{\partial a_f^2} = -N_{\text{group}} \phi N_{\text{cont}} \text{Var}(\chi_{\text{cont}}) - N_{\text{group}} N_{\text{seed}} (2W + Y),$$

where $\text{Var}(\chi_{\text{cont}}) = \langle \chi_{\text{cont}}^2 \rangle - \langle \chi_{\text{cont}} \rangle^2$ is the variance in the homozygote balance across groups for the contact populations and

$$W = -\log(h) \langle (\chi_{\text{cont}} - \langle \chi_{\text{cont}} \rangle)(\chi_{\text{seed}} - \chi_{\text{cont}}) \rangle,$$
$$Y = (1-h) \left\langle (\chi_{\text{seed}} - \chi_{\text{cont}})^2 \right\rangle - \tfrac{N_{\text{seed}}}{\phi N_{\text{cont}}} \log^2(h) \langle \chi_{\text{seed}} - \chi_{\text{cont}} \rangle^2 . \qquad \text{(A24)}$$

Substituting the results in Eq.(A23) into the observed Fisher information matrix in Eq.(A7) leads to the matrix presented in Eq.(4).
