## Additional file 3 for "Optimal experimental designs for estimating genetic and non-genetic effects underlying infectious disease transmission"

### Inversion of the observed Fisher information matrix

The inverse of the Fisher information matrix is given by

$$\mathbf{M}^{-1} = \frac{1}{M_{gg}M_{ff} - M_{gf}^2}\begin{pmatrix} M_{ff} & -M_{gf} \\ -M_{gf} & M_{gg} \end{pmatrix}. \tag{A1}$$

The diagonals of this matrix give the inverse of the marginalised variances in $a_g$ and $a_f$:

$$\begin{aligned} \frac{1}{\mathrm{Var}(a_g)} &= M_{gg} - \frac{M_{gf}^2}{M_{ff}}, \\ \frac{1}{\mathrm{Var}(a_f)} &= M_{ff} - \frac{M_{gf}^2}{M_{gg}}. \end{aligned} \tag{A2}$$

Substituting in terms from the matrix in Eq.(4) gives

$$\begin{aligned} \frac{1}{\mathrm{Var}(a_g)} &= N_{\mathrm{group}}\phi N_{\mathrm{cont}}\left(\langle H_{\mathrm{cont}}\rangle - \langle \chi_{\mathrm{cont}}\rangle^2\right) - \frac{\left(N_{\mathrm{group}}\phi N_{\mathrm{cont}}\mathrm{Var}(\chi_{\mathrm{cont}}) + N_{\mathrm{group}}N_{\mathrm{seed}}W\right)^2}{N_{\mathrm{group}}\phi N_{\mathrm{cont}}\mathrm{Var}(\chi_{\mathrm{cont}}) + N_{\mathrm{group}}N_{\mathrm{seed}}(2W+Y)}, \\ \frac{1}{\mathrm{Var}(a_f)} &= N_{\mathrm{group}}\phi N_{\mathrm{cont}}\mathrm{Var}(\chi_{\mathrm{cont}}) + N_{\mathrm{group}}N_{\mathrm{seed}}(2W+Y) - \frac{\left(N_{\mathrm{group}}\phi N_{\mathrm{cont}}\mathrm{Var}(\chi_{\mathrm{cont}}) + N_{\mathrm{group}}N_{\mathrm{seed}}W\right)^2}{N_{\mathrm{group}}\phi N_{\mathrm{cont}}\left(\langle H_{\mathrm{cont}}\rangle - \langle \chi_{\mathrm{cont}}\rangle^2\right)}. \end{aligned}$$

(A3)

The first expression can be rearranged in the following way:

$$\begin{aligned} \frac{1}{\mathrm{Var}(a_g)} &= N_{\mathrm{group}}\phi N_{\mathrm{cont}}\left(\langle H_{\mathrm{cont}}\rangle - \langle \chi_{\mathrm{cont}}^2\rangle\right) + N_{\mathrm{group}}N_{\mathrm{seed}}\frac{\phi N_{\mathrm{cont}}\mathrm{Var}(\chi_{\mathrm{cont}})Y - N_{\mathrm{seed}}W^2}{\phi N_{\mathrm{cont}}\mathrm{Var}(\chi_{\mathrm{cont}}) + N_{\mathrm{seed}}(2W+Y)}, \\ &= N_{\mathrm{group}}\phi N_{\mathrm{cont}}\left(\langle H_{\mathrm{cont}}\rangle - \langle \chi_{\mathrm{cont}}^2\rangle\right) + N_{\mathrm{group}}N_{\mathrm{seed}}\left[Y - \frac{N_{\mathrm{seed}}(W+Y)^2}{\phi N_{\mathrm{cont}}\mathrm{Var}(\chi_{\mathrm{cont}}) + N_{\mathrm{seed}}(2W+Y)}\right], \end{aligned}$$

(A4)

as shown in Eq.(6). The expression for $\mathrm{Var}(a_f)$ from Eq.(A3) can be written

$$
\begin{aligned}
\frac{1}{\mathrm{Var}(a_f)} &= N_{\mathrm{group}} N_{\mathrm{seed}} Y + \\
&\frac{\left(N_{\mathrm{group}}\phi N_{\mathrm{cont}} \mathrm{Var}\left(\chi_{\mathrm{cont}}\right) + 2N_{\mathrm{group}} N_{\mathrm{seed}} W\right)\left(\left\langle H_{\mathrm{cont}}\right\rangle - \left\langle \chi_{\mathrm{cont}}\right\rangle^2\right) - \frac{\left(N_{\mathrm{group}}\phi N_{\mathrm{cont}} \mathrm{Var}\left(\chi_{\mathrm{cont}}\right) + N_{\mathrm{group}} N_{\mathrm{seed}} W\right)^2}{N_{\mathrm{group}}\phi N_{\mathrm{cont}}}}{\left\langle H_{\mathrm{cont}}\right\rangle - \left\langle \chi_{\mathrm{cont}}\right\rangle^2}, \\
&= N_{\mathrm{group}} N_{\mathrm{seed}} Y + \frac{\left(N_{\mathrm{group}}\phi N_{\mathrm{cont}} \mathrm{Var}\left(\chi_{\mathrm{cont}}\right) + 2N_{\mathrm{group}} N_{\mathrm{seed}} W\right)\left(\left\langle H_{\mathrm{cont}}\right\rangle - \left\langle \chi^2_{\mathrm{cont}}\right\rangle\right) - \frac{N^2_{\mathrm{seed}}}{\phi N_{\mathrm{cont}}} N_{\mathrm{group}} W^2}{\left\langle H_{\mathrm{cont}}\right\rangle - \left\langle \chi_{\mathrm{cont}}\right\rangle^2}, \\
&= N_{\mathrm{group}} N_{\mathrm{seed}} \left[ Y + \frac{2W\left(\left\langle H_{\mathrm{cont}}\right\rangle - \left\langle \chi^2_{\mathrm{cont}}\right\rangle\right) - \frac{N_{\mathrm{seed}}}{\phi N_{\mathrm{cont}}} W^2}{\left\langle H_{\mathrm{cont}}\right\rangle - \left\langle \chi_{\mathrm{cont}}\right\rangle^2} \right] + N_{\mathrm{group}}\phi N_{\mathrm{cont}} \frac{\left\langle H_{\mathrm{cont}}\right\rangle - \left\langle \chi^2_{\mathrm{cont}}\right\rangle}{\left\langle H_{\mathrm{cont}}\right\rangle - \left\langle \chi_{\mathrm{cont}}\right\rangle^2} \mathrm{Var}(\chi_{\mathrm{cont}}),
\end{aligned}
$$

(A5)

which is given in Eq.(7).
