## Additional file 4 for "Optimal experimental designs for estimating genetic and non-genetic effects underlying infectious disease transmission"

### Derivation of standard deviations for recoverability SNP effect

Here we derive the standard deviation (SD) for the SNP parameter related to recoverability $a_r$. As in the analysis for the infection process, we ignore fixed effects and residual contributions in Eq.(2). Consequently the fractional deviation in recoverability for individual $m$ is simply given by the SNP effect, *i.e.* $r_m = r_m^{\text{SNP}}$.

In Additional file 2 Eq.(A1), the gamma distributed probability density function for $\delta t_m$ (*i.e.* the infection duration for individual $m$) is given by

$$F_\Gamma(\delta t_m \mid w_m, k) = \Gamma(k)^{-1} w_m^{-k} k^k \delta t_m^{k-1} e^{-k\delta t_m / w_m}, \tag{A1}$$

where $k$ is a shape parameter that determines the recovery profile and $w_m$ is the individual-based mean infection duration given in Eq.(1). Explicitly substituting this into Eq.(A1), and then placing this into the likelihood in Additional file 2 Eq.(A1) yields the distribution

$$\pi(\gamma, a_r, \Delta_r \mid \xi, k) \propto \gamma^{kN_R} e^{-\gamma K} \prod_z \left[\prod_{m\in z} e^{kr_m^{\text{SNP}}}\right] \quad \text{where } K = \sum_z \left[\sum_{m\in z} e^{r_m^{\text{SNP}}} k\delta t_m\right], \tag{A2}$$

where $N_R$ is the total number of individuals which recover. This can be integrated w.r.t. $\gamma$, leaving

$$\begin{aligned} \pi(a_r, \Delta_r \mid \xi, k) &\propto \int_0^\infty \gamma^{kN_R} e^{-\gamma K} \mathrm{d}\gamma \prod_z \left[\prod_{m\in z} e^{kr_m^{\text{SNP}}}\right] \\ &\propto [kN_R]! K^{-kN_R-1} \prod_z \left[\prod_{m\in z} e^{kr_m^{\text{SNP}}}\right]. \end{aligned} \tag{A3}$$

To estimate the standard deviation in $a_r$ we assume the case of no dominance such that

$$r_m^{\text{SNP}} = \kappa_m a_r, \tag{A4}$$

where $\kappa_m$={1,0,-1} depending on whether the genotype of individual $m$ is {*AA*,*AB*,*BB*}. Taking the double derivative of the log of Eq.(A3) w.r.t. $a_r$ gives

$$\frac{\partial^2 \log\left[\pi(a_r \mid \xi, \Delta_r, k)\right]}{\partial a_r^2} \cong -kN_R \left[\frac{1}{K}\sum_z \left[\sum_{m\in z} \kappa_m^2 e^{r_m^{\text{SNP}}} k\delta t_m\right] - \left(\frac{1}{K}\sum_z \left[\sum_{m\in z} \kappa_m e^{r_m^{\text{SNP}}} k\delta t_m\right]\right)^2\right]. \tag{A5}$$

Following the approximations used in Additional file 2 Eq.(A11), this becomes

$$\frac{\partial^2 \log\left[\pi(a_r \mid \xi, \Delta_r, k)\right]}{\partial a_r^2} \cong -kN_R \left(\overline{H} - \overline{\chi}^2\right), \tag{A6}$$

where

$$\overline{H} = \frac{N_{\text{seed}}\langle H_{\text{seed}}\rangle + \phi N_{\text{cont}}\langle H_{\text{cont}}\rangle}{N_{\text{seed}} + \phi N_{\text{cont}}}, \quad \overline{\chi} = \frac{N_{\text{seed}}\langle \chi_{\text{seed}}\rangle + \phi N_{\text{cont}}\langle \chi_{\text{cont}}\rangle}{N_{\text{seed}} + \phi N_{\text{cont}}} \tag{A7}$$

represent the total homozygosity and homozygote balance for the entire infected population (*i.e.* including seeders as well as contacts infected during the experiment), respectively.

Since $N_{\mathrm{seed}}$ individuals are initially infected in each contact group and on average $\phi N_{\mathrm{cont}}$ become infected, the total number of recoveries is given by

$$N_R = N_{\mathrm{group}}\left(N_{\mathrm{seed}} + \phi N_{\mathrm{cont}}\right). \tag{A8}$$

Substituting this into Eq.(A6) gives

$$\frac{\partial^2 \log\left[\pi(a_r \mid \xi, \Delta_r, k)\right]}{\partial a_r^2} \cong -kN_{\mathrm{group}}\left(N_{\mathrm{seed}} + \phi N_{\mathrm{cont}}\right)\left(\overline{H} - \overline{\chi}^2\right). \tag{A9}$$

In the case of recoverability, the observed Fisher information matrix is a scalar quantity, and so the standard deviation is simply given by

$$\text{SD in } a_r = \left(-\frac{\partial^2 \log\left[\pi(a_r \mid \xi, \Delta_r, k)\right]}{\partial a_r^2}\right)^{-\frac{1}{2}} = \frac{1}{\sqrt{kN_{\mathrm{group}}\left(N_{\mathrm{seed}} + \phi N_{\mathrm{cont}}\right)\left(\overline{H} - \overline{\chi}^2\right)}}. \tag{A10}$$
