## Additional file 5 for "Optimal experimental designs for estimating genetic and non-genetic effects underlying infectious disease transmission"

### Dominance

#### 1) Susceptibility

Here we derive the standard deviation (SD) in the susceptibility dominance parameter $\Delta_g$ assuming a known SNP effect size $a_g$. For clarity the derivation below assumes $a_f$=*0*, but in fact it is generally true provided $a_f$ is small (which anyway is an assumption already used for much of the analysis). Under this circumstance the distribution from Additional file 2 Eq.(A5) is given by

$$\pi(\Delta_g \mid \xi, a_g, a_f, \Delta_f) \propto N_I! J^{-N_I-1} \prod_z \left[ \prod_{j\in z} \left( e^{g_j^{\mathrm{SNP}}} I_j \right) \right], \quad \text{(A1)}$$
$$J = \sum_z \left( \sum_{e\in z} \left[ \left( \sum_{s\in e} e^{g_s^{\mathrm{SNP}}} \right) I_e (t_e - t_{e-1}) \right] \right),$$

where $N_I$ is the total number of infections during the experiment, $z$ goes over all contact groups and in each group $j$ goes over individuals that get infected, $e$ goes over events (infections and recoveries), and $s$ goes over all susceptible individuals immediately prior to event $e$. The infected population size $I_j$ is evaluated prior to individual $j$ being infected and $I_e$ is the corresponding quantity prior to the time of event $e$.

The log of Eq.(A1) is (up to a constant and assuming $N_I \gg 1$)

$$\log(P) = -N_I \log(J) + \sum_z \left[ \sum_{j\in z} g_j^{\mathrm{SNP}} \right], \quad \text{(A2)}$$

where $P = \pi(\Delta_g|\xi, a_g, a_f, \Delta_f)$. Using the expression for $g_j^{\mathrm{SNP}}$ in Eq.(3) and taking the double derivative of Eq.(A2) w.r.t. $\Delta_g$ gives

$$\frac{\partial^2 \log(P)}{\partial \Delta_g^2} = -\frac{N_I}{J} a_g^2 \left( \sum_z \left( \sum_{e\in z} \left[ \left( \sum_{s\in e} \tau_s^2 e^{g_s^{\mathrm{SNP}}} \right) I_e (t_e - t_{e-1}) \right] \right) \right) \quad \text{(A3)}$$
$$+ \frac{N_I}{J^2} a_g^2 \left( \sum_z \left( \sum_{e\in z} \left[ \left( \sum_{s\in e} \tau_s e^{g_s^{\mathrm{SNP}}} \right) I_e (t_e - t_{e-1}) \right] \right) \right)^2,$$

where $\tau_s$={0,1,0} depends on whether individual $s$ has genotype {*AA*,*AB*,*BB*}. The following approximations can be made (which are valid for small $a_g$)

$$\sum_{s\in e} \tau_s e^{g_s^{SNP}} \cong (1 - H_{\mathrm{cont},z}) S_e, \quad \text{(A4)}$$
$$\sum_{s\in e} \tau_s^2 e^{g_s^{SNP}} \cong (1 - H_{\mathrm{cont},z}) S_e,$$
$$\sum_{s\in e} e^{g_s^{SNP}} \cong S_e,$$

where $H_{\mathrm{cont},z}$ is the homozygosity (*i.e.* proportion of *AA* and *BB* individuals) of the contacts in group $z$. Using this, along with the expression from Additional file 2 Eq.(A15), finally gives

$$\frac{\partial^2 \log(P)}{\partial \Delta_g^2} = a_g^2 \left[ -\phi N_{\text{cont}} \left( \sum_z (1 - H_{\text{cont},z}) \right) + \frac{\phi N_{\text{cont}}}{N_{\text{group}}} \left( \sum_z (1 - H_{\text{cont},z}) \right)^2 \right]$$
$$= -N_{\text{group}} \phi N_{\text{cont}} a_g^2 \left[ \langle H_{\text{cont}} \rangle - \langle H_{\text{cont}} \rangle^2 \right]. \qquad \text{(A5)}$$

The observed Fisher information matrix is a scalar quantity, and so the standard deviation is simply given by one over the square root of minus this quantity, as shown in Eq.(10).

### 2) Infectivity

Here we derive the SD in the susceptibility dominance parameter $\Delta_f$ assuming a known SNP effect size $a_f$. Again, for clarity the derivation below assumes $a_g$=0, but in fact it is generally true provided $a_g$ is small. The distribution from Additional file 2 Eq.(A5) is given by

$$\pi(\Delta_f \mid \xi, a_g, a_f, \Delta_g) \propto N_I! J^{-N_I-1} \prod_z \left[ \prod_{j\in z} \left( \sum_{i\in j} e^{f_i^{\text{SNP}}} \right) \right],$$
$$J = \sum_z \left( \sum_{e\in z} \left[ S_e \left( \sum_{i\in e} e^{f_i^{\text{SNP}}} \right) (t_e - t_{e-1}) \right] \right). \qquad \text{(A6)}$$

Taking the log of this expression gives

$$\log(P) = -N_I \log(J) + \sum_z \left[ \sum_{j\in z} \left( \sum_{i\in j} e^{f_i^{\text{SNP}}} \right) \right], \qquad \text{(A7)}$$

where $P=\pi(\Delta_f|\xi,a_g,a_f,\Delta_g)$. Next we make use of the following approximations (valid in the limit of small $a_f$ and taking the continuum limit, see Eqs.(A4) and Additional file Eq.(A18) for previously derived analogous results):

$$\sum_{i\in e} \tau_i e^{f_i^{\text{SNP}}} \cong (1 - H_{I,z}(t)) I_e,$$
$$\sum_{i\in e} \tau_i^2 e^{f_i^{\text{SNP}}} \cong (1 - H_{I,z}(t)) I_e,$$
$$\sum_{i\in e} e^{f_i^{\text{SNP}}} \cong I_e,$$
$$\sum_{e\in z} \left[ (1 - H_{I,z}(t)) S_e I_e (t_e - t_{e-1}) \right] \cong \tfrac{1}{\beta} \sum_{j\in z} (1 - H_{I,z}(t)), \qquad \text{(A8)}$$

where $I_e$ is the number of infected individuals immediately prior to event $e$ and $H_{I,z}(t) = \nu_{AA,z}^I(t) + \nu_{BB,z}^I(t)$ is the time varying homozygosity in the infected population in group $z$. Taking the double derivative of Eq.(A7) w.r.t. $\Delta_f$ and using the results in Eq.(A8) leads to

$$\frac{\partial^2 \log(P)}{\partial \Delta_f^2} = a_f^2 \left[ -\sum_z \left[ \sum_{j\in z} (1 - H_{I,z}(t))^2 \right] + \frac{1}{N_I} \left[ \sum_z \left( \sum_{j\in z} (1 - H_{I,z}(t)) \right) \right]^2 \right]. \qquad \text{(A9)}$$

As in Additional file 2 Eq.(A20), we model this time variation using

$$H_{I,z}(t) = H_{\text{cont},z} + \frac{N_{\text{seed}}}{N_{\text{seed}} + n} \left( H_{\text{seed},z} - H_{\text{cont},z} \right), \qquad \text{(A10)}$$

where $n$ is the number of infections that have occur up to time $t$. Substituting this into Eq.(A9) and making use of the definitions in Additional file 2 Eq.(A22) finally gives

$$\frac{\partial^2 \log(P)}{\partial \Delta_f^2} = -a_f^2 \left[ N_{\text{group}} \phi N_{\text{cont}} \text{Var}\left(H_{\text{cont}}\right) + N_{\text{group}} N_{\text{seed}} \left(2W_H + Y_H\right) \right], \qquad \text{(A11)}$$

where

$$\begin{aligned} W_H &= -\log(h)\left\langle (H_{\text{cont}} - \langle H_{\text{cont}} \rangle)(H_{\text{seed}} - H_{\text{cont}}) \right\rangle, \\ Y_H &= (1-h)\left\langle \left(H_{\text{seed}} - H_{\text{cont}}\right)^2 \right\rangle - \tfrac{N_{\text{seed}}}{\phi N_{\text{cont}}} \log^2(h) \left\langle H_{\text{seed}} - H_{\text{cont}} \right\rangle^2. \end{aligned} \qquad \text{(A12)}$$

The standard deviation in $\Delta_f$ is given by one over the square root of minus this quantity, as shown in Eq.(11).

#### 3) Recoverability

Assuming the effect size $a_r$ is known, taking the double derivative of the log of Additional file 4 Eq.(A3) w.r.t. $\Delta_r$ leads to (following the same basic approach as above)

$$\frac{\partial^2 \log(P)}{\partial \Delta_r^2} = -a_r^2 k N_{\text{group}} \left(N_{\text{seed}} + \phi N_{\text{cont}}\right)\left(\overline{H} - \overline{H}^2\right), \qquad \text{(A13)}$$

where the average homozygosity $\overline{H}$ is defined in Eq.(9). This gives the final result in Eq.(13).
