## Additional file 6 for "Optimal experimental designs for estimating genetic and non-genetic effects underlying infectious disease transmission"

### Epidemic extinction

Figure S1 shows how the probability of epidemic extinction varies as a function of the number of seeder individuals $N_{\text{seed}}$ for different basic reproductive ratios $R_0$. This graph was generated by performing a large number of epidemic simulations (using the Doob-Gillespie algorithm [36] described in Additional file 1) that were initialised with a given infected seeder population $N_{\text{seed}}$ and an assumed large (arbitrary) uninfected contact population $N_{\text{cont}}$. The fraction of simulations in which the epidemic did not proliferate gave the extinction probability.

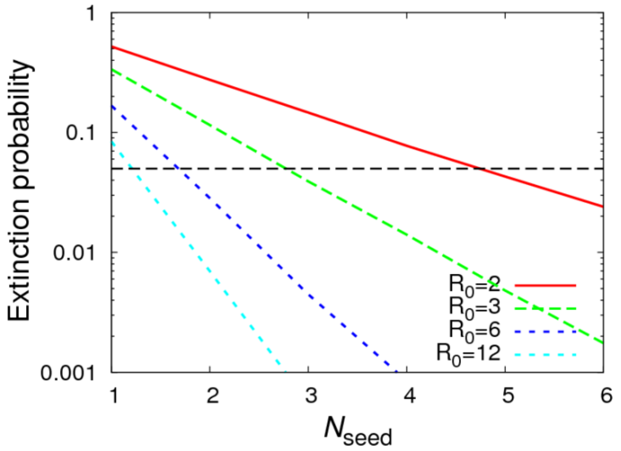


**Figure S1.** *Epidemic extinction.* Shows the probability that an epidemic undergoes an extinction (*i.e.* the infected seeders do not lead to an epidemic) as a function of the number of seeders. The curves represent different basic reproductive ratios $R_0$. The horizontal dashed line represents an arbitrary 5% cut-off.

From an experimental point of view, having a large proportion of contact groups not undergoing epidemics is problematic because it substantially reduces the statistical power with which to perform inference (many individuals don't even become exposed, so can provide no information regarding their susceptibility, infectivity or recoverability). This paper assumes $R_0$=3, hence Fig. S1 suggests that at least 3 seeders are required to ensure that the probability of epidemic extinction is less than 5%. As $R_0$ reduces closer to one, more and more seeders are needed to avoid substantial extinction probability.
