## Additional file 7 for "Optimal experimental designs for estimating genetic and non-genetic effects underlying infectious disease transmission"

### Further details for the single contact group design

Under the single contact group design in Fig. 2a, the SDs in the SNP effects are given by:

$$\begin{aligned}
&\text{SD in } a_g \cong \frac{1}{\sqrt{N_{\text{total}}\left(1-h\right)}}, \\
&\text{SD in } a_f \cong \frac{1}{\sqrt{N_{\text{total}}\left(h(1-h)-\dfrac{h^2}{1-h}\log^2\left(h\right)\right)\left(\chi_{\text{seed}}-\chi_{\text{cont}}\right)^2}}, \\
&\text{SD in } a_r = \frac{1}{\sqrt{kN_{\text{total}}}},
\end{aligned} \qquad \text{(A1)}$$

where $h=N_{\text{seed}}/G_{\text{size}}$ is the fraction of seeder individuals (here contacts are all assumed to become infected, *i.e.* $\phi\approx1$, appropriate for reasonably large $R_0$).

Since this design contains no heterozygote *AB* individuals, no information is available for estimating the dominance parameters. The expressions reported in Table 2 come from substituting the optimal design values from Fig. 2a into Eq.(A1).
