## Additional file 8 for "Optimal experimental designs for estimating genetic and non-genetic effects underlying infectious disease transmission"

### Impact of changing $R_0$

As the basic reproductive ratio $R_0$ becomes closer to one, so the fraction of contact individuals that become infected during experiments is expected to go down. This additional file numerically investigates how this affects the optimal design choices outlined in Fig. 2.

For each experimental design different values of $R_0$ were chosen (by varying $\beta$ and fixing $\gamma$). A large number of simulations were performed ($10^5$) to establish the average fraction of contacts $\phi$ which become infected. The SD in $a_f$ was estimated using the analytical expression from Eq.(7). The procedure was repeated whilst varying critical design parameters (*e.g.* the fraction of individual which are seeders in the case of the single contact group). The parameter value that minimised the SD in $a_f$ is plotted in Fig. S1 (the five sub-plots correspond to the designs in Fig. 2). We observe that all the optimal designs are largely insensitive to the exact value of $R_0$.

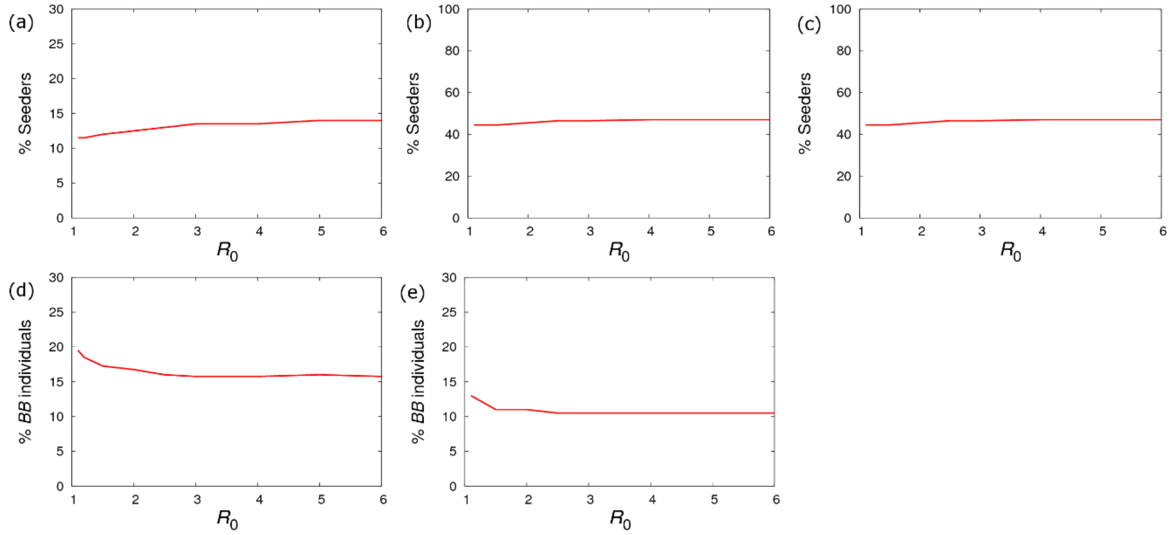


**Figure S1.** *Optimum design as a function of $R_0$.* Shows how values for key parameters that optimise the precision of $a_f$ (as calculated using Eq.(7)) vary as a function of $R_0$ for each of the experimental designs in Fig. 2. (a) Single contact group (without dominance) % of seeders. (b) Pure design (without dominance) % of seeders. (c) Pure design (with dominance) % of seeders. (d) Mixed design (without dominance) % of *BB* in group 1 (which is equal to the % of *AA* in group 2). (e) Mixed design (with dominance) % of *BB* in group 1 (which equals the % of *AB*). Other groups have the same proportions but with genotypes permuted.
