## Additional file 9 for "Optimal experimental designs for estimating genetic and non-genetic effects underlying infectious disease transmission"

### Further details for the "pure" design

Here we provide additional information about the pure design. For the four group design in Fig. 2b (without dominance), the SDs in SNP effects are given by

$$\text{SD in } a_g \cong \frac{1}{\sqrt{N_{\text{total}}\left(1-h-\dfrac{[1-h+h\log(h)]^2}{1+h-2h^2+2h\log(h)}\right)}},$$
$$\text{SD in } a_f \cong \frac{1}{\sqrt{N_{\text{total}}\left(2h(1-h)-\dfrac{h^2}{1-h}\log^2(h)\right)}}, \quad \text{(A1)}$$
$$\text{SD in } a_r = \frac{1}{\sqrt{kN_{\text{total}}}},$$

where $h$=$N_{\text{seed}}/G_{\text{size}}$ is the fraction of seeder individuals (here contacts are all assumed to become infected, *i.e.* $\phi$≈1, appropriate for reasonably large $R_0$). Since this design contains no heterozygote *AB* individuals, no information is available for estimating the dominance parameters.

For the nine group design in Fig. 2b (with dominance), the SDs in SNP effects are given by √(3/2)=1.2 times the expressions in Eq.(A1). Estimates for the SD in the dominance parameters are given by

$$\text{SD in } \Delta_g \cong \frac{1}{|a_g|\sqrt{\frac{2}{9}N_{\text{total}}(1-h)}},$$
$$\text{SD in } \Delta_f \cong \frac{1}{|a_f|\sqrt{\frac{2}{9}N_{\text{total}}\left(1+h-2h^2+2h\log(h)\right)}}, \quad \text{(A2)}$$
$$\text{SD in } \Delta_r = \frac{1}{|a_r|\sqrt{\frac{2}{9}kN_{\text{total}}}}.$$

Note the expressions in Eq.(A2) show that in order to determine information regarding dominance it is necessary for the SNP effect itself to have a non-zero value. The left-hand graph in Fig. S1a shows how the SD in $\Delta_g$ varies as a function of $a_g$. We find, as expected, that the larger $a_g$, the greater the precision with which dominance can be identified. Comparing Fig. S1b to Fig. 4a, we observe that even though the pure design with dominance contains significantly fewer homozygotes, the estimates for the SDs are only marginally increased.

The expressions reported in Table 2 come from substituting the optimal design values from Fig. 2b into Eqs.(A1) and (A2).

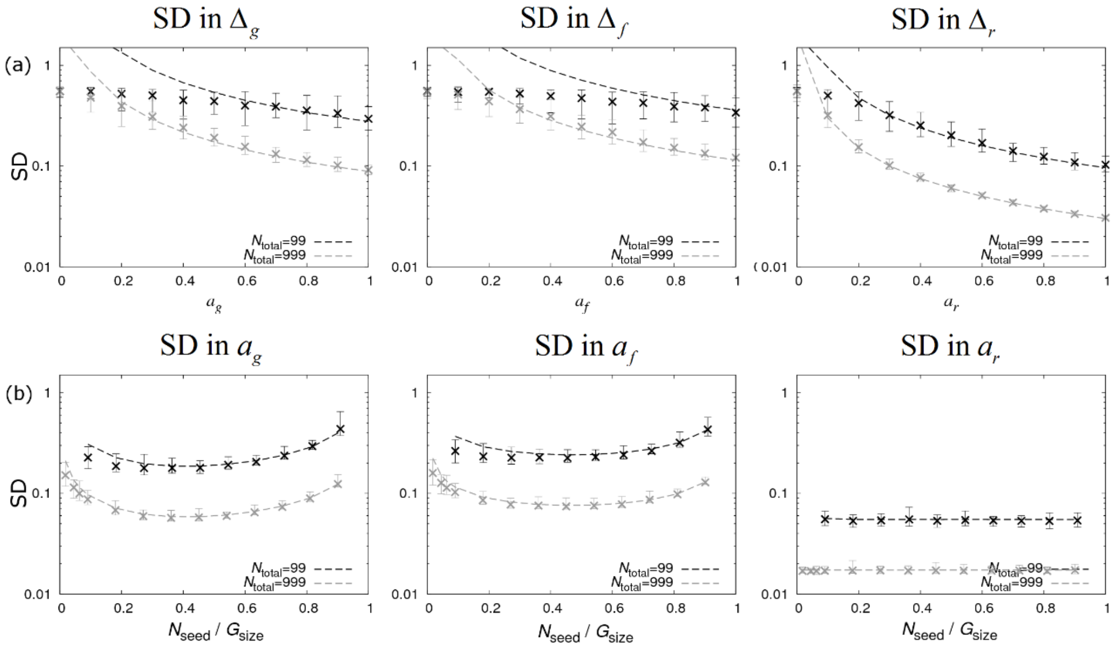


**Figure S1.** *Precision estimates for the pure design (with dominance) in Fig. 2b.* (a) The left, middle and right graphs show standard deviations in scaled dominance parameters for susceptibility $\Delta_g$, infectivity $\Delta_f$ and recoverability $\Delta_r$ as a function of the corresponding SNP effect sizes (SNP effects and dominance factors on the other traits are zero). (b) The left, middle and right graphs show standard deviations in the SNP effects for susceptibility $a_g$, infectivity $a_f$ and recoverability $a_r$ as a function of the fraction of individuals which are seeders (with zero dominance). Dashed lines represent analytical results and crosses come from posterior estimates from simulated data (see Additional file 1). $N_{\text{total}}$ refers to the total number of individuals.
