## Additional file 10 for "Optimal experimental designs for estimating genetic and non-genetic effects underlying infectious disease transmission"

### Further details for the "mixed" design

Here we provide additional information about the mixed design. For the two group design in Fig. 2c (without dominance), the SDs in SNP effects are given by

$$\begin{aligned} \text{SD in } a_g &\cong \frac{1}{\sqrt{N_{\text{total}}\left(1-\chi^2_{\text{cont},1}\right)}}, \\ \text{SD in } a_f &\cong \frac{1}{\sqrt{N_{\text{total}}\left(1-\chi^2_{\text{cont},1}\right)\chi^2_{\text{cont},1}}}, \\ \text{SD in } a_r &= \frac{1}{\sqrt{kN_{\text{total}}}}, \end{aligned} \qquad \text{(A1)}$$

where it is assumed that $N_{\text{seed}}$ is small compared to $G_{\text{size}}$ and all contacts become infected, *i.e.* $\phi$=1 (appropriate for reasonably large $R_0$). Since this design contains no heterozygote *AB* individuals, no information is available for estimating dominance parameters.

For the three group design in Fig. 2c (with dominance), the SDs in the SNP effects are given by √(3/2)=1.2 times the expressions in Eq.(A1). Estimates for the SDs in the dominance parameters are given by

$$\begin{aligned} \text{SD in } \Delta_g &\cong \frac{1}{|a_g|\sqrt{\tfrac{2}{9}N_{\text{total}}}}, \\ \text{SD in } \Delta_f &\cong \frac{1}{|a_f|\sqrt{\tfrac{2}{9}N_{\text{total}}\chi^2_{\text{cont},1}}}, \\ \text{SD in } \Delta_r &= \frac{1}{|a_r|\sqrt{\tfrac{2}{9}kN_{\text{total}}}}. \end{aligned} \qquad \text{(A2)}$$

Results comparing these analytical expressions with numerical posterior estimates are presented in Fig. S1. As with the pure results from Additional file 9 Fig. S1a, we find that the precision of dominance estimation increases with effect size, and as with the mixed results in Fig. 5a, the SD in $a_f$ is minimised when the homozygote balance is given by $\chi_{\text{cont},1}$=- $\chi_{\text{cont},2}$=1/√2 =0.71.

The expressions reported in Table 2 come from substituting the optimal design values from Fig. 2c into Eqs.(A1) and (A2).

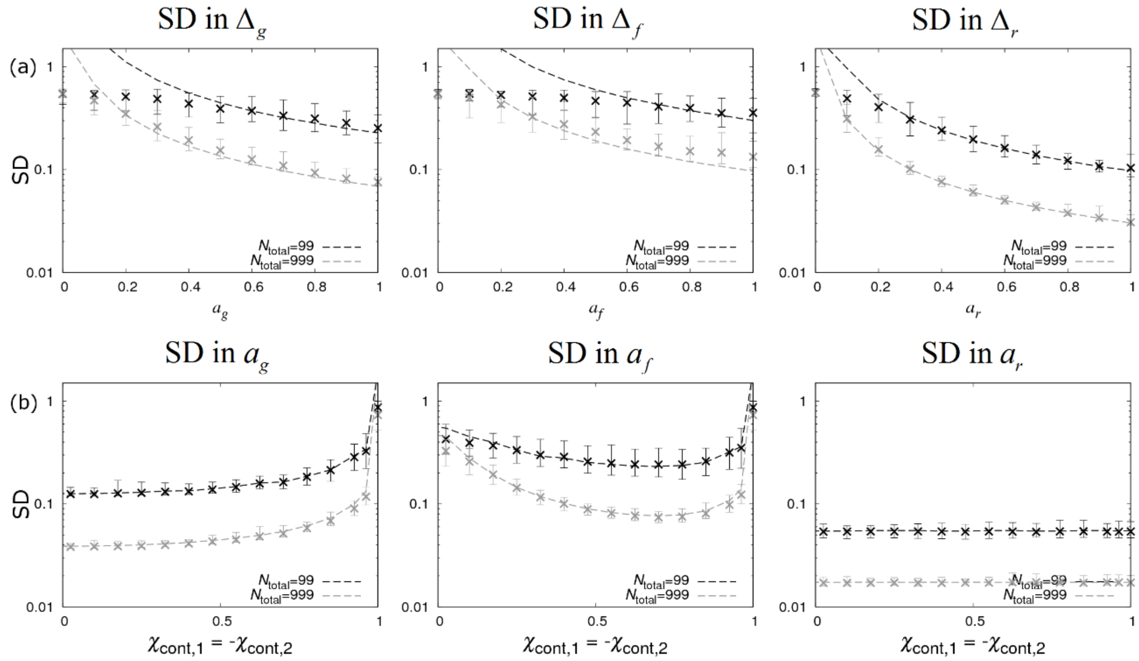


**Figure S1.** *Precision estimates for the mixed design (with dominance) in Fig. 2c.* (a) The left, middle and right graphs show standard deviations in scaled dominance parameters for susceptibility $\Delta_g$, infectivity $\Delta_f$ and recoverability $\Delta_r$ as a function of the corresponding SNP effect sizes (SNP effects and dominance factors on other traits are zero and $\chi_{cont,1}$=- $\chi_{cont,2}=1/\sqrt{2}$). (b) The left, middle and right graphs show standard deviations in the SNP effects for susceptibility $a_g$, infectivity $a_f$ and recoverability $a_r$ as a function of $\chi_{cont,1}$ (where the sizes of the two minority genotypes are taken to be the same, *e.g.* the number of *AB* and *BB* individuals in group 1 of Fig. 2c (with dominance) is the same). Dashed lines represent analytical results and crosses come from posterior estimates from simulated datasets (see Additional file 1). $N_{seed}$ =3 is used in all cases and $N_{total}$ refers to the total number of individuals.
