## Additional file 11 for "Optimal experimental designs for estimating genetic and non-genetic effects underlying infectious disease transmission"

### Impact of realistic model/data scenarios on design

Considering the optimal designs in Fig. 2, this additional file investigates the impact of separately introducing five additional sources of variation into the model/data (which, for the purposes of analysis, were ignored):

**Residual contributions** – The residual contributions $\boldsymbol{\varepsilon}_g$, $\boldsymbol{\varepsilon}_f$, and $\boldsymbol{\varepsilon}_r$ in Eq.(2) (with covariance matrix arbitrarily chosen to be $\Sigma_{gg}=\Sigma_{ff}=\Sigma_{rr}=1$, $\Sigma_{gf}=0.3$, $\Sigma_{gr}=-0.4$, and $\Sigma_{fr}=-0.2$) are now included in both the simulated data and inference. These account for other sources of variation in the traits, *e.g.* environmental and genetic contributions not coming from the SNP under analysis. The resulting estimated SDs in the infectivity SNP effect $a_f$ under the single contact group, pure and mixed designs (without dominance) are shown in Fig. S1a. Note, the crosses in all these graphs lie higher than in the corresponding graphs for the simplified model (see the middle columns in Figs. 3a, 4a and 5a for reference) reflecting the additional level of uncertainty in the infection dynamics. Interestingly, as discussed in the section below, most of this difference comes from simply incorporating residuals into the model, with only a relatively small contribution actually depending on their magnitude. The left hand graph in Fig. S1a shows that residuals are found to reduce the statistical power coming from the single contact group the most, with the optimum SDs for the pure and mixed designs exhibiting a similar increase compared to the analytical dashed curves. Importantly, the minima in both the crosses and the analytical curves occur at approximately the same position, meaning that the optimum designs in Fig. 2 (which require ~15% of seeders for the single group design, ~47% of seeders for the pure design and 83%17% split in *AA*/*BB* in the mixed design) still remain valid. A similar picture is seen for the designs in Fig. 2 with dominance (results not shown), and also largely remains true for the other sources of variation considered below.

**Group effect** – Random group effects in Eq.(1) account for group-specific factors that influence the overall speed of an epidemic in one contact group relative to another (*e.g.* animals kept in different management conditions or environmental differences). Here we introduce a group effect with standard deviation $\sigma_G=0.2$, which corresponds to a random ≈20% variation in the disease transmission rate across groups. Whilst the addition of this makes little difference to the single contact group design in Fig. S1b, the statistical power to infer infectivity SNP effects in the pure and mixed designs is substantially reduced (especially for the mixed design). The reason lies in the fact that almost all information regarding infectivity differences comes from the relative rate of epidemics in different contact groups. When a group effect is added such variation can equally be explained in terms of variation in $G_z$ leading to confounding. To overcome this difficulty two approaches can be taken: Firstly, if it is truly believed that no significant uncontrolled factors affect disease transmission then group effects can justifiably be left out of the model. Secondly, if group effects are considered necessary, their influence can markedly be reduced by increasing the number of design replicates, as demonstrated in the section below.

**Fixed effect** – As an example we here consider the case in which there is a sex difference in the traits (with fixed effect size $b_{g0}=b_{f0}=b_{r0}=0.2$) with individuals randomly allocated to be male or female. Since Fig. S1c exhibits almost no difference compared to the basic SNP model, it can be concluded that fixed effects have very little impact on SNP effect precisions. However, as discussed in Additional file 14, there can be a reduction in statistical power when there are significant correlations between the SNP and the fixed effect (leading to confounding). Therefore, when designing experiments care should be taken to randomise (across groups) factors other than the genotype of the SNP under investigation.

**Unknown infection times** – Figure S1d shows the case in which infection times are unknown and so must themselves be inferred from the available recovery/death data. The pattern here is similar to when residuals were introduced in Fig. S1a, with the single contact group design suffering the most and the pure and mixed designs suffering a relatively moderate reduction in statistical power.

**Periodic disease status checks** – Under this scenario rather than measuring infection and recovery times themselves, individuals are periodically checked for their disease status (every 3 time units from the start of the experiment). The results from Fig. S1e demonstrate that this leads to only a small loss in precision for SNP effect estimates in this particular example (indeed in [15] it is shown that a substantial reduction in precision only occurs when the period of measurements becomes similar to the overall timescale of epidemics themselves).

For completeness, results for the SDs in susceptibility $a_g$ and recoverability $a_r$ SNP effects are shown in Figs. S2 and S3.

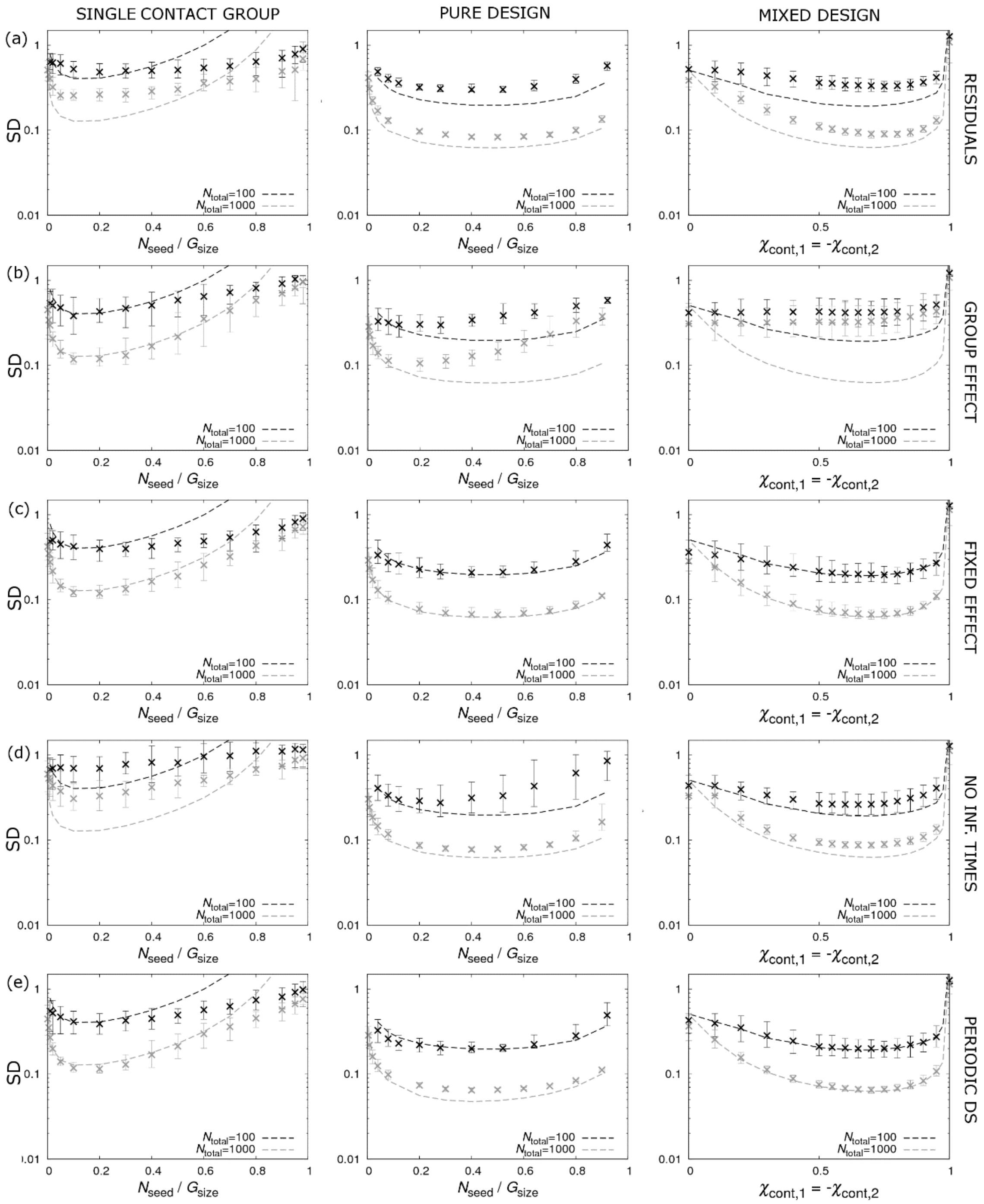


**Figure S1.** *Impact of sources of model/data variation on optimal design.* Shows key graphs (where the *y*-axis represents SDs in the infectivity SNP effect $a_f$) which can be used to identify the optimal percentages in Fig. 2 when various sources of variation are (individually) introduced into the model: (a) residual contributions ($\Sigma_{gg}$=$\Sigma_{ff}$=$\Sigma_{rr}$=1, $\Sigma_{gf}$=0.3, $\Sigma_{gr}$=-0.4, and $\Sigma_{fr}$=-0.2), (b) group effect ($\sigma_G$=0.2), (c) a fixed effect (for sex with individuals randomly allocated to male or female and effect fixed effect size $b_{g0}$=$b_{f0}$=$b_{r0}$=0.2), (d) unknown infection times and (e) periodic measurement of disease status (made every Δt=3 time units from the start of the experiments). Results in black represent a total of 1000 individuals and those in grey 100 individuals.

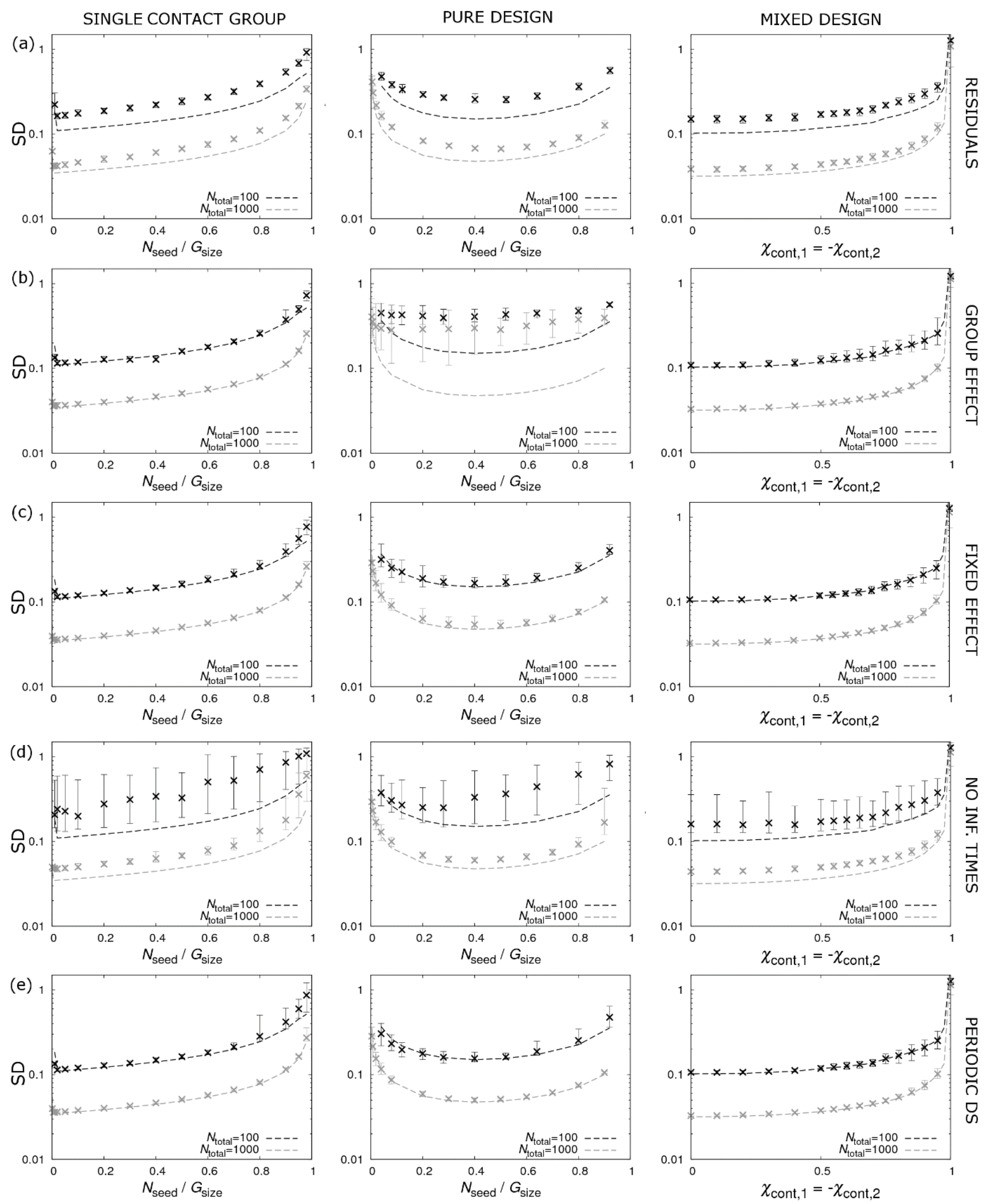


**Figure S2** *Impact of sources of model/data variation on optimal design.* Shows key graphs (where the *y*-axis represents SDs in the susceptibility SNP effect $a_g$) which can be used to identify the optimal percentages in Fig. 2 when various sources of variation are (individually) introduced into the model: (a) residual contributions ($\Sigma_{gg}$=$\Sigma_{ff}$=$\Sigma_{rr}$=1, $\Sigma_{gf}$=0.3, $\Sigma_{gr}$=-0.4, and $\Sigma_{fr}$=-0.2), (b) group effect ($\sigma_G$=0.2), (c) a fixed effect (for sex with individuals randomly allocated to male or female and effect fixed effect size $b_{g0}$=$b_{f0}$=$b_{r0}$=0.2), (d) unknown infection times and (e) periodic measurement of disease status (made every Δt=3 time units from the start of the experiments). Results in black represent a total of 1000 individuals and those in grey 100 individuals.

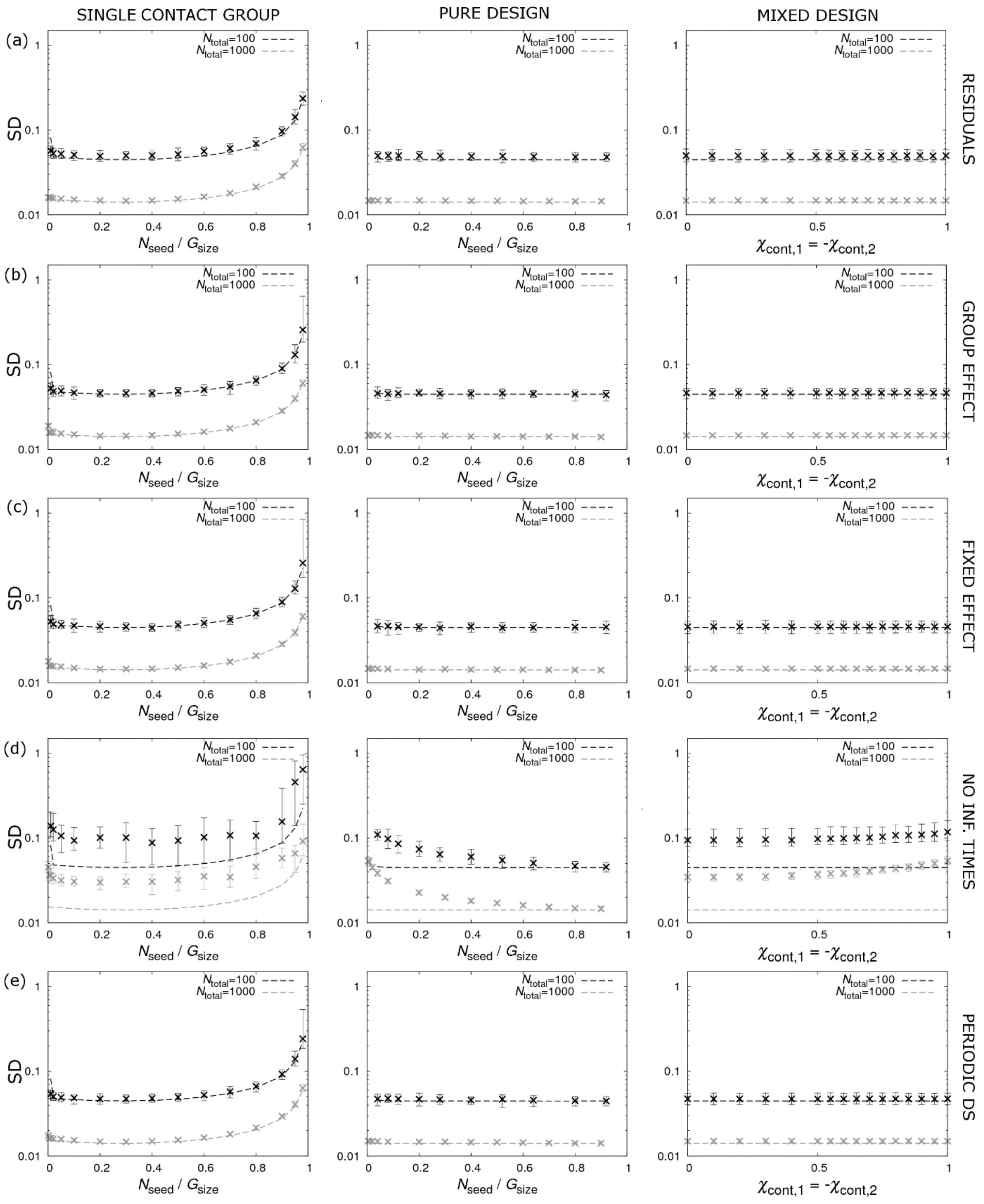


**Figure S3.** *Impact of sources of model/data variation on optimal design.* Shows key graphs (where the $y$-axis represents SDs in the recoverability SNP effect $a_r$) which can be used to identify the optimal percentages in Fig. 2 when various sources of variation are (individually) introduced into the model: (a) residual contributions ($\Sigma_{gg}=\Sigma_{ff}=\Sigma_{rr}=1$, $\Sigma_{gf}=0.3$, $\Sigma_{gr}=-0.4$, and $\Sigma_{fr}=-0.2$), (b) group effect ($\sigma_G=0.2$), (c) a fixed effect (for sex with individuals randomly allocated to male or female and effect fixed effect size $b_{g0}=b_{f0}=b_{r0}=0.2$), (d) unknown infection times and (e) periodic measurement of disease status (made every $\Delta t=3$ time units from the start of the experiments). Results in black represent a total of 1000 individuals and those in grey 100 individuals.

### Effect of residuals on precision of SNP effects

This additional file introduces residuals into the basic SNP model and investigates how they reduce the precision with which SNP effects can be estimated under the pure and mixed designs. Specifically, Fig. S4 shows how the SDs in $a_g$, $a_f$ and $a_r$ increase as a function of a control parameter $u$, which determines the size of the residual contributions.

In the case of the pure design in Fig. S4a the increase in the SD of $a_g$ and $a_f$ is roughly the same and is largely independent of the actual size of the effect (*i.e.* simply introducing residual contributions into the model, even though in reality they may be zero, is sufficient to substantially increase the SDs). In the case of the SD in $a_r$ a slightly different pattern emerges, in which the influence of the residual size does markedly change the reduction in precision.

For the mixed design in Fig. S4b the overall features are much the same, except in comparison with the pure design the SD in the susceptibility SNP effect $a_g$ is slightly smaller and the infectivity SNP effect $a_f$ is slightly larger. Since the covariance matrix **Σ** is unknown for real diseases, the degree to which residual contributions reduce the precision with which SNP effects can be estimated is as yet an unanswered question.

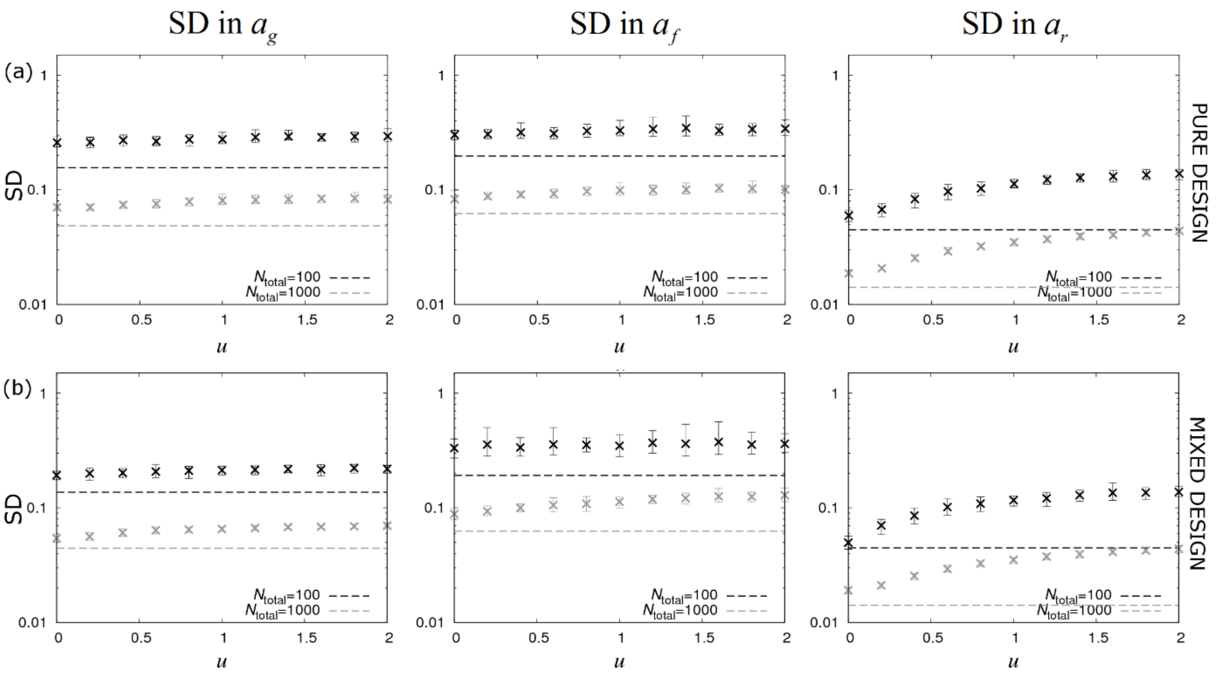


**Figure S4.** *Impact of residual contributions.* The left, middle and right columns show graphs for standard deviations in the SNP effects for susceptibility $a_g$, infectivity $a_f$ and recoverability $a_r$ as a function of the size of the residual contributions (as parameterized by $u$, where the covariance matrix is given by $\Sigma_{gg}=\Sigma_{ff}=\Sigma_{rr}=u$, $\Sigma_{gf}=0.3\times u$, $\Sigma_{gr}=-0.4\times u$, and $\Sigma_{fr}=-0.2\times u$) under optimised (a) pure and (b) mixed designs (without dominance). As residual contributions are increased, so the standard deviations increase above the analytical expressions. Dashed lines represent analytical results and crosses come from posterior estimates from simulated data (see Additional file 1). $N_{\text{total}}$ refers to the total number of individuals.
