## Additional file 12 for "Optimal experimental designs for estimating genetic and non-genetic effects underlying infectious disease transmission"

### Partitioning contributions to the SDs of SNP effects

Figure 6 in the paper shows the result of sequentially adding residuals, group effects and a fixed effect to the basic SNP model. This analysis assumed approximately 1000 individuals in the population. Figure S1 presents the corresponding results assuming only 100 individuals. Fewer individuals imply less statistical power which leads to higher bars in Fig. S1 compared to Fig. 6. In fact the factor of 10 difference in population size converts to a factor √10=3.2 difference in bar height. Note, however, that this only applies to the SNP, residual and fixed effect contributions. The contribution from the group effects is approximately the same in Figs. S1 and 6 (because there are the same number of contact groups in both cases).

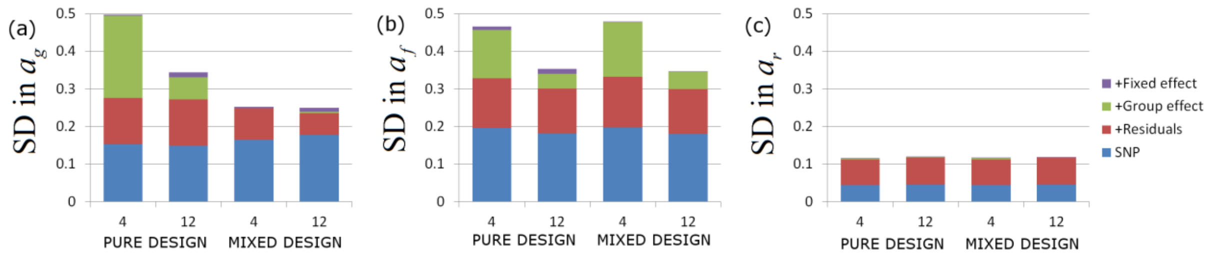


**Figure S1.** *Partitioning contributions to the SDs of SNP effects.* Residuals, group effects and a fixed effect are sequentially added to the basic SNP model. The corresponding increase in the SDs of the SNP effects is investigated for (a) the susceptibility $a_g$, (b) the infectivity $a_f$, and (c) the recoverability $a_r$. For comparison, four different scenarios are investigated: a pure design (without dominance) with respectively $N_{group}$=4 (*i.e.* a single version of the basic design) and $N_{group}$=12 (*i.e.* 3 replicates of the basic design) and a mixed design (without dominance) with $N_{group}$=4 (*i.e.* 2 replicates) and $N_{group}$=12 (*i.e.* 6 replicates). In each case approximately 100 individuals were partitioned equally amongst the contact groups. The residuals were chosen to have covariance matrix $\Sigma_{gg}=\Sigma_{ff}=\Sigma_{rr}=1$, $\Sigma_{gf}=0.3$, $\Sigma_{gr}=-0.4$, and $\Sigma_{fr}=-0.2$, the group effects had a SD of $\sigma_G=0.2$ and the fixed effect (assumed to represent sex with gender randomly allocated) had size $b_{g0}=b_{f0}=b_{r0}=0.2$. Results were found to be largely insensitive to these essentially arbitrary choices.
