## Additional file 13 for "Optimal experimental designs for estimating genetic and non-genetic effects underlying infectious disease transmission"

### Design replication

It was found in Additional file 11 that incorporating group effects substantially reduces the precision of SNP effects. This additional file investigates how this reduction can be moderated by means of design replication (that is repeating the same basic designs in Fig. 2 several times). Figure S1 shows that increasing the number of contact groups $N_{\text{group}}$ for a fixed total number of individuals (through replication of the basic design) leads to the SD in SNP effects reducing towards their analytical expectation.

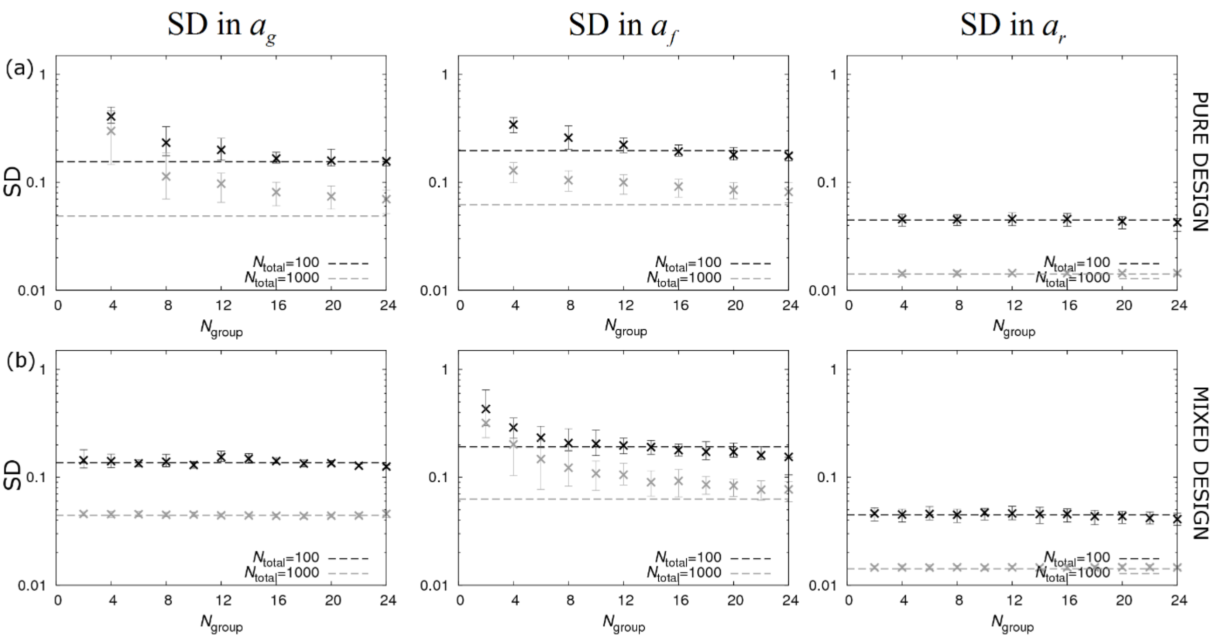


**Figure S1.** *Reducing the influence of group effects on precision of SNP effect estimates by increasing the number of groups through experimental replicates.* The left, middle and right columns show graphs for standard deviations in the SNP effects for susceptibility $a_g$, infectivity $a_f$ and recoverability $a_r$ as a function of the number of contact groups (with fixed number of individuals) for a model containing a group effect under optimised (a) pure design without dominance (with 4 contact groups per replicate, see Fig. 2b) and (b) mixed design without dominance (with two contact groups per replicate, see Fig. 2c). Dashed lines represent analytical results and crosses come from posterior estimates from simulated data (see Additional file 1). $N_{\text{total}}$ refers to the total number of individuals.
