## Additional file 14 for "Optimal experimental designs for estimating genetic and non-genetic effects underlying infectious disease transmission"

### Fixed effect correlated with SNP

This additional file investigates adding a single large fixed effect (with size $b_{g0}=b_{f0}=b_{r0}=0.2$) in Eq.(2), with elements in the design matrix $\mathbf{X}$ (which is now a vector because only a single fixed effect is considered) set to values 0 or 1 with equal probability in such a way as to give a certain degree of correlation with the SNP. This correlation is defined by

$$Corr(SNP, X) = \frac{\langle \boldsymbol{X}, \boldsymbol{W} \rangle}{\sigma_X \sigma_W}, \quad \text{(A1)}$$

where $\mathbf{W}$ is a vector giving the number of *A* alleles for each individual in the population.

Figure S1 shows results for the pure and mixed designs (without dominance). We find that provided there is little or no correlation between the fixed effect and SNP under investigation, the precision with which parameters can be estimated is largely unaffected. However, if there is significant correlation (especially in the case of inferring $a_f$) this can result in a significant loss in statistical power due to confounding. Therefore, when designing experiments, care should be taken to randomise (across groups) all factors other than the genotype of the SNP under investigation.

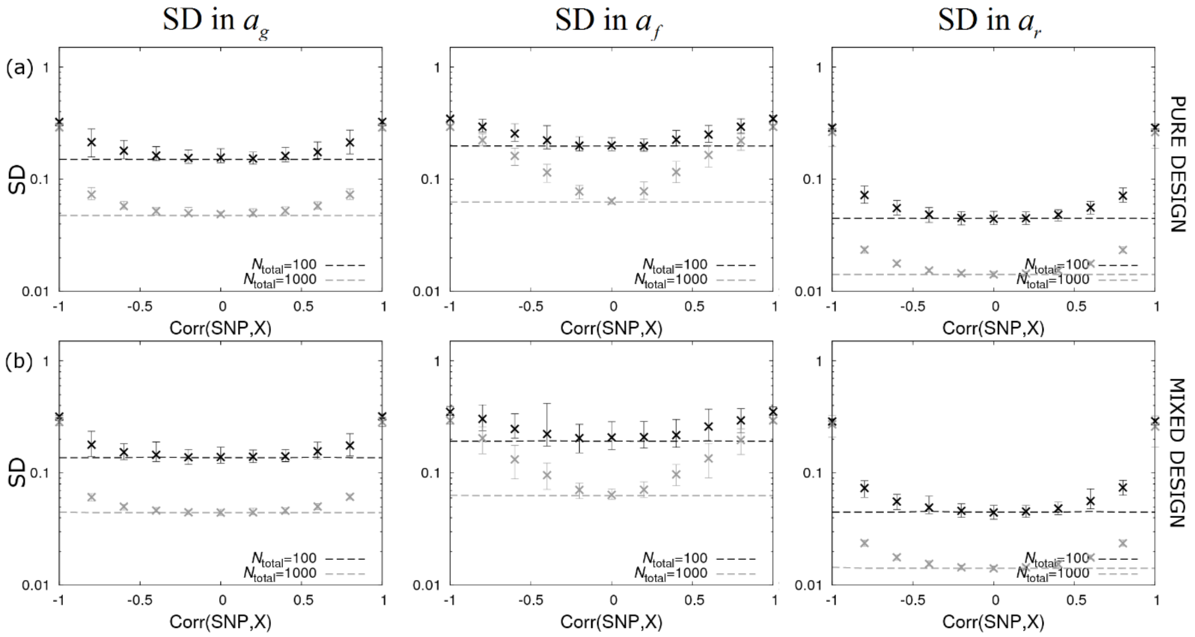


**Figure S1.** *Impact of a single fixed effect.* The left, middle and right columns show graphs for standard deviations in the SNP effects for susceptibility $a_g$, infectivity $a_f$ and recoverability $a_r$ as a function of the correlation between the number of *A* alleles at the SNP (which takes values 0, 1 or 2) and elements of the design matrix $\mathbf{X}$ for a single fixed effect under optimised (a) pure and (b) mixed designs (without dominance). The size of the fixed effect is taken to be $b_{g0}=b_{f0}=b_{r0}=0.2$. Dashed lines represent analytical results and crosses come from posterior estimates from simulated data (see Additional file 1). $N_{\text{total}}$ refers to the total number of individuals.
