## Additional file 15 for "Optimal experimental designs for estimating genetic and non-genetic effects underlying infectious disease transmission"

### Genotype determined by Hardy-Weinberg equilibrium

Rather than defining the proportions of genotypes in the seeder and contact populations, here we consider the case in which individual genotypes are randomly allocated with *A* allele frequency $p$. Assuming the SNP is in Hardy-Weinberg equilibrium [32] and that individuals are unrelated then for a population of size $N$ the following results can be derived:

$$\begin{aligned} \langle \chi \rangle &= 2p-1, \quad \mathrm{Var}(\chi) = \frac{2p(1-p)}{N}, \\ \langle H \rangle &= 1-2p(1-p), \quad \mathrm{Var}(H) = \frac{2p(1-p)\left(1-2p(1-p)\right)}{N}. \end{aligned} \tag{A1}$$

Applying these expression separately to the seeder and contact populations and substituting into the analytical expressions in Eqs.(6)-(8) leads to the curves shown in Fig. S1. Crucially we find that the precision of both the SNP effects and dominance parameters is maximised when the number of seeders $N_{\text{seed}}$ is small. In this limit the following results can be derived:

$$\begin{aligned} \text{SD in } a_g &\cong \frac{1}{\sqrt{2p(1-p)N_{\text{total}}}}, \\ \text{SD in } a_f &\cong \frac{1}{\sqrt{2p(1-p)\left(2-\frac{1}{G_{\text{size}}-1}\right)N_{\text{group}}}}, \\ \text{SD in } a_r &= \frac{1}{\sqrt{2p(1-p)kN_{\text{total}}}}, \end{aligned} \tag{A2}$$

and for the SDs in the dominance parameters:

$$\begin{aligned} \text{SD in } \Delta_g &\cong \frac{1}{|a_g|\sqrt{2p(1-p)\left(1-2p(1-p)\right)N_{\text{total}}}}, \\ \text{SD in } \Delta_f &\cong \frac{1}{|a_f|\sqrt{4p(1-p)\left(1-2p(1-p)\right)N_{\text{group}}}}, \\ \text{SD in } \Delta_r &= \frac{1}{|a_r|\sqrt{2p(1-p)\left(1-2p(1-p)\right)kN_{\text{total}}}}. \end{aligned} \tag{A3}$$

A key point to note is that the SD in SNP effect for infectivity $a_f$ in Eq.(A2) (and also for the SD in $\Delta_f$ in Eq.(A3)) now contains $N_{\text{group}}$ in the denominator instead of $N_{\text{total}}$. This means that increasing the number of individuals in contact groups does not substantially increase the precision with which $a_f$ can be estimated (a feature that was noted in [15]).

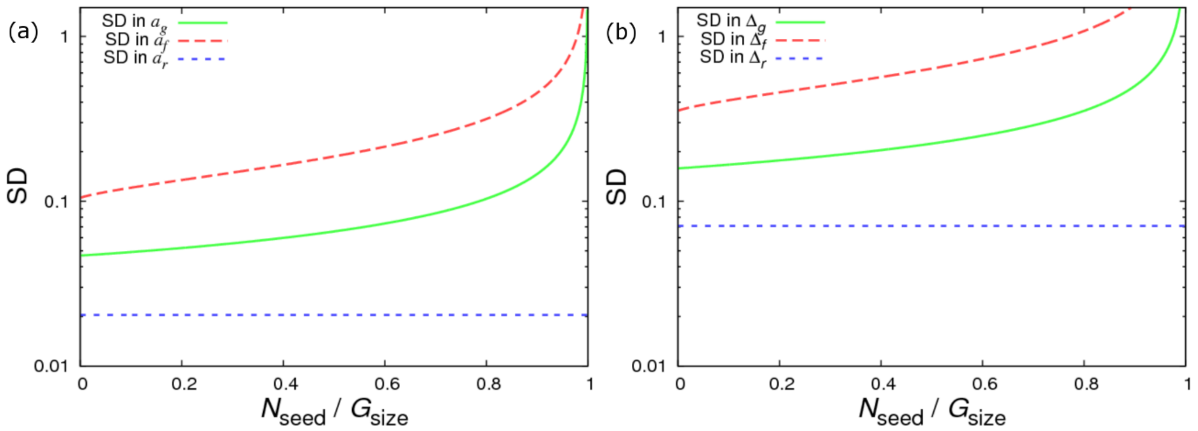


**Figure S1.** *Precision estimates for the SNP parameters with random genotype allocation.* These results come from the analytical expressions from Eqs.(6-13) (with Eqs.(A2) and (A3) representing the limit of small $N_{seed}$): (a) SNP effects and (b) dominance parameters. Individual genotypes are randomly sampled assuming Hardy-Weinberg equilibrium with mean *A* allele frequency $p$=0.4 (over all groups). Furthermore it is assumed that all individuals become infected (*i.e.* $\phi$=1) and parameters use were $k$=5, $G_{size}$=10 and $N_{group}$=100.
